# Intraspecific chemical variation of *Tanacetum vulgare* affects plant growth and reproductive traits in field plant communities

**DOI:** 10.1101/2023.03.27.534338

**Authors:** Lina Ojeda-Prieto, Pamela Medina-van Berkum, Sybille B. Unsicker, Robin Heinen, Wolfgang W. Weisser

**Affiliations:** Terrestrial Ecology Research Group, Department for Life Science Systems, TUM School of Life Sciences, Technical University of Munich, 85354 Freising, Germany; Department for Biochemistry, Max Planck Institute for Chemical Ecology, 07745 Jena, Germany; Plant-Environment-Interactions Group, Botanical Institute, University of Kiel, 24118 Kiel, Germany

**Keywords:** tansy, terpenoid, plant-plant competition, complementarity, functional traits, chemodiversity, volatile organic compound (VOC)

## Abstract

1. Intraspecific plant chemodiversity plays a fundamental role in interactions between plants and their interaction partners. However, how chemodiversity at the stand level (plant communities that vary in the number and type of plant chemotypes that grow in them, i.e., chemotype richness) affects ecosystem functioning is not fully understood.
2. We describe a biodiversity experiment using six chemotypes of common tansy (*Tanacetum vulgare* L., Asteraceae) to manipulate intraspecific plant chemodiversity at the plot level. We tested the effects of chemotype identity and plot-level chemotype richness (1-6) on plant growth and reproductive traits at plant and plot levels.
3. We found that chemotypes differed in growth and reproductive traits and that traits were affected by the plot-level chemotype richness. Although morphological differences became less pronounced over time, reproductive phenology patterns persisted. It suggests that chemotypes initially adopted different growth strategies, which may facilitate their establishment in nature.
4. Although chemotype richness did not lead to overyielding effects, plot-level trait means were affected by the presence or absence of certain chemotypes in a plot, and the direction of the effect depended on the chemotype.
5. We analyzed plot-level headspace emissions and found that blends released from plant communities were neither richer nor more diverse with increasing plot-level chemotype richness. However, we found that plots became more dissimilar in their headspace terpenoids as they were more dissimilar in their leaf-terpenoid profiles.
6. This long-term field experiment will allow further investigation into plant-insect interactions and insect community assembly in response to intraspecific chemodiversity.

## Introduction

Individuals of the same plant species can exhibit varying phenotypes, which reflect variation in growth, reproductive, physiological, and other traits (Raffard *et al.,* 2018; Siefert *et al.,* 2015; de Bello *et al.,* 2011; Fridley & Grime, 2010; Bolnick *et al.,* 2003). This variation within plant species is a major element in individual performance and population-, community- and ecosystem-scale processes (Westerband, Funk & Barton, 2021; Guisan *et al.,* 2019; Siefert *et al*., 2015; Violle *et al.,* 2012). Functional traits are attributes of species that can be measured at the individual level and are related to their response to environmental conditions and impact ecosystem properties and ecosystem functioning (Isbell *et al*., 2011). In addition to visible morphology-related functional traits, plants can vary intraspecifically in less apparent traits, such as chemical composition (Wetzel & Whitehead, 2020). For instance, many plant species show pronounced intraspecific variation in specialized metabolites along environmental gradients (Bakhtiari *et al*., 2019; Moore *et al*., 2014) or even at finer spatial scales, such as plant patches located in areas less than a few square kilometers (Kleine & Müller, 2011). Variation in chemodiversity has recently gained increased attention in ecology as it might structure community assembly or community composition as well as species interactions and fulfill ecosystem functions, including structuring plant-associated food webs and biodiversity (Erb & Kliebenstein, 2020; Wetzel & Whitehead, 2020; Müller *et al.,* 2020; Bálint *et al.,* 2016).

Individuals of a plant species can be clustered into distinct groups - also called chemotypes - by their dominant compounds, the composition of volatile and non-volatile compound blends, or by comparisons of specific specialized metabolites produced by individuals (Eilers, 2021). The consequences of specialized metabolites for intra- and interspecific interactions have been studied in various plant model systems (Moore *et al*., 2014), including *Tanacetum vulgare* (Neuhaus-Harr *et al.,* 2023; Eilers *et al.,* 2021; Clancy *et al.,* 2016; Kleine & Müller, 2011), *Jacobaea vulgaris* (Caralho *et al.,* 2014; Kostenko & Bezemer, 2013; Macel, 2011), *Plantago lanceolata* (Wurst *et al.,* 2008; Harvey *et al.,* 2005), *Senecio inaequidens* (Macel & Klinkhamer, 2010; Cano et al., 2009), and *Brassica oleracea* (Bustos-Segura *et al.,* 2017; Kos *et al.,* 2011; Kabouw *et al.,* 2010; Poelman *et al.,* 2009; Gols *et al.,* 2008). Specialized metabolites can be stored in specialized structures, such as glandular trichomes in *T. vulgare*, and released into the environment as volatiles. Consequently, some volatile organic compounds (VOCs) can be found in leaves and headspace. Conversely, some compounds are only produced de novo and released immediately, for instance, in plants without storage structures. These VOCs and compounds found in leaves shape the assemblage and interaction within the plant-associated insect community (Ponzio *et al*., 2013). Although ecologists are beginning to understand the consequences of plant chemodiversity for plant-insect interactions, much less is known about how it affects plant-plant interactions (Thorpe *et al.,* 2011).

Plants can display intraspecific chemodiversity at different levels. Due to variations in chemical composition at a small scale, groups of plants in a natural stand may differ in their chemotypes (Clancy, 2021; Eilers, 2021; Müeller *et al.,* 2020; Clancy *et al.,* 2018; Seft *et al.,* 2017). In such scenarios, intraspecific chemodiversity can be described by the richness (i.e., the number of chemotypes) and relative abundance of different chemotypes at the patch level. So far, the ecological consequences of variation in chemodiversity at the stand level have rarely been investigated.

The effects of increasing plant diversity on processes at the ecosystem level have been intensively studied in the framework of biodiversity-ecosystem functioning research (Weisser *et al.,* 2017; Chapin III *et al*., 1992). In most empirical work, plant diversity was manipulated by creating plant communities that differed in the number of plant species they contained. In contrast to the wealth of studies manipulating plant community diversity at the plant species level, there are far fewer studies where plant community diversity is manipulated at the within-species level, i.e., by creating plant communities with the same number of plant species, but with different extent of intraspecific variation. In a seminal study, Crutsinger *et al*. (2006) showed that in one-species *Solidago altissima* communities that differed in the number of *Solidago* genotypes they contained, increasing genotypic diversity enlarged arthropod community species richness and increased plant community biomass. The number of studies manipulating the intraspecific diversity of plant communities has been increasing in the past years (Raffard *et al*., 2019; Koricheva & Hayes, 2018; Genung *et al*., 2012). However, the detailed results differ between studies and taxa, even if one focuses on the effects on herbivores (Fernandez-Conradi *et al*., 2022; Hauri *et al*., 2022; Bustos-Segura *et al.,* 2017; Barton *et al*., 2015). A general trend is that increased intraspecific genotypic diversity of the plant community increases the diversity of the associated arthropod community and that these diversity effects on natural enemies of herbivores are generally higher than effects on the herbivores themselves, suggesting changes in top-down control (see meta-analysis in Koricheva *et al*., 2018; Hauri *et al*., 2021; Wetzel *et al.,* 2018). The mechanisms underlying these effects are still being discussed. In plots differing in genotypic diversity, effects on the herbivore community were found to be due to changes in plant size that varied both with genotype identity and with the particular genotypic composition of the plant community (Bustos-Segura *et al.,* 2017; Genung, 2012). In addition, there were effects of genotypic identity on the herbivore community that were not mediated by plant size (Bustos-Segura *et al.,* 2017; Genung, 2012).

Plants in intra-specifically diverse communities may compete with their neighboring conspecifics, resulting in changes in trait expression such as growth (Ziaja & Müller, 2022; Bustos-Segura *et al.,* 2017; Genung, 2012; Viola *et al*., 2010). On the other hand, plants may also experience reduced competition due to niche partitioning because of a priori differences in functional traits, such as phenology (Gallien, 2017; Kuppler *et al*., 2016). Little is known about how the chemotypes of plants differ in other traits and how intraspecific chemotype richness at the stand level affects individual plant performance.

Here, we designed a field experiment using *Tanacetum vulgare* L. (Asteraceae), in which we manipulated plot-level chemotype richness and the presence of particular plant chemotypes within the plots, to test the effects on plant growth and volatile emission. *T. vulgare* exhibits a high intraspecific variation in specialized metabolites (Bálint *et al*., 2016; Rohloff *et al.,* 2004), mainly mono- and sesquiterpenes (Ziaja & Müller, 2022; Keskitalo *et al*., 2001), and is easy to propagate clonally (Bálint *et al*., 2016). Moreover, intraspecific chemodiversity of *T. vulgare* occurs in different geographical regions and within populations, meaning that plant-plant interactions of distinct chemotypes often occur in close proximity in nature (Clancy *et al*., 2016; Kleine & Müller, 2011).

We measured growth and reproductive traits over two growing seasons and sampled headspace volatiles to test the following hypotheses:

1. At the individual plant level, chemotypes will differ in growth and reproductive traits, which will be affected by the plot-level chemical richness of the plots they grow in.
2. At the plot level, higher plot-level chemotype richness will increase plant growth, in line with the generally observed positive effects of biodiversity in the ecological literature.
3. Individual chemotypes will differ in their effect on plot-level effects.
4. Plant communities with higher plot-level chemotype richness will emit a more chemically diverse headspace of volatile organic compounds (VOCs) than plots with lower plot-level chemotype richness.

## Materials and methods

### Chemotypic characterization of <u>T. vulgare</u> lines and biological replicates

Plant chemotyping and chemotype selection are described in detail in Supplementary Methods and Neuhaus-Harr *et al*. (2023). The leaf terpenoid composition of the *T. vulgare* chemotypes used is shown in Fig. 1a and Table S1-1. We named the chemotypes by their dominant compounds as follows. The ’Athu-Bthu’ chemotype had both α- and β-thujone as prevalent compounds. The ’Bthu-high’ and ’Bthu-low’ chemotypes were dominated by β-thujone but had high or low relative levels of this terpenoid. There was a strong dominance of chrysanthenyl acetate in the ’Chrys-acet’ chemotype. The ’Mixed-high’ chemotype featured three terpenoids with a high relative total concentration, whereas the ’Mixed-low’ chemotype featured several terpenoids evenly contributing to the total profile.

**Figure 1:**
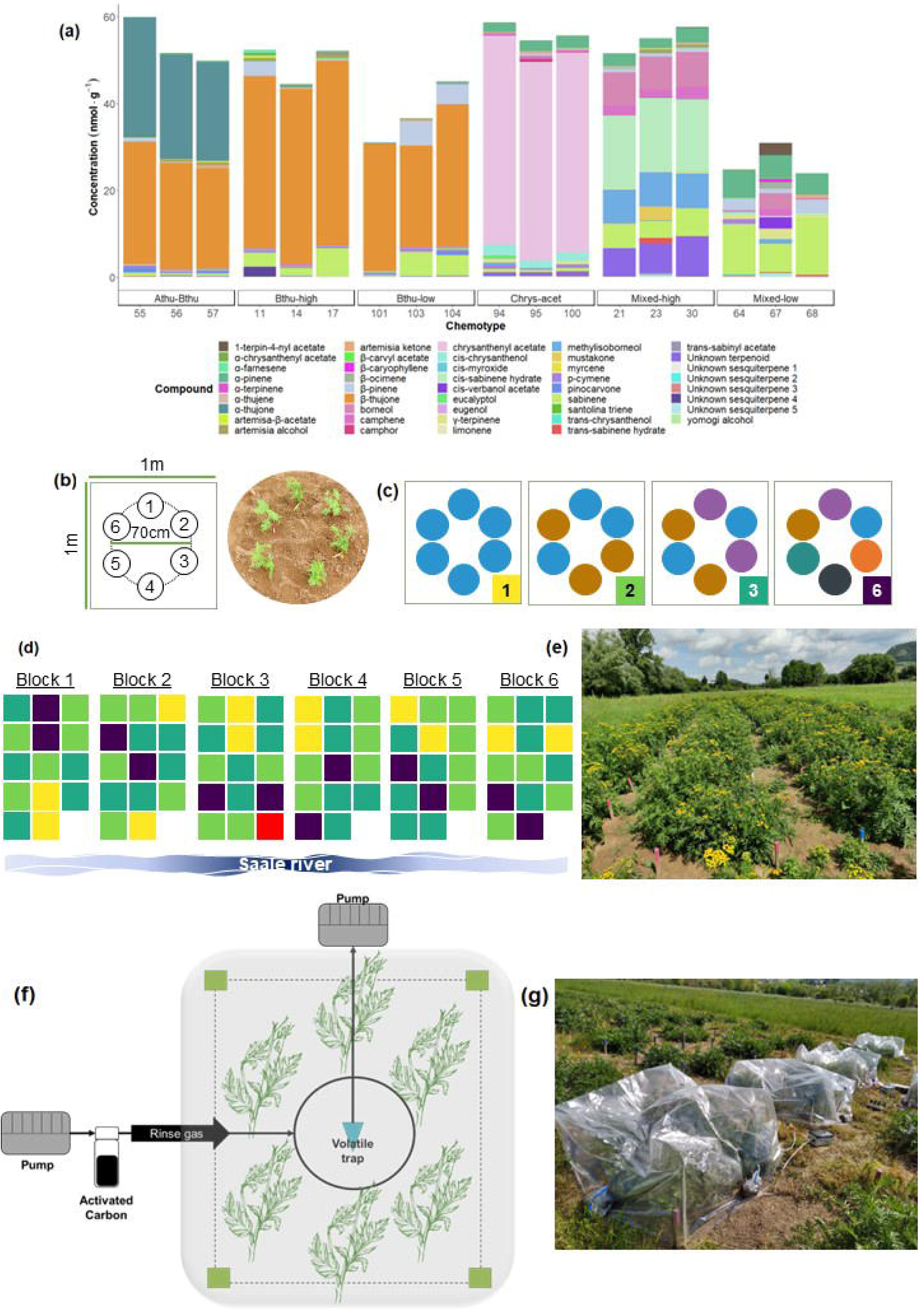
Field experimental design. **(a)** Stacked bars modified from Neuhaus-Harr *et al*. (2023) show approximate concentrations of terpenoid compounds (nmol g^-1^) extracted from leaf samples of the 18 selected daughters (exact values are reported in Table S1-1). Daughter replicates are clustered by their chemotype. **(b)** Plant arrangement within the plot: six *T. vulgare* plants were evenly distributed around a 70 cm diameter circle. The identity of tansy chemotypes in each plot was assigned to plots *a priori*, and plot position was assigned randomly. **(c)** Plot-level chemotype richness: The number of different chemotypes in one plot ranged from 1, 2, 3, to 6. **(d)** Block design showing plot-level chemotype richness 1, 2, 3, and 6 by yellow, light green, dark green, and purple squares, respectively. N = 84 plots were distributed equally in six randomized blocks. Each block consisted of 14 plots: two plots of chemotype richness level 1, five plots of chemotype richness level 2, five plots of chemotype richness level 3, and two plots with chemotype richness level 6. The red plot indicates the location of the background volatile profile plot used only for VOC analysis. **(e)** Picture of the field in June 2021. **(f)** Diagram of the closed-push-pull headspace VOC collection system. Purified air enters the collection PET bag and is pulled through a volatile trap located in the middle of the 6 plants at the higher side of the bag. **(g)** VOC collection setup in May 2022.

### Propagation of plant material for the field experiment

Plants were propagated via stem cuttings. A detailed description can be found in the Supplementary Methods.

### Experimental design

The field experiment took place in the Jena Experiment site located on the floodplain of the Saale River in Jena, Germany (50°55’N, 11°35’E, 130 m.a.s.l., Weisser *et al*., 2017). In May 2021, all vegetation was removed, and the soil was mechanically tilled. The field was divided into 84 plots (1 x 1 m) separated from each other by 70 cm footpaths and distributed in 6 replicated blocks (Fig. 1d). No fertilizer and no chemical insecticides or fungicides were used, and weeds were removed by hand every two weeks during the growing season.

We created plots of six tansy plants that were evenly distributed around a 70 cm diameter circle (84 plots x 6 plants = 504 plants total; Fig. 1b). The design followed suggestions of the design for biodiversity experiments: each plot differed in the number and identity of tansy chemotypes assigned to them. The number of different chemotypes in one plot (hereafter: plot-level chemotype richness) ranged from 1, 2, 3, to 6 (Fig. 1c). For instance, a plot with a plot-level chemotype richness of 1 was assigned six plants of the same chemotype. Within the plot, we maximized the number of daughters per chemotype. Daughters from each chemotype were equally distributed over the different treatment plots where possible or structurally assigned where an equal division was not possible (e.g., in the case of a shortage of plants of one daughter, they were replaced with a randomly picked daughter of the remaining two daughters of the same chemotype). So, a plot with plot-level chemotype richness 1 was assigned two clones from each of the three daughters of the plot chemotype; for plot-level chemotype richness 2 there was one clonal plant of each of the three daughters of each of the two chemotypes in the plot, etc. Lastly, a plot with plot-level chemotype richness 6 was assigned one clonal plant from one daughter of each of the six different chemotypes.

Each plot’s plot-level chemotype richness composition was *a priori* assigned - before in-depth analysis of the chemotype terpenoid composition - to avoid biases favoring the presence of one chemotype over another. Each unique chemotype combination was thus replicated twice, except for plot-level chemotype richness level 6, which was replicated twelve times but with different daughters. Hence, our field experiment contained 12, 30, 30, and 12 replicate plots for plot-level chemotype richness levels 1, 2, 3, and 6, respectively. Plots were distributed equally in six randomized blocks. Each block consisted of 14 plots: two plots of chemotype richness level 1, five plots of chemotype richness level 2 and 3, and two plots with chemotype richness level 6 (Fig. 1d-e). The exact assignment of plants to plots is given in Table S1-2.

### Plant morphological trait measurements

We measured six morphological plant traits for individual plants, including four growth traits (the number of stems per plant, the height of the highest stem, and above-ground fresh and dry weight) and two reproductive traits (the cumulative number of flower heads and the flowering index). A more detailed description of each variable and time-points used are found in Supplementary Methods.

### Headspace VOC collection

Headspace volatile organic compound (VOC) emissions were collected at the plot level. A detailed description of the process is available in the Supplementary Methods.

### Statistical analysis

All analyses were performed in R version 4.2.2 and RStudio 2022.07.2+576 (R Core Team, 2021). A description of all models are found in Supplementary Methods.

To analyze plot-level chemotype richness, we calculated plot-level diversity metrics using the ‘*chemodiv*’ package (Petrén *et al*., 2023). Note that we calculated theoretical chemotype diversity metrics based on the cumulative terpenoid profiles of each respected daughter present in a plot and calculated realized volatile diversity metrics based on our headspace VOC collection at the plot level. Data were visualized using the ‘*ggplot2’*, *‘grid’, ‘gridExtra’*, and *‘ggpubr’* packages (Wickham, 2016; R Core Team, 2021; Auguie, 2017; Kassambra, 2020).

We distinguished between plant-level and plot-level analyses, whereby plant-level analyses focused on the performance of the individual plants growing in the different diversity plots, controlling for the plot they grow in. Plot-level analyses focused on plot-level measures of plant performance calculated by averaging over all plants in a plot. All analyses of the effects of chemodiversity on plant traits were done using mixed-effect models using the ‘*lme4*’ package (Bates *et al*., 2015). All variables were analyzed separately for each time point. For evaluating the effects on traits of individual plants, p-values were estimated by type II Wald-Chi-Squared tests using the *Anova()* function in the ‘*car’* package (Fox & Weisberg, 2019) and by type I Wald-Chi-Squared tests using the *anova()* function in base R for evaluating the effects of chemotype presence on plot-level measurements and overyielding indices. After fitting a model, *post hoc* pair-wise comparisons among factor levels (i.e., chemotypes) were assessed with the ‘*emmeans’* package with Tukey adjustment (Russell, 2021).

### Effects of chemotype and plot-level chemotype richness on traits of individual plants

To test the effects of chemotype and plot-level chemotype richness on traits of individual plants growing in the different plots with six plants each, both factors (chemotype, plot-level chemotype richness) and their interaction were included as fixed factors in Generalized Linear Mixed-Effect Models (GLMM), with Poisson distribution for counting data (number of stems for both years and the cumulative number of flower heads), and with Linear Mixed-Effect Models (LMM) for the other individual plant traits (plant height, flowering index, and square root transformed above-ground dry weight and square root transformed above-ground fresh weight). We treated chemotype richness as a continuous variable with one degree of freedom. In the main analysis, random effects were Daughter ID (to account for technical replication of each daughter via cuttings) and Plot ID (84 in total) nested in Block ID (R scripts are provided as S3). In some variables, the random effect structure of the model led to singularity. We performed a second model excluding Plot ID since it had the lowest explanatory power of all random effects. We then chose the higher quality statistical model using the estimator of prediction error AIC.

If the main effect of a factor was significant, *post hoc* pair-wise comparisons with Tukey adjustment were conducted to assess significant differences between factor levels. While this is the most correct analysis from the point of daughter distribution within chemotypes, it results in a low degree of freedom. Hence, we carried out additional analyses to complement our understanding of within- and between-chemotype differences.

We tested separate analyses to specifically test for variation between Daughter IDs within and between chemotypes and directly compared the daughter’s performance, which can be found in more detail in Supplementary Methods.

### Effects of chemotype and plot-level chemotype richness on plot-level means

We first averaged each plant trait at the plot level to test the effects of chemotype and chemotype richness on plot-level plant traits. To test the effect of the presence/absence of a chemotype on plot-level trait values, we carried out a separate analysis per chemotype. This was done by running a linear model with Block ID, chemotype presence (indicating whether a specific chemotype is present in the plot), plot-level chemotype richness, and the interaction between chemotype presence and plot-level chemotype richness, separately for each chemotype.

### Overyielding calculations

We assessed the plot-level performance (i.e., the plot-level mean yield of the measured trait) of plants growing in plots of different chemotype richness levels (plot-level chemotype richness = 2, 3, and 6) and compared this to their performance in a monoculture (i.e., the respective chemotype-specific plots in plot-level chemotype richness = 1). This was done by calculating the overyielding index (OI) according to Hector et al. (2002). Overyielding indices are positive when the yield for a given chemotype in a mixture is greater than expectations from monocultures and indicates overyielding (Loreau, 1998). Overyielding indices for each plot-level chemotype richness (*i* = 1, 2, 3, or 6) were obtained for above-ground dry weight, above-ground fresh weight, the cumulative number of flower heads, and the flowering index for each plot. Mathematically, we calculated OI*_i_* = (*Oc_i_* – *Ec*) x *Ec*^-1^, where *Oc_i_* is the observed yield of a mixture plot I obtained by means of yields of the plants in the mixture, and *Ec* is the expected yield for the plot. *Ec* was calculated by averaging the yield of monoculture of each of the chemotypes present in the plot (Table S2-11).

In separate analyses per trait, we carried out LMM models with the calculated overyielding indices for all plot-level chemotype richness levels, with plot-level chemotype richness as a fixed factor and Block ID as a random factor.

### Plot-level theoretical leaf and plot-level realized volatile chemodiversity metrics

We calculated chemodiversity metrics at the plot level based on the leaf terpenoid profiles of individual chemotypes before planting (plot-level *theoretical* leaf chemodiversity) and on the headspace VOC measurements in the field (plot-level *realized* volatile chemodiversity). Plot-level theoretical leaf chemodiversity metrics were calculated by summing up the absolute leaf terpenoid concentrations (nmol x g^-1^) produced by each of the six specific daughters present in each plot (Fig. 1a). Those absolute leaf terpenoid profiles were obtained from the chemical analysis performed on leaves from greenhouse plants in 2020 before making the cuttings and planting them in the field. Based on the individual values of each plant present in a plot, we calculated plot-level theoretical leaf terpenoid richness, concentration, Hill Shannon index, and Hill evenness (Petrén *et al.,* 2022). Plot-level realized volatile chemodiversity metrics were calculated based on the plot-level absolute terpenoid emissions (ng x h^-1^) detected by headspace VOC collection.

As plants were chemotyped *a priori* (in Neuhaus-Harr *et al*., 2023) before transplanting them to the field and chemotyping was hence not affected by field conditions, plot-level theoretical leaf chemodiversity was analyzed by linear models with plot-level chemotype richness as the independent variable, without putting block as a random factor into the model. Plot-level realized volatile chemodiversity metrics based on field-collected headspace VOCs were analyzed by linear models with plot-level chemotype richness as the independent variable and Block ID (6 blocks) and Collection Day ID (3 days) as random factors.

We also analyzed the correlation between leaf and headspace terpenoid profiles by calculating Bray-Curtis dissimilarity matrixes for hierarchical analyses and performing a Mantel test by specifying 9999 permutations using the *‘vegan’* package (Oksanen *et al*., 2022).

## Results

### Field establishment

The establishment of the field experiment was successful; all 504 plants survived until the first seasonal harvest date (October 28, 2021). By early May of the second year (May 2, 2022), 96.6% of the plants (487 of 504 plants) showed shoot regrowth. Of these 487 plants, 4 more plants, which had shown some above-ground growth, naturally died and were not present at the harvest date of the second year (October 05, 2022). Missing data was excluded from analyses of plant traits at the plant level, with 17 plants missing for the first (May), 20 missing for the second (July), and 21 missing for the final measurements (October). Due to regrowth and mortality, the missing plants between time points only partially overlap. Plot-level averages were calculated according to the number of plants registered in the plot at the measurement date, but we kept the original plot chemotype richness level for the statistical analysis as this was the treatment variable.

### Effects of chemotype and plot-level chemotype richness on traits of individual plants

We used analyses separated for each time point to assess the effect of chemotype and plot-level chemotype richness across and between seasons.

The number of stems of a plant was significantly affected by plant chemotype identity on June 01 (χ^2^_5_ = 24.99, P <0.001, Fig. S2-1) and June 22, 2021 (χ^2^_5_ = 21.49, P <0.001, Fig. 2a), but not on the other time points (P > 0.05, Table S2-2). *Post hoc* analyses revealed that in the 2021 season, the Mixed-high chemotype had the highest number of stems compared with all other five chemotypes. Thus, the effects of chemotype on the number of stems across chemotypes were not stable across nor between growing seasons (Fig. S2-1, Table S2-2). Differences in the number of stems across chemotypes were less pronounced towards the end of the first season (Fig. S2-1a-c) and became even indistinct for 2022 (χ^2^_5_ = 3.28, P = 0.657, Fig. 2d).

**Figure 2:**
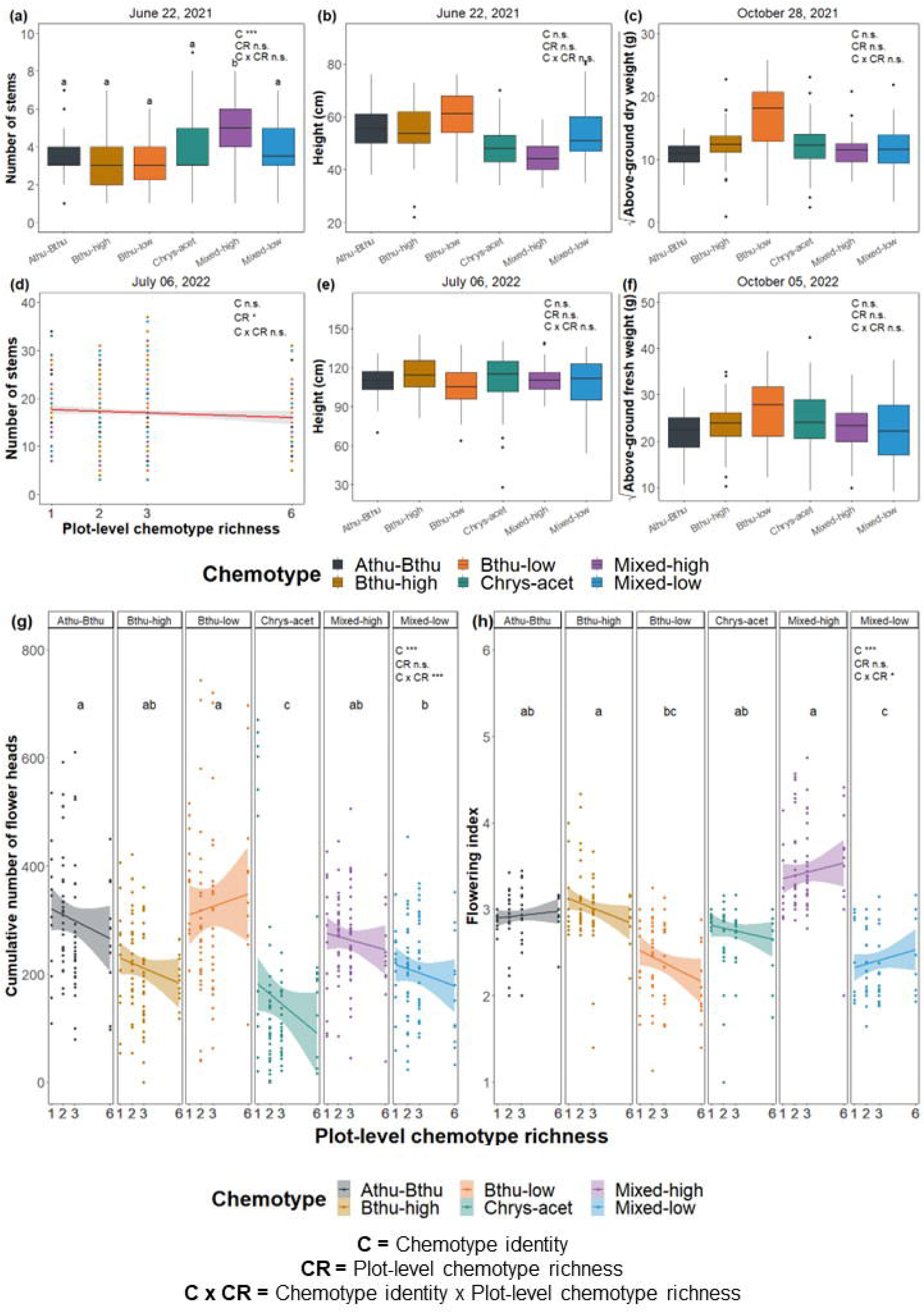
Effect of chemotype identity (C), plot-level chemotype richness (CR), and the interaction between chemotype identity and plot-level chemotype richness (C x CR) on growth traits of individual plants of *T. vulgare.* **(a)** Number of stems 2021, **(b)** height (cm) 2021, **(c)** square root above-ground dry weight (g) 2021, **(d)** number of stems 2022, **(e)** height (cm) 2022, **(f)** square root above-ground fresh weight (g) 2022, **(g)** cumulative number of flower heads 2021, and **(h)** flowering index in 2022. Significance is indicated as follows: n.s. = not significant, * P <0.05, ** P <0.01 and *** P <0.001. Degrees of freedom, Wald’s Chi-square statistics, and p-values are reported in Tables S2-2 - S2-5. Tukey *post hoc* significant differences between chemotypes are indicated with different letters.

For plant height, chemotypes were marginally different on June 22, 2021 (χ^2^_5_ = 10.54, P = 0.061, Fig. 2b) and strongly different when harvesting on October 28, 2021 (χ^2^_5_ = 12.26, P = 0.031, Fig. S2-2c). At this time point, several outliers caused deviation from a normal distribution that could not be solved with transformation. Therefore, for this time point, the chemotype effect must be interpreted cautiously. For instance, even though the chemotype had a significant effect, a *post hoc* test for October 28, 2021, did not reveal differences in height across chemotypes. Moreover, no effect of chemotype on height was kept in 2022 (Fig. 2e, Fig. S2d-f, Table S2-3).

For plant biomass, the above-ground weight of the Bthu-low chemotype was slightly higher in both growing seasons, although the effect of the chemotype was not significant (above-ground dry weight 2021: χ^2^_5_ = 9.35, P = 0.096, Fig. 2c; above-ground fresh weight 2022: χ^2^_5_ = 2.65, P = 0.753, Fig. 2f, Table S2-4).

There was no significant effect of plot-level chemotype richness nor the interaction between chemotype and plot-level chemotype richness for any of the growth traits measured on individual plants, except for an effect of plot-level chemotype richness on the number of stems in 2022 (χ^2^_5_ = 4.67, P = 0.031, Fig. 2d).

Both reproductive plant traits were significantly affected by plant chemotype: the cumulative number of flower heads (χ^2^_5_ = 55.08, P <0.001) and the flowering phenology (χ^2^_5_ = 51.07, P <0.001). Interestingly, reproductive plant traits depended on both chemotype and the plot-level chemotype richness, as indicated by an interaction between the two factors (cumulative number of flower heads: χ^2^_5_ = 779.33, P <0.001; flowering index: χ^2^_5_ = 14.93, P = 0.011). For example, the Bthu-low chemotype produced more flower heads in plots with higher plot-level chemotype richness in the first growing season. The other five chemotypes produced fewer flower heads in plots with higher plot-level chemotype richness (Fig. 2g). The Mixed-high, Mixed-low, and Athu-Bthu chemotypes showed more advanced flowering phenology (i.e., higher flowering index) in plots with higher plot-level chemotype richness. In contrast, the Bthu-low, Chrys-acet, and Bthu-high chemotypes showed more advanced phenology in plots with lower plot-level chemotype richness (Fig. 2h).

We found strong variation among daughters – across all chemotypes and within individual chemotypes –. More detailed results of the effect of daughter and plot-level chemotype richness on plant traits can be found in Supplementary Results.

### Chemotype presence and plot-level chemotype richness effects on plot-level measurements

The presence or absence of certain chemotypes in a plot affected plot-level variables related to plant growth: number of stems, plant height, above-ground dry weight, and reproduction: the cumulative number of flower heads. However, the effects of particular chemotypes were variable and differed with time in the growing season. Effects on the number of stems, number of flower heads, and flowering phenology are described and presented below, and additional effects on plant height and biomass are described in the Supplementary Results and Figures S2-8 – S2-10.

With respect to the mean number of stems per plot, three chemotypes significantly affected these in 2021, i.e., Bthu-low, Mixed-high, and Athu-Bthu on July 01, and two in July 2022, i.e., Mixed-high and Bthu-high (Fig. 3a), but not in the other time-point in 2022. The number of stems at the plot level significantly increased when Mixed-high was present in both time points (July 01: F_1,75_ = 12.85, P <0.001; July 22: F_1,75_ = 10.35, P = 0.002, Fig. S2-7). The Mixed-high chemotype was also the one with the highest number of stems in 2021 for both dates. The presence of the Bthu-high chemotype lowered the number of stems at the plot level on June 22, 2021 (F_1,75_ = 8.10, P = 0.006, Fig. 3a). The effect of the presence/absence of the Mixed-high or Bthu-high chemotypes did not differ across chemotype richness levels, indicated by the absence of interactions (Table S2-7). In 2022, chemotype presence/absence patterns on the number of stems differed slightly. Although we did not find a significant main effect of the plot-level chemotype richness of any of the chemotypes on the number of stems, we did observe an interaction between plot-level chemotype richness and the presence of the Mixed-high chemotype (F_1,75_ = 4.29, P = 0.042, Fig. 3b). In plots where the Mixed-high chemotype was present, the number of stems did not differ, whereas, in its absence, the number of stems decreased sharply with increasing plot-level chemotype richness. Across all six chemotype-specific models and in all time points in both years, plot-level chemotype richness had marginally significant negative effects on the mean number of stems per plot.

**Figure 3:**
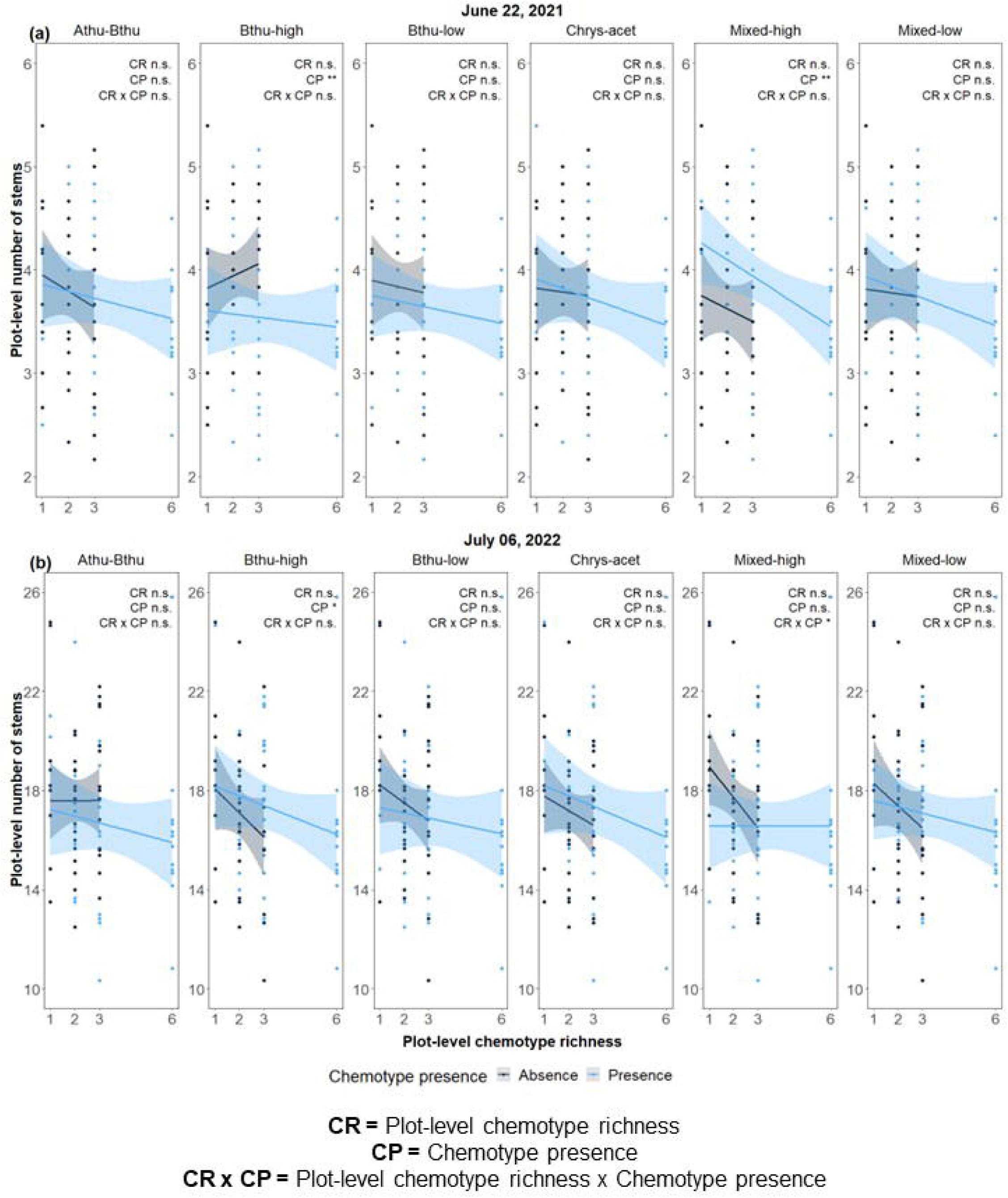
Effect of chemotype presence (CP), plot-level chemotype richness (CR), and the interaction between chemotype presence and plot-level chemotype richness (CP x CR) on average plot-level number of stems of *T. vulgare* in **(a)** 2021 and **(b)** 2022. Significance is indicated as follows: n.s. = not significant, * P <0.05, ** P <0.01 and *** P <0.001. Degrees of freedom, Wald’s Chi-square statistics, and p-values are reported in Table S2-7.

The number of flower heads per plot was affected positively by the presence of the Bthu-low chemotype in 2021 (F_1,75_ = 13.08, P <0.001, Fig. 4a), probably because the Bthu-low chemotype produced the highest number of flower heads among the chemotypes (Fig. 2g). In contrast, the presence of the Chrys-acet chemotype negatively affected the number of flower heads (F_1,75_ = 12.12, P <0.001). We also observed an interactive effect between the presence of the Bthu-low chemotype and plot-level chemotype richness (F_1,75_ = 5.25, P = 0.025), between the presence of the Mixed-high chemotype and plot-level chemotype richness (F_1,75_ = 4.98, P = 0.029), between the presence of the Bthu-high chemotype and plot-level chemotype richness (F_1,75_ = 7.08, P = 0.010), and between the presence of the Mixed-low chemotype and plot-level chemotype richness (F_1,75_ = 9.60, P = 0.003). In all the interactions, plot-level chemotype richness negatively affected flower number when the chemotypes were absent but not when they were present (Fig. 4a, Table S2-10).

**Figure 4:**
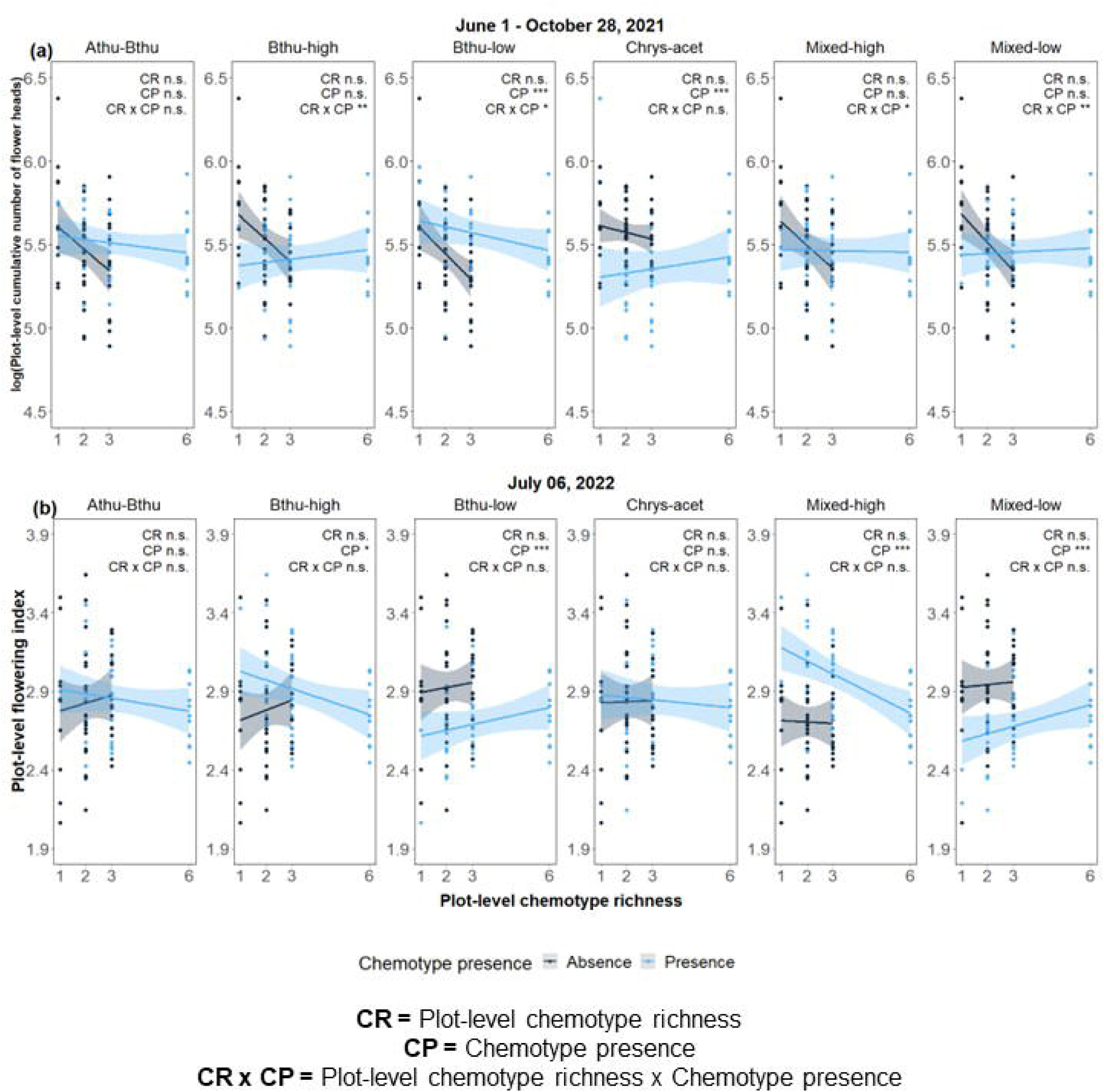
Effect of chemotype presence (CP), plot-level chemotype richness (CR), and the interaction between chemotype presence and plot-level chemotype richness (CP x CR) on reproductive traits of *T. vulgare* plants at plot level: **(a)** logarithm of the cumulative number of flower heads in 2021, and **(b)** flowering index in 2022. Significance is indicated as follows: n.s. = not significant, * P <0.05, ** P <0.01 and *** P <0.001. Degrees of freedom, F-statistics, and p-values are reported in Table S2-10.

For the second growing season (2022), the presence/absence of certain chemotypes also strongly affected the flowering phenology of plots. This was true for the Bthu-high, Bthu-low, Mixed-low, and Mixed-high chemotypes (Fig. 4b, Table S2-10). The presence of the Bthu-high or Mixed-high chemotype resulted in a higher flowering index, i.e., advancing flowering phenology compared to the plots where they were not present (Bthu-high chemotype presence: F_1,75_ = 5.88, P = 0.018; Mixed-high chemotype presence: F_1,75_ = 37.90, P <0.001). Conversely, plot-level flowering phenology was pulled down (i.e., less advanced flowering phenology) when the Bthu-low or Mixed-low chemotype was present than when it was not (Bthu-low chemotype presence: F_1,75_ = 16.27, P <0.001; Mixed-low chemotype presence: F_1,75_ = 22.42, P <0.001). All other models of the presence/absence of other chemotypes were non-significant.

### Overyielding calculations

We did not observe an overyielding effect in plot-level plant performance on plots with higher chemotype richness for any plot-level plant traits (plot-level chemotype richness always P >0.05, Table S2-12). However, we observed a weak negative tendency toward plants producing fewer flower heads (χ2 = 2.57, P = 0.109) when they were associated with more chemotypes (chemotype richness 2, 3, 6) compared to their performance in monocultures (Fig. S2-11).

### Headspace VOC analysis

The *T. vulgare* headspace VOC collections led to the identification of 60 compounds (Table S2-1). Classification of VOCs and comparison between plot-level headspace VOC profiles and plot-level chemotype richness are available in Supplementary Results.

### Plot-level theoretical leaf and plot-level realized volatile chemodiversity metrics

Our results confirmed that plots that had a higher plot-level chemotype richness (i.e., the number of different chemotypes present in the plot) also had higher *theoretical* leaf terpenoid richness, diversity, and evenness at the plot level (Fig. 5b-d, Table S2-14). However, the theoretical leaf terpenoid concentration did not show any relationship with plot-level chemotype richness (F_1,82_ = 0.00, R^2^ = 0.00, P = 0.967, Fig. 5a). For the terpenoids in the headspace VOC profiles, plot-level chemotype richness had no significant effect on the abundance of terpenes released as VOCs, or on terpene diversity (Fig 5e-5h, Table S2-15).

**Figure 5:**
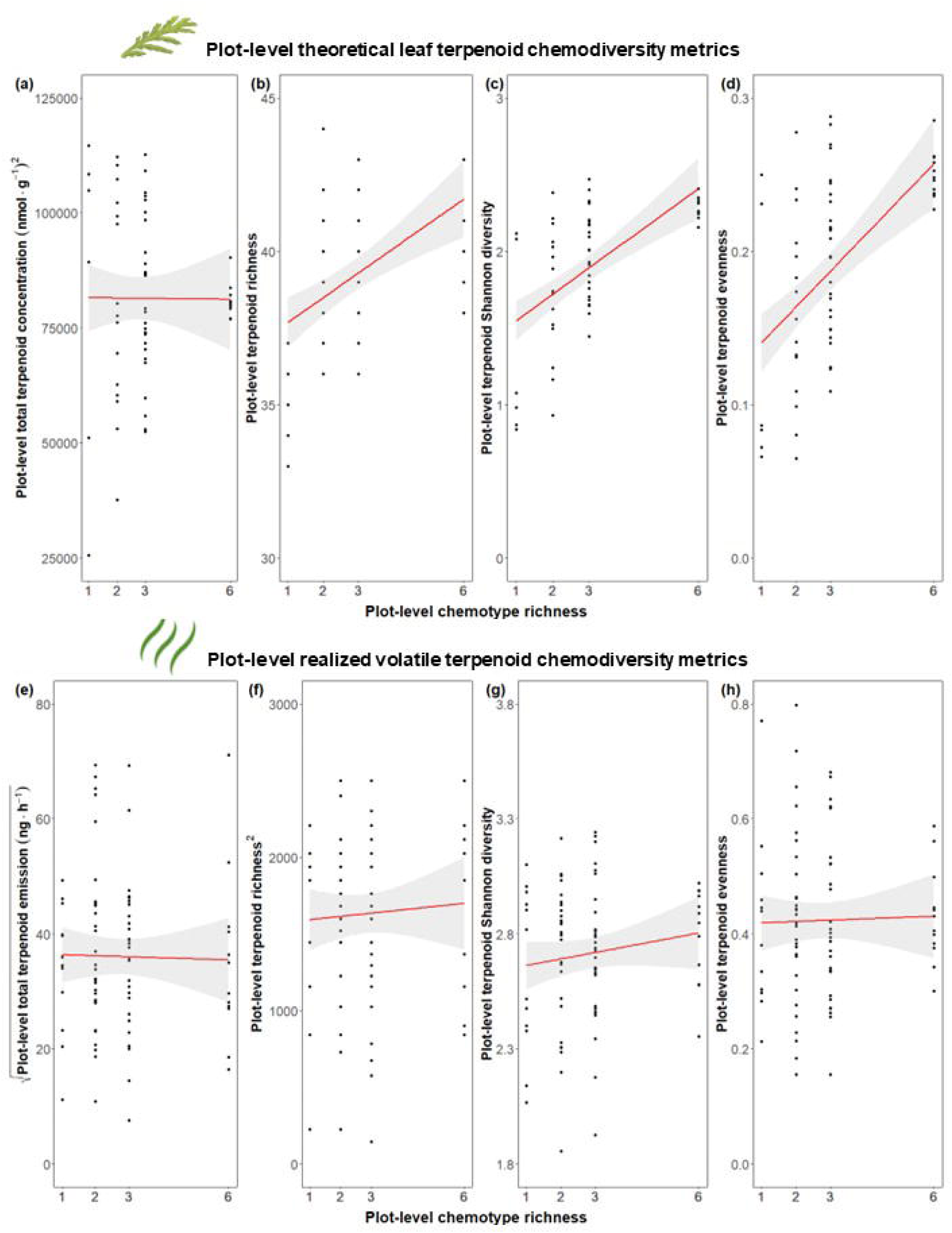
Effects of plot-level chemotype richness on theoretical plot-level chemodiversity metrics based on leaf terpenoid profiles (a-d) and realized plot-level volatile chemodiversity metrics (e-h). *Theoretical plot-level leaf* terpenoid diversity metrics were calculated by summing up the absolute leaf terpenoid concentrations (nmol x g^-1^) of each chemotype/daughter present in each plot based on the chemical analysis of leaves from greenhouse plants in 2020 (Neuhaus-Harr *et al*., 2023). Realized plot-level volatile chemodiversity was based on terpenoids collected in the headspace (ng x h^-1^) in May 2022. Diversity metrics were calculated using the ‘*chemodiv’* package (Pétren *et al*., 2022). Plot-level chemotype richness effects on *theoretical plot-level leaf* (a) squared total terpenoid concentration (nmol x g^-1^), **(b)** terpenoid richness, **(c)** terpenoid Shannon diversity, and **(d)** terpenoid evenness (summary of linear models in Table S5-13). Plot-level chemotype richness effects on *realized plot-level volatile* (e) squared root of total terpenoid emission (ng x h^-1^), **(f)** squared terpenoid richness, **(g)** terpenoid Shannon diversity, and **(h)** terpenoid evenness (summary of linear mixed-models in Table S2-14, S2-15).

Plots that were more diverse in terms of their leaf terpenoids were not necessarily more diverse in their plot-level headspace terpenoid profile. However, there was a weak positive correlation between leaf and headspace terpenoid profiles (Mantel statistic R^2^ = 0.05, P <0.001). As plots became more dissimilar in terms of their leaf terpenoids, they also became more dissimilar in terms of their headspace terpenoids.

## Discussion

Here, we designed a biodiversity field experiment in which we manipulated the number of chemotypes of *Tanacetum vulgare* plants at plot level to study how surrounding chemodiversity affects plant performance in the first two years after establishment. Our study showed that chemotypes initially differed in the studied morphological traits, confirming our first hypothesis. Concerning our second hypothesis, the effects of plot-level chemodiversity on plot-level traits were only found for plot-level averages of reproductive traits, but not for growth-related traits. The third hypothesis was also confirmed, i.e., the presence or absence of certain chemotypes and the plot-level chemotype richness influenced plot-level trait means. Importantly, the relationships between chemotypes, plot-level chemotype richness, and traits decreased over time in our 2-year study. Lastly, we found that the theoretical plot-level leaf terpenoid profiles significantly predicted the plot-level headspace terpenoid profiles, but the variance explained was low. Lastly, we observed relationships between chemodiversity and plant performance that changed over time.

Our results show that tansy chemotypes vary not only in their chemical profile but also in growth-related (number of stems and height) and reproductive traits (number of flower heads and flowering phenology) under field conditions. These findings broadly support the work of previous studies that used the same plant model system and found links between plant chemotypes and plant traits (Neuhaus-Harr *et al.,* 2023; Keskitalo *et al*., 2001). For instance, in a previous study, tansy chemotypes with a high concentration of camphor were taller compared to other chemotypes rich in trans-thujone, artemisia ketone, 1,8-cineole, or davadone-D, and chemotypes rich in artemisia ketone or davadone-D produced more flower heads and flowered later compared with the other four chemotypes (Keskitalo *et al*., 2001). However, since Keskitalo *et al*. (2001) analyzed the effect of chemical composition on certain traits at a broader geographic scale, and the chemotypes used by them and the ones used here are very different in their chemical composition, the current study’s findings are - although conceptually similar – hard to directly compare. In an earlier study using the same chemotype lineages used here, the chemotypes showed similar growth patterns (Neuhaus-Harr et al., 2023). This is not entirely surprising but reveals that clonally produced chemotypes show a high degree of consistency in phenotypes, at least in the early weeks of growth. Variability in growth traits across chemotypes may enable individuals to partition local resources into different growth strategies and avoid intraspecific competition (Gallien, 2017; Messier *et al*., 2010). For instance, the Bthu-low chemotype developed large biomass and typically few but tall and thick stems compared with other chemotypes such as Mixed-high, which developed more stems but shorter and thinner. Interestingly, however, the morphological differences between the chemotypes diminished over time. Only the Bthu-low chemotype remained quite distinct in both years, possibly due to its pronounced strategy of growing: tall and thick stems and high biomass. A likely explanation is the tendency for different daughters, even within chemotypes, to diverge in their trait expression over time, introducing enough variation to diminish chemotype-specific trait expression. This emphasizes the role of different growth strategies for individual survival, particularly during early establishment, and suggests that intraspecific chemodiversity might mediate niche realization processes (Müller & Junker, 2022).

Contrary to our hypothesis, no differences were observed in plant growth parameters when plants grew in plots that differed in plot-level chemotype richness, indicating that the morphology of each chemotype is consistent across the environments they grow in. Various possible explanations exist for why plant growth did not respond to plot-level chemotype richness. For instance, genotypes of *T. vulgare* can differ in their competitive ability in response to the presence of other plant species (Tse, 2014), but little is known about the response to the presence of conspecifics. Although intra- or interspecific competition is expected to affect plant growth, differences between *T. vulgare* plants might be strongly determined by the genotype, especially in the early growth stage, so intraspecific competition does not affect growth traits. It may very well be that growth responses to plot-level chemotype richness are not adaptive in *T. vulgare*, but as we will discuss below, responses occur in other traits.

The measured *T. vulgare* reproductive traits also pronouncedly differed between chemotypes, but in contrast to what we observed for growth traits, plant-level reproductive traits responded to plot-level chemotype richness. Although Moreira *et al*. (2016) and Hughes *et al*. (2008) suggested that plants growing in plots that contained more chemotypes express a higher individual plant fitness, in our study, all studied chemotypes, except for the Bthu-low chemotype, were inhibited (i.e., had lower flower head numbers) in plots with high plot-level chemotype richness compared to low-chemotype richness plots. It suggests that growth strategies that made the Bthu-low chemotype very dominant in height and weight in the first year might also bring some fitness advantages to the Bthu-low chemotype in highly diverse plant communities. It also suggests that most *T. vulgare* chemotypes may be negatively affected in more chemically diverse environments. This sharply contrasts with our expectations that chemically diverse environments would benefit plants. Our interpretation is based on the effect of chemodiversity on plant traits. Chemically diverse environments might still benefit plants by affecting plant interaction partners, as in Ziaja & Müller (2023), who reported that some *T. vulgare* chemotypes benefit from neighbors that differ in chemotype in terms of lower herbivore load of *Uroleucon tanaceti* and *Macrosiphoniella tanacetaria* aphids. However, the authors did not report any plant performance parameters, and therefore, a direct comparison between their and our studies investigating plot-level chemotype richness effects is currently impossible.

Another key finding was that plot-level chemotype richness influenced flowering phenology in *T. vulgare*. In the case of the Bthu-low chemotype, the effect of plot-level chemotype richness was positive on the number of flower heads but negative in the flowering index value, indicating a delayed onset of flowering. It appears that in different chemical environments, the flowering strategy differs across *T. vulgare* chemotypes. For instance, the Mixed-high, Athu-Bthu, and Mixed-low chemotypes had a more advanced flowering status at the time of assessment (high flowering index value) in plots with higher plot-level chemotype richness, while the Bthu-low, Chrys-acet, and Bthu-high chemotypes had a more delayed phenology (low flowering index value) in such plots. We speculate that *T. vulgare* may be able to sense their neighboring plants, either through direct competition (e.g., for space and nutrients), via the perception of volatiles (Ninkovic *et al*., 2021; Ninkovic *et al.,* 2019; Kessler & Kalske, 2018; Karban *et al*., 2014; Heil & Karban, 2010), or absorption of semi-volatile compounds emitted by neighboring plants (Himanen *et al.,* 2010). As a result, *T. vulgare* may avoid competition by being reproductively active at different times, thereby optimizing their fitness. Variations in flowering phenology may also correlate with variations in interacting arthropod communities, which may affect reproductive success (Kuppler *et al*., 2016). Moreover, differences in flowering phenology across chemotypes may constitute a strategy to avoid cross-pollination between certain tansy chemotypes that can result in poor seed production (Keskitalo *et al.,* 1998).

As predicted, plot-level average trait values were, in 2021, higher in mixture plots containing certain chemotypes that had either more stems, were higher, had larger above-ground dry weight, or produced more flower heads. In the same line, plot-level trait values decreased when chemotypes with fewer stems, smaller, lighter, or that produced fewer flower heads were present. Several types of interactions can be observed, ranging from adverse effects due to competition, possibly for a limiting resource, to positive effects through facilitation, for instance, through increased resource availability or a decrease in herbivory (Ziaja & Müller, 2023; Marquard *et al.,* 2009; Roscher *et al.,* 2005). In our study, the influence of a highly chemically diverse environment led to a higher performance of the Bthu-low chemotype (i.e., a higher cumulative number of flower heads). At the plot level, individual chemotype contributions appear to become less pronounced over time, suggesting that the plots become more similar as they age, at least in terms of their morphological structure. Furthermore, we found no overyielding effects in our system, but instead, we found a tendency toward lower plot-level plant trait values when increasing chemotype richness. This is important for future and ongoing work in this field experiment, which will investigate the role of chemodiversity on insect community assembly. A reduction of the plant- and plot-level differences in growth traits reduces the strength of potential confounding effects of plant growth on insect community assembly and will, therefore, help draw more robust conclusions and deepen our understanding of the consequences of chemodiversity. Moreover, chemotypes of tansy plants have been studied regionally in their range of distribution (Rahimova *et al.,* 2023; Clancy, 2021; Wolf *et al*., 2012), and little attention has been paid to the implications of naturally co-occurring neighboring chemotypes.

In line with our predictions, most diversity metrics based on theoretical plot-level leaf terpenoid profiles increased with increasing plot-level chemotype richness. We observed no differences between plot types for the total approximate terpenoid concentration. In contrast to what we hypothesized, we found no effect of plot-level chemotype richness on diversity metrics calculated based on headspace terpenoids. More diverse plots based on the theoretical leaf terpenoid of each chemotype were not necessarily more diverse in their plot-level headspace terpenoid profile. Headspace volatile terpenoid of a plant community is strongly dependent not only on the constitutive specialized metabolite profile of their individual plants but also on the intrinsic chemical properties of their compounds (e.g., volatility and solubility) and on physiological and morphological plant features (such as the presence of trichomes; see He *et al.,* 2011; Aschenbrenner *et al.,* 2013; Gershenzon & Dudareva, 2007), and biological characteristics such as the abundance and interacting organisms above- and below-ground, plant age, and plant size (Fabisch *et al*., 2019; Kessler & Kalske, 2018; McCormick *et al*., 2012; Takabayashi *et al*., 1994). Although volatile production is hard to accurately predict (Dicke *et al*., 2009; Dudareva *et al.,* 2006), it is plausible that the volatile headspace profile will be related to the compounds found and stored in glandular trichomes in leaves of *T. vulgare*, since this plant has the capacity of store VOCs. However, while the hexane extraction was performed from the leaves of young plants before planting them in the field in 2021, the headspace terpenoids were obtained in the spring season one year after planting (2022) under field conditions (where the plants interacted with each other and other organisms) and analyzed in a different lab. This could have led to changes in terpenoid diversity that may be reflected in our analysis (Eckert *et al.,* 2023). Moreover, emitted volatile profiles might also differ from the stored ones because some specific volatile organic compounds are produced only upon attack or abiotic stress (like green leaf volatiles, benzenoids, and terpenoids known as herbivore-induced plant volatile -HIPV-, Rashid & Chung, 2017; Unsicker *et al*., 2009). For example, Clancy and collaborators (2016) found that emitted *T. vulgare* headspace volatiles differed from compounds found in leaves at the individual level by 82%. However, the extent to which the chemotype richness and stored chemical profiles of a group of plants affect the volatiles in their communal headspace remains poorly understood. It will be interesting to see how the community-level volatile profile will develop over time when plants grow further and competition intensifies.

Further research should be undertaken to investigate the qualitative changes in plot-level volatile composition and how they correspond to plot-level cumulative leaf terpenoid profiles, but this was beyond the scope of this study. Future investigations of leaf terpenoids and headspace terpenoids under field conditions over time, within and across growing seasons, upon induction, and in response to environmental stresses would help us understand the temporal dynamics of plant terpenoid composition and volatile emission. Furthermore, research on the mechanistic understanding of different defensive strategies, such as the storage of defense metabolites and the emission of volatiles, might help to understand these divergent patterns we observed.

A mechanistic understanding of biodiversity-ecosystem functioning requires analyzing the role of not only species number and functional groups but also phylogenetic diversity and within-group variation in functional traits (Tilman *et al*., 1997b). Our results provide insights into the underlying processes through which intraspecific chemodiversity acts on plant growth and reproductive traits. Although this study focuses on plant traits, this established field experiment also raises the possibility of further studying the role of intraspecific chemodiversity in interactions between plants and associated interaction partners, such as herbivorous insects and their natural enemies and pollinators. Insect diversity and abundance are typically positively correlated with plant species diversity and interspecific diversity of functional traits (Junker *et al*., 2015). However, the effect of intraspecific plant chemodiversity on shaping such interactions in the field has received limited attention to date (but see Bustos-Segura *et al.,* 2017). The findings of this study suggest that intraspecific chemodiversity might influence ecosystem properties such as primary productivity, resource use efficiency, ecosystem stability, and resilience. Given the degradation of morphological trait differences over time, this field experiment offers a unique opportunity to study the effect of chemodiversity by means of constitutive plant terpenoid profiles in nature.

## Supporting information

Supplementary Results

## Acknowledgments

We thank the staff at TUM PTC Dürnast, particularly Sabine Zuber and Petra Scheuerer, for providing excellent care during the plant propagation and preparation phase. We thank Caroline Müller and Elisabeth Eilers for chemotyping the plants, and Anne Ebeling and Nico Eisenhauer for providing space within the Jena Experiment and allowing us to use core facilities under Gerlinde Kratzsch’s guidance. We thank Nafiseh Mahdavi for her help in establishing the field experiment, Leonardo Moreno for his help in the maintenance of the plants and in collecting data on both years of the field experiment, and the Jena Experiment gardening staff for providing advice and help in the maintenance of the experiment. Many thanks to Beate Piecha, Annika Neuhaus-Harr, and other friends and colleagues who helped during the harvests. This study was funded by the German Research Foundation (DFG), project 415496540 (WE3081/40-1), as part of the Research Unit (RU) FOR 3000. We thank all members of the RU for their valuable discussions.

## Author contributions

WWW conceived the original idea and designed the experiment. RH prepared the field experiment and propagation of plants. LOP, RH, and WWW planted the field experiment. LOP collected all data and maintained the field experiment. LOP analyzed ecological data with input from RH and WWW. PMvB, SBU, RH, and LOP prepared and executed volatile collections. PMvB chemically analyzed volatiles. LOP wrote the manuscript with input from RH and WWW. All authors contributed to the final version and approved the final submission.

## Data availability statement

Data is available on request.

