## Supplementary Results for "Intraspecific chemical variation of *Tanacetum vulgare* affects plant growth and reproductive traits in field plant communities"

**Supplementary Information 2 (S2): Supplementary Results**

**Table of content**

**2.1. Detailed description of results**

*2.1.1. Effects of daughter and plot-level chemotype richness on traits of individual plants.*

*2.1.2. Effects of chemotype and plot-level chemotype richness on plot-level means.*

*2.1.3. Headspace VOC analysis.*

**2.2. Compounds identified in plot headspace analysis**

**2.3. Details of the statistical analyses**

*2.3.1. Tables.*

2.3.1.1. Effects of chemotype and plot-level chemotype richness on traits of individual plants.

2.3.1.2. Effects of daughter and plot-level chemotype richness on traits of individual plants.

2.3.1.3. Effects of chemotype and plot-level chemotype richness on plot-level means.

2.3.1.4. Overyielding calculations.

2.3.1.5. Headspace VOC collection.

2.3.1.6. Plot-level theoretical leaf and plot-level realized volatile chemodiversity metrics.

*2.3.2. Figures.*

2.3.2.1. Effects of chemotype and plot-level chemotype richness on traits of individual plants.

2.3.2.2. Effects of daughter and plot-level chemotype richness on traits of individual plants.

2.3.2.3. Effects of chemotype and plot-level chemotype richness on plot-level means.

2.3.2.4. Overyielding indexes.

2.3.2.5. Headspace VOC collection.

### 2.1. Detailed description of results

#### *2.1.1. Effects of daughter and plot-level chemotype richness on traits of individual plants*

We found strong variation among daughters – across all chemotypes and even within individual chemotypes – in terms of the number of stems (Fig. S2-3), height (Fig. S2-4), and above-ground dry weight (Fig. S2-5) on October 28, 2021, and the cumulative number of flower heads in 2021 (Fig. S2-6, Table S2-6). Within chemotypes, only in the Athu-Bthu chemotype were no differences among daughters in above-ground dry weight ($\chi^2_2$ = 2.26, P =0.322) and flower count ($\chi^2_2$ = 5.28, P = 0.071). In the Mixed-high chemotype, we did not find differences between daughters in number of stems ($\chi^2_2$ = 4.01, P = 0.135) and height ($\chi^2_2$ = 50.04, P = 0.818).

#### *2.1.2. Effects of chemotype and plot-level chemotype richness on plot-level means*

Plant height in plots was increased in the presence of the Bthu-low chemotype in 2021 (June 22: $F_{1,75}$ = 14.41, P <0.001; October 28: $F_{1,75}$ = 24.00, P <0.001, Fig. S2-8) but not for 2022. Plants of the Bthu-low chemotype were comparatively high (Fig. 2b), which drove this effect. In contrast, the plot-level plant height in 2021 was always lower when the Mixed-high chemotype was present (Fig. S2-8, Table S2-8). The presence of the Chrys-acet chemotype also lowered the plot-level average height but only for June 22 ($F_{1,75}$ = 5.54, P = 0.021, Fig. S2-8b). The effect was stronger in low plot-level chemotype richness when present and high plot-level chemotype richness when absent, as indicated by a marginally significant interaction between plot-level chemotype richness and chemotype presence ($F_{1,75}$ = 3.80, P = 0.055). In 2022, only the presence of the Mixed-low chemotype lowered the mean plot-level height, but only in May 2022 (Fig. S2-9a). We observed no effects of plot-level chemotype richness on the mean plot-level height at any time point in any year (Fig. S2-8 – S2-9, Table S2-8).

In 2021, plot-level mean plant aboveground dry weight strongly increased when the Bthu-low chemotype was present in the plot ($F_{1,75}$ = 59.54, P <0.001, Fig. S2-10a). Similarly, but less pronounced, aboveground fresh weight increased when the Bthu-low chemotype was present in 2022 ($F_{1,75}$ = 10.70, P = 0.002, Fig. S2-10b). The presence of the Athu-Bthu chemotype reduced the mean above-ground dry weight at the plot level ($F_{1,75}$ = 9.89, P = 0.002), and it was greater than the effect found on the above-ground fresh weight for the year 2022 ($F_{1,75}$ = 7.24, P = 0.009). In both cases, the presence of the Athu-Bthu chemotype reduced mean above-ground weight (Table S2-9). All other models of individual chemotypes were non-significant.

*2.1.3. Headspace VOC analysis*

The *T. vulgare* headspace VOC collections led to the identification of 60 compounds (Supplementary Information S4). Volatile organic compounds were classified into four major classes: green leaf volatiles (GLV), monoterpenes, sesquiterpenes, alkanes, and other compounds that did not belong to these groups. The plot-level headspace VOC profiles did not vary when plot-level chemotype richness changed; for instance, plots that contained more chemotypes did not necessarily produce more diverse headspace VOC profiles. Monoterpenes and sesquiterpenes represented the most abundant and diverse groups of compounds in the headspace (84% of the compounds), followed by GLV with 15% (Fig. S2-12). Only terpenoids identified with the database were considered for comparison purposes.

While 44 terpenoids were detected by hexane extraction from leaf material on earlier chemotype characterizations before planting in the field, we found 49 terpenoids among the VOCs in the headspace of tansy communities once established in the field. Only terpenoids identified with the database were considered for comparison purposes; unknown terpenoids were excluded. It resulted in 28 terpenoids uniquely identified in the headspace of tansy communities (43.5%), 16 terpenoids solely identified on tansy leaves (24.6%), and 21 terpenoids identified by both methods (32.3%), i.e., headspace VOC collection and hexane leaf extraction (Table S2-13).

### 2.2. Compounds identified in plot headspace analysis

**Table S2-1:** Volatile organic compounds (VOCs) of *Tanacetum vulgare* at plot level collected by headspace push-pull system in the field. The table is partitioned into three sections, showing the group in order. Volatile compounds and their mean (+SE) concentration in ng/h in the different plot-level chemotype richness plots of *T. vulgare* (i.e., 1, 2, 3, 6)**.** Retention time, Kovats index, and classification of each compound are provided. VOCs were identified by comparing retention times and mass spectra with commercial standards (indicated with *) and by comparison to the mass spectral libraries Wiley 275 (Wiley and National Institute of Standards) and NIST05a (National Institute of Standards and Technology, Gaithersburg, MD). The VOC identity was verified by calculating Kovats retention indices. Gas chromatography with flame ionization (GC-FID) was used for the quantification of VOCs.

| Compound | Group | RT | KI | 1 | 2 | 3 | 6 |
|---|---|---|---|---|---|---|---|
| 2-hexanol | Fatty acids (GLV) | 4.45 | 825 | 205 ± 48.01 | 226.49 ± 27.93 | 208.04 ± 26.45 | 220.84 ± 47.55 |
| cis-3-hexen-1-ol* | Fatty acids (GLV) | 5.42 | 878 | 34.58 ± 6.43 | 35.95 ± 3.42 | 36.3 ± 3.44 | 38.34 ± 6.43 |
| cis-3-hexenyl acetate* | Fatty acids (GLV) | 9.01 | 1028 | 108.93 ± 27.39 | 91.23 ± 11.19 | 83.94 ± 11.02 | 75.54 ± 9.5 |
| heptanal | Fatty acids (GLV) | 6.21 | 916 | 7.44 ± 2.53 | 3.36 ± 0.54 | 2.45 ± 0.56 | 2.56 ± 0.98 |
| hexyl formate | Fatty acids (GLV) | 5.68 | 892 | 8.1 ± 1.82 | 9.27 ± 1.35 | 6.99 ± 0.82 | 5.89 ± 1.46 |
| trans-2-hexenal* | Fatty acids (GLV) | 5.31 | 872 | 4.58 ± 2.08 | 8.91 ± 1.82 | 8.12 ± 1.56 | 10.53 ± 4.63 |
| (-)-borneol* | Monoterpenes | 12.9 | 1183 | 9.54 ± 6.48 | 7.5 ± 2.91 | 8.59 ± 2.29 | 4.86 ± 2.24 |
| (-)-Terpinen-4-ol | Monoterpenes | 13.48 | 1207 | 21.63 ± 3.94 | 24.38 ± 3.14 | 23.94 ± 3.37 | 25.98 ± 8.45 |
| (+)-isothujol | Monoterpenes | 12.14 | 1152 | 10.95 ± 3 | 24.36 ± 7.3 | 15.45 ± 4.02 | 15.03 ± 5.12 |
| 2-carene | Monoterpenes | 10.99 | 1106 | 0.54 ± 0.37 | 0.65 ± 0.27 | 1.59 ± 0.65 | 2.11 ± 1.63 |
| bornyl acetate | Monoterpenes | 15.68 | 1302 | 7.55 ± 5.35 | 6.55 ± 1.98 | 5.45 ± 0.99 | 6.19 ± 2.17 |
| camphene | Monoterpenes | 7.43 | 965 | 26.92 ± 5.7 | 28.26 ± 4.17 | 21.8 ± 1.83 | 22.77 ± 2.58 |
| camphenol | Monoterpenes | 11.55 | 1129 | 3.2 ± 1.84 | 1.1 ± 0.44 | 1.16 ± 0.56 | 0.76 ± 0.52 |
| camphor* | Monoterpenes | 12.41 | 1163 | 53.65 ± 30.39 | 54.26 ± 21.83 | 32.95 ± 9.73 | 32.71 ± 8.89 |
| chrysanthenone | Monoterpenes | 11.93 | 1144 | 18.88 ± 12.23 | 11.94 ± 3.48 | 6.25 ± 1.55 | 5.76 ± 2.53 |
| cis-sabinol | Monoterpenes | 12.26 | 1158 | 4.76 ± 1.43 | 11.03 ± 3.9 | 9.21 ± 2.68 | 7.56 ± 2.25 |
| eucalyptol* | Monoterpenes | 9.61 | 1052 | 24.83 ± 11.82 | 37.41 ± 12.72 | 20.64 ± 3.19 | 26.33 ± 9.51 |
| limonene | Monoterpenes | 9.5 | 1047 | 47.23 ± 9.37 | 57.21 ± 7.11 | 51.61 ± 7.03 | 55.82 ± 17.74 |
| myroxide | Monoterpenes | 11.24 | 1116 | 18.54 ± 3.79 | 23.34 ± 4.1 | 88.17 ± 51.35 | 15.12 ± 3.62 |
| p-cimene* | Monoterpenes | 9.33 | 1040 | 54.42 ± 12.8 | 63.64 ± 9.48 | 48.35 ± 6.63 | 55.86 ± 18.74 |

**Table S2-1 *(continued)***: *This section shows the results for the second part of Table S2-1.*

| Compound | Group | RT | KI | 1 | 2 | 3 | 6 |
|---|---|---|---|---|---|---|---|
| pinocarvone | Monoterpenes | 12.64 | 1173 | 5.96 ± 0.94 | 7.7 ± 0.85 | 6.82 ± 0.73 | 6.77 ± 0.75 |
| sabinene hydrate* | Monoterpenes | 10.51 | 1087 | 96.76 ± 59.31 | 113.25 ± 52.5 | 55.03 ± 15.53 | 63.48 ± 20.32 |
| sabinene* | Monoterpenes | 8.13 | 993 | 98.58 ± 27.36 | 170.77 ± 32.63 | 152.09 ± 35.87 | 164.17 ± 48.95 |
| terpinen-4-ol | Monoterpenes | 13.16 | 1194 | 8.81 ± 2.64 | 9.1 ± 1.66 | 5.61 ± 0.73 | 6.17 ± 1 |
| terpinolene* | Monoterpenes | 11.14 | 1112 | 8.35 ± 2.85 | 5.54 ± 1.54 | 4.26 ± 0.67 | 3.79 ± 1.34 |
| thujone* | Monoterpenes | 11.38 | 1122 | 35.82 ± 17.71 | 18.9 ± 1.25 | 17.23 ± 1.22 | 16.19 ± 1.63 |
| trans-chrysanthenyl acetate | Monoterpenes | 14.51 | 1252 | 63.48 ± 44.32 | 20.32 ± 5.51 | 23.99 ± 7.74 | 24.63 ± 10.08 |
| trans-sabinene hydrate | Monoterpenes | 9.73 | 1056 | 3.52 ± 1.02 | 5.68 ± 0.82 | 5.57 ± 0.83 | 6.46 ± 2.02 |
| trans-β-ocimene* | Monoterpenes | 9.98 | 1066 | 137.97 ± 24.34 | 144.2 ± 23.11 | 139.01 ± 17.92 | 141 ± 41.24 |
| α -terpinene | Monoterpenes | 9.17 | 1034 | 7.51 ± 2.14 | 7.97 ± 1.66 | 6.47 ± 1.09 | 6.67 ± 1.57 |
| α-phellandrene | Monoterpenes | 8.89 | 1023 | 8.76 ± 1.76 | 11.29 ± 1.14 | 9.95 ± 0.72 | 10.66 ± 1.41 |
| α-pinene* | Monoterpenes | 7.16 | 954 | 28.24 ± 9.78 | 22.78 ± 3.02 | 17.84 ± 2.3 | 19.7 ± 4.75 |
| α-terpineol* | Monoterpenes | 13.79 | 1221 | 8.4 ± 1.29 | 9.39 ± 0.79 | 9.78 ± 1.21 | 10.71 ± 2.2 |
| α-thujene* | Monoterpenes | 6.99 | 947 | 14.5 ± 5.81 | 17.1 ± 4.14 | 13.59 ± 1.75 | 16.12 ± 2.96 |
| β-myrcene | Monoterpenes | 8.57 | 1010 | 26.46 ± 4.66 | 35.3 ± 4.39 | 34.09 ± 5.33 | 36.55 ± 13.2 |
| β-terpinene | Monoterpenes | 8.33 | 1001 | 7.33 ± 1.41 | 7.02 ± 1.01 | 5.18 ± 0.84 | 8.29 ± 2.37 |
| β-thujone | Monoterpenes | 11.66 | 1133 | 179.7 ± 67.27 | 463.07 ± 118.74 | 346.81 ± 64.7 | 337.91 ± 74.72 |
| γ-terpinene* | Monoterpenes | 10.24 | 1076 | 41.32 ± 11.42 | 54.32 ± 9.74 | 41.55 ± 6.38 | 53.64 ± 21.81 |
| γ-terpineol | Monoterpenes | 14.02 | 1230 | 0.75 ± 0.75 | 1.17 ± 0.73 | 0.2 ± 0.2 | 0.43 ± 0.43 |
| Unknown 1 | Other | 7.26 | 958 | 4.13 ± 1.6 | 3.97 ± 0.83 | 3.58 ± 0.79 | 4.01 ± 1.78 |
| Unknown 2 | Other | 18.91 | 1451 | 2.19 ± 1.08 | 1.85 ± 0.73 | 2.27 ± 0.76 | 2.05 ± 1.37 |
| Unknown 3 | Other | 19.54 | 1480 | 3.87 ± 1.28 | 3.51 ± 0.72 | 5.23 ± 2.28 | 3.77 ± 1.2 |
| 7-epi-silphiperfol-5-ene | Sesquiterpenes | 17.35 | 1378 | 1.73 ± 0.79 | 1.23 ± 0.33 | 1.56 ± 0.43 | 2.09 ± 0.94 |
| Alloaromadendrene | Sesquiterpenes | 19.15 | 1462 | 3.85 ± 1.19 | 3.04 ± 0.63 | 2.7 ± 0.58 | 3.53 ± 0.93 |
| Copaene | Sesquiterpenes | 17.64 | 1391 | 5.01 ± 0.91 | 4.71 ± 0.93 | 4.57 ± 0.73 | 5.35 ± 2.6 |
| germacrene D | Sesquiterpenes | 19.84 | 1495 | 4.02 ± 1.88 | 2.77 ± 0.87 | 2.78 ± 0.77 | 9.32 ± 7.28 |
| Unknown sesquiterpene 1 | Sesquiterpenes | 16.94 | 1359 | 4.29 ± 1.3 | 6.59 ± 1.49 | 5.69 ± 0.92 | 5.91 ± 2.11 |
| Unknown sesquiterpene 2 | Sesquiterpenes | 17.23 | 1373 | 1.28 ± 0.7 | 1.1 ± 0.4 | 1.67 ± 0.69 | 1.93 ± 0.92 |
| α -muurolene | Sesquiterpenes | 20.1 | 1508 | 7.47 ± 2.2 | 9.63 ± 1.66 | 8.85 ± 1.47 | 8.13 ± 3.1 |

**Table S2-1 *(continued)***: *This section shows the results for the third part of Table S2-1.*

| Compound | Group | RT | KI | 1 | 2 | 3 | 6 |
|---|---|---|---|---|---|---|---|
| α-bourbonene | Sesquiterpenes | 17.94 | 1405 | 4.86 ± 2.68 | 3.15 ± 0.88 | 2.47 ± 0.77 | 5.16 ± 4 |
| α-cadinol | Sesquiterpenes | 21.82 | 1596 | 1.47 ± 0.65 | 2.81 ± 0.79 | 2.71 ± 0.77 | 5.45 ± 2.49 |
| α-caryophyllene* | Sesquiterpenes | 19.27 | 1468 | 4.59 ± 1.35 | 5.46 ± 2.13 | 7.4 ± 3.18 | 8.06 ± 5 |
| α-farnesene | Sesquiterpenes | 20.26 | 1516 | 18.25 ± 7.21 | 18.59 ± 4.62 | 23.48 ± 5.86 | 42.2 ± 23.94 |
| β-caryophyllene* | Sesquiterpenes | 18.57 | 1435 | 4.42 ± 1.76 | 10.01 ± 3.22 | 7.52 ± 2.56 | 33.89 ± 28.58 |
| β-cubebene | Sesquiterpenes | 18.06 | 1411 | 1.16 ± 0.65 | 3.32 ± 0.84 | 1.99 ± 0.49 | 2.43 ± 1.23 |
| β-selinene | Sesquiterpenes | 19.94 | 1500 | 10.99 ± 2.2 | 10.94 ± 1.31 | 9.93 ± 1.8 | 13.77 ± 5.03 |
| γ-cadinene | Sesquiterpenes | 20.46 | 1527 | 0.26 ± 0.26 | 0.51 ± 0.26 | 0.45 ± 0.21 | 0.9 ± 0.9 |
| γ-elemene | Sesquiterpenes | 18.81 | 1446 | 2.31 ± 0.76 | 2.39 ± 0.47 | 1.43 ± 0.38 | 2.5 ± 0.89 |
| δ-cadinene | Sesquiterpenes | 20.65 | 1536 | 6.2 ± 1.66 | 8.92 ± 1.69 | 8.02 ± 1.49 | 8.44 ± 3.55 |

#### 2.3.1. Tables

*2.3.1.1. Effects of chemotype and plot-level chemotype richness on traits of individual plants*

**Table S2-2:** Summary of generalized linear mixed-effect models (GLMM) testing chemotype identity (C), plot-level chemotype richness (CR), and the interaction between chemotype identity and plot-level chemotype richness (CP x CR) effects on the number of stems of *T. vulgare* plants at different time points in 2021 and 2022. Degrees of freedom, Wald's Chi-square statistics, and p-values are reported. Significant values (P <0.05) are reported in bold.

| **Number of stems** | | **Sampling date** | | | |
|---|---|---|---|---|---|
| | | **01.06.2021** | **22.06.2021** | **28.10.2021** | **06.07.2022** |
| **Factor** | **d.f.** | **χ2 (p-value)** | **χ2 (p-value)** | **χ2 (p-value)** | **χ2 (p-value)** |
| C | 5 | 24.99 **(<0.001)** | 21.49 **(<0.001)** | 1.72 (0.887) | 3.28 (0.657) |
| CR | 1 | 1.47 (0.225) | 0.79 (0.373) | 0.55 (0.459) | 4.67 **(0.031)** |
| C x CR | 5 | 1.69 (0.890) | 4.25 (0.514) | 3.40 (0.638) | 7.86 (0.164) |

**Table S2-3:** Summary of linear mixed-effect models (LMM) testing chemotype identity (C), plot-level chemotype richness (CR), and the interaction between chemotype identity and plot-level chemotype richness (CP x CR) effects on the height (cm) of *T. vulgare* plants at different time points in 2021 and 2022. Degrees of freedom, Wald's Chi-square statistics, and p-values are reported. Significant values (P <0.05) are reported in bold.

| **Height (cm)** | | **Sampling date** | | | | | |
|---|---|---|---|---|---|---|---|
| | | **01.06.2021** | **22.06.2021** | **28.10.2021** | **14.05.2022** | **06.07.2022** | **05.10.2022** |
| **Factor** | **d.f.** | **χ2 (p-value)** | **χ2 (p-value)** | **χ2 (p-value)** | **χ2 (p-value)** | **χ2 (p-value)** | **χ2 (p-value)** |
| C | 5 | 5.74 (0.332) | 10.54 (0.061) | 12.26 **(0.031)** | 5.75 (0.332) | 1.58 (0.903) | 2.53 (0.772) |
| CR | 1 | 0.42 (0.519) | 0.03 (0.852) | 0.00 (0.999) | 1.66 (0.198) | 0.30 (0.584) | 0.06 (0.807) |
| C x CR | 5 | 7.97 (0.158) | 5.42 (0.367) | 2.25 (0.814) | 3.45 (0.632) | 7.24 (0.203) | 3.47 (0.628) |

**Table S2-4:** Summary of linear mixed-effect models (LMM) testing chemotype identity (C), plot-level chemotype richness (CR), and the interaction between chemotype identity and plot-level chemotype richness (CP x CR) effects on the square root above-ground dry weight (g) in 2021, and the square root above-ground fresh weight (g) in 2022 of *T. vulgare* plants. Degrees of freedom, Wald's Chi-square statistics, and p-values are reported. Significant values (P <0.05) are reported in bold.

| **sqrt(above-ground dry weight (g))** | | **Sampling date 28.10.2021** | **sqrt(above-ground fresh weight (g))** | **Sampling date 05.10.2022** | |
|---|---|---|---|---|---|
| **Factor** | **d.f.** | **$\chi 2$ (p-value)** | **Factor** | **d.f.** | **$\chi 2$ (p-value)** |
| C | 5 | 9.35 (0.096) | C | 5 | 2.65 (0.753) |
| CR | 1 | 0.20 (0.652) | CR | 1 | 0.62 (0.430) |
| C x CR | 5 | 5.00 (0.416) | C x CR | 5 | 1.79 (0.877) |

**Table S2-5:** Summary of generalized linear mixed-effect models (GLMM) and linear mixed-effect models (LMM) testing chemotype identity (C), plot-level chemotype richness (CR), and the interaction between chemotype identity and plot-level chemotype richness (CP x CR) effects on the cumulative number of flower heads in 2021 and the flowering index in 2022 of *T. vulgare* plants, respectively. Degrees of freedom, Wald's Chi-square statistics, and p-values are reported. Significant values (P <0.05) are reported in bold.

| **Cumulative flower heads** | | **Sampling date 01.06.2021 - 28.10.2021** | **Flowering index** | | **Sampling date 06.07.2022** |
|---|---|---|---|---|---|
| **Factor** | **d.f.** | **χ2 (p-value)** | **Factor** | **d.f.** | **χ2 (p-value)** |
| C | 5 | 55.08 **(<0.001)** | C | 5 | 51.07 **(<0.001)** |
| CR | 1 | 1.40 (0.236) | CR | 1 | 0.76 (0.382) |
| C x CR | 5 | 779.33 **(<0.001)** | C x CR | 5 | 14.93 **(0.011)** |

*2.3.1.2. Effects of daughter and plot-level chemotype richness on traits of individual plants*

**Table S2-6:** Summary of generalized linear mixed-effect models (GLMM) testing daughter identity (D), plot-level chemotype richness (CR), and the interaction between daughter identity and plot-level chemotype richness (D x CR) effects on the number of stems, and summary of linear mixed-effect models (LMM) on the effects on height (cm), the squared root above-ground dry weight (g), and the cumulative number of flower heads of *T. vulgare* plants in 2021. Degrees of freedom, Wald's Chi-square statistics, and p-values are reported. Significant values (P <0.05) are reported in bold.

| | | | **Number of stems** | **Height (cm)** | **sqrt(above-ground dry weight (g))** | **Cumulative number of flower heads** |
|---|---|---|---|---|---|---|
| | | | **28.10.2021** | **28.10.2021** | **28.10.2021** | **01.06.2021 – 28.10.2021** |
| | **Factor** | **d.f.** | **χ2 (p-value)** | **χ2 (p-value)** | **χ2 (p-value)** | **χ2 (p-value)** |
| Athu-Bthu | D | 2 | 40.19 **(<0.001)** | 11.34 **(0.003)** | 2.26 (0.322) | 5.28 (0.071) |
| | CR | 1 | 0.16 (0.693) | 1.65 (0.199) | 1.12 (0.290) | 1.66 (0.198) |
| | D x CR | 2 | 2.56 (0.278) | 1.08 (0.581) | 2.24 (0.326) | 0.86 (0.651) |
| Bthu-high | D | 2 | 36.72 **(<0.001)** | 27.76 **(<0.001)** | 32.86 **(<0.001)** | 37.16 **(<0.001)** |
| | CR | 1 | 0.52 (0.469) | 0.20 (0.652) | 0.14 (0.709) | 2.77 (0.096) |
| | D x CR | 2 | 1.69 (0.429) | 0.07 (0.965) | 4.30 (0.117) | 3.60 (0.165) |
| Bthu-low | D | 2 | 22.69 **(<0.001)** | 54.40 **(<0.001)** | 122.70 **(<0.001)** | 50.48 **(<0.001)** |
| | CR | 1 | 0.17 (0.679) | 0.42 (0.517) | 0.61 (0.435) | 6.54 **(0.010)** |
| | D x CR | 2 | 2.45 (0.294) | 1.46 (0.480) | 6.03 **(0.049)** | 6.90 **(0.032)** |
| Chrys-acet | D | 2 | 33.27 **(<0.001)** | 6.84 **(0.033)** | 23.25 **(<0.001)** | 1.70 (0.428) |
| | CR | 1 | 1.11 (0.292) | 0.07 (0.789) | 0.51 (0.477) | 0.67 (0.412) |
| | D x CR | 2 | 1.30 (0.522) | 0.70 (0.704) | 2.78 (0.249) | 1.72 (0.422) |
| Mixed-high | D | 2 | 4.01 (0.135) | 50.04 (0.818) | 38.77 **(<0.001)** | 15.43 **(<0.001)** |
| | CR | 1 | 1.43 (0.232) | 0.24 (0.622) | 2.17 (0.141) | 0.88 (0.347) |
| | D x CR | 2 | 1.29 (0.525) | 1.41 (0.494) | 2.98 (0.225) | 2.14 (0.344) |
| Mixed-low | D | 2 | 35.10 **(<0.001)** | 50.04 **(<0.001)** | 58.25 **(<0.001)** | 12.73 **(0.002)** |
| | CR | 1 | 0.23 (0.633) | 0.97 (0.324) | 0.57 (0.450) | 1.43 (0.232) |
| | D x CR | 2 | 0.93 (0.628) | 1.76 (0.414) | 1.97 (0.373) | 0.24 (0.888) |
| All chemotypes | D | 17 | 201.42 **(<0.001)** | 299.29 **(<0.001)** | 574.63 **(<0.001)** | 253.23 **(<0.001)** |
| | CR | 1 | 0.44 (0.508) | 0.00 (0.989) | 0.21 (0.643) | 2.00 (0.158) |
| | D x CR | 17 | 13.68 (0.690) | 8.23 (0.961) | 28.37 (**0.041**) | 32.57 **(0.001)** |

*2.3.1.3. Effects of chemotype and plot-level chemotype richness on plot-level means*

**Table S2-7:** Summary of linear models (LM) testing Block (B), chemotype presence (CP), plot-level chemotype richness (CR), and the interaction between chemotype presence and plot-level chemotype richness (CP x CR) effects on averaged number of stems of *T. vulgare* plants at plot level at different time points in 2021 and 2022. Degrees of freedom, F-statistics, and p-values are reported. Significant values (P <0.05) are reported in bold.

| **Average plot-level number of stems** | | | **Sampling date** | | | |
|---|---|---|---|---|---|---|
| | | | **01.06.2021** | **22.06.2021** | **28.10.2021** | **06.07.2022** |
| **Chemotype** | **Factor** | **d.f.** | **F (p-value)** | **F (p-value)** | **F (p-value)** | **F (p-value)** |
| Athu-Bthu | B | 5, 75 | 2.16 (0.067) | 11.58 **(<0.001)** | 2.73 **(0.026)** | 8.23 **(<0.001)** |
| | CR | 1, 75 | 4.49 **(0.037)** | 3.06 (0.084) | 0.23 (0.634) | 3.37 (0.070) |
| | CP | 1, 75 | 4.42 **(0.039)** | 0.00 (0.950) | 2.30 (0.134) | 2.14 (0.148) |
| | CP x CR | 1, 75 | 0.02 (0.896) | 0.68 (0.413) | 1.19 (0.279) | 0.07 (0.793) |
| Bthu-high | B | 5, 75 | 2.12 (0.072) | 12.92 **(<0.001)** | 2.73 **(0.026)** | 8.36 **(<0.001)** |
| | CR | 1, 75 | 4.40 **(0.039)** | 3.41 (0.069) | 0.23 (0.634) | 3.42 (0.068) |
| | CP | 1, 75 | 0.63 (0.429) | 8.10 **(0.006)** | 0.22 (0.640) | 2.28 (0.135) |
| | CP x CR | 1, 75 | 2.24 (0.139) | 1.37 (0.245) | 3.20 (0.078) | 1.27 (0.263) |
| Bthu-low | B | 5, 75 | 2.76 **(0.024)** | 11.56 **(<0.001)** | 2.65 **(0.029)** | 8.06 **(<0.001)** |
| | CR | 1, 75 | 5.72 **(0.019)** | 3.05 (0.085) | 0.22 (0.639) | 3.30 (0.073) |
| | CP | 1, 75 | 26.14 **(<0.001)** | 0.59 (0.445) | 1.24 (0.269) | 0.00 (0.984) |
| | CP x CR | 1, 75 | 0.00 (0.941) | 0.00 (0.987) | 0.02 (0.879) | 0.72 (0.398) |
| Chrys-acet | B | 5, 75 | 2.06 (0.080) | 11.49 **(<0.001)** | 2.73 **(0.025)** | 8.13 **(<0.001)** |
| | CR | 1, 75 | 4.27 **(0.042)** | 3.03 (0.086) | 0.22 (0.641) | 3.33 (0.072) |
| | CP | 1, 75 | 0.31 (0.581) | 0.01 (0.923) | 0.05 (0.825) | 1.12 (0.292) |
| | CP x CR | 1, 75 | 0.28 (0.598) | 0.11 (0.743) | 0.37 (0.546) | 0.19 (0.665) |
| Mixed-high | B | 5, 75 | 2.56 **(0.033)** | 13.07 **(<0.001)** | 2.63 **(0.030)** | 8.68 **(<0.001)** |
| | CR | 1, 75 | 5.34 **(0.024)** | 3.45 (0.067) | 0.22 (0.641) | 3.55 (0.063) |
| | CP | 1, 75 | 12.85 **(<0.001)** | 10.35 **(0.002)** | 0.19 (0.661) | 2.21 (0.141) |
| | CP x CR | 1, 75 | 6.68 **(0.012)** | 0.08 (0.783) | 0.30 (0.583) | 4.29 **(0.042)** |
| Mixed-low | B | 5, 75 | 2.06 (0.080) | 11.54 **(<0.001)** | 2.62 **(0.031)** | 8.09 **(<0.001)** |
| | CR | 1, 75 | 4.26 **(0.042)** | 3.04 (0.085) | 0.22 (0.642) | 3.31 (0.073) |
| | CP | 1, 75 | 0.15 (0.703) | 0.06 (0.804) | 0.07 (0.792) | 0.10 (0.755) |
| | CP x CR | 1, 75 | 0.25 (0.621) | 0.34 (0.560) | 0.02 (0.895) | 0.88 (0.350) |

**Table S2-8:** Summary of linear models (LM) testing Block (B), chemotype presence (CP), plot-level chemotype richness (CR), and the interaction between chemotype presence and plot-level chemotype richness (CP x CR) effects on averaged height (cm) of *T. vulgare* plants at plot level at different time points in 2021 and 2022. Degrees of freedom, F-statistics, and p-values are reported. Significant values (P <0.05) are reported in bold.

| **Average plot-level height (cm)** | | | **Sampling date** | | | | | |
|---|---|---|---|---|---|---|---|---|
| | | | **01.06.2021** | **22.06.2021** | **28.10.2021** | **14.05.2022** | **06.07.2022** | **05.10.2022** |
| **Chemotype** | **Factor** | **d.f.** | **F (p-value)** | **F (p-value)** | **F (p-value)** | **F (p-value)** | **F (p-value)** | **F (p-value)** |
| Athu-Bthu | B | 5, 75 | 3.06 **(0.014)** | 2.25 (0.058) | 1.71 (0.143) | 0.25 (0.940) | 5.36 **(<0.001)** | 3.96 **(0.003)** |
| | CR | 1, 75 | 0.15 (0.698) | 0.21 (0.648) | 0.09 (0.770) | 1.39 (0.242) | 0.28 (0.595) | 0.09 (0.766) |
| | CP | 1, 75 | 6.70 **(0.012)** | 0.94 (0.335) | 1.80 (0.183) | 0.74 (0.394) | 1.03 (0.313) | 1.69 (0.198) |
| | CP x CR | 1, 75 | 3.33 (0.072) | 0.31 (0.581) | 0.01 (0.920) | 0.01 (0.923) | 0.56 (0.458) | 0.02 (0.902) |
| Bthu-high | B | 5, 75 | 2.80 **(0.022)** | 2.23 (0.060) | 1.70 (0.144) | 0.25 (0.938) | 5.46 **(<0.001)** | 3.90 **(0.003)** |
| | CR | 1, 75 | 0.14 (0.710) | 0.21 (0.650) | 0.08 (0.770) | 1.42 (0.237) | 0.29 (0.592) | 0.09 (0.767) |
| | CP | 1, 75 | 2.69 (0.105) | 0.56 (0.454) | 0.62 (0.434) | 1.46 (0.231) | 2.55 (0.115) | 0.55 (0.462) |
| | CP x CR | 1, 75 | 0.33 (0.567) | 0.01 (0.936) | 1.00 (0.322) | 0.70 (0.404) | 0.50 (0.483) | 0.04 (0.836) |
| Bthu-low | B | 5, 75 | 2.78 **(0.023)** | 2.64 **(0.030)** | 2.24 (0.059) | 0.24 (0.942) | 5.27 **(<0.001)** | 4.08 **(0.002)** |
| | CR | 1, 75 | 0.14 (0.712) | 0.25 (0.621) | 0.11 (0.738) | 1.38 (0.244) | 0.28 (0.598) | 0.09 (0.762) |
| | CP | 1, 75 | 0.80 (0.375) | 14.41 **(<0.001)** | 24.00 **(<0.001)** | 0.10 (0.749) | 0.38 (0.542) | 3.89 (0.052) |
| | CP x CR | 1, 75 | 1.50 (0.225) | 0.02 (0.891) | 1.65 (0.203) | 0.02 (0.887) | 0.00 (0.985) | 0.01 (0.912) |
| Chrys-acet | B | 5, 75 | 2.90 **(0.019)** | 2.48 **(0.039)** | 1.68 (0.149) | 0.25 (0.937) | 5.32 **(<0.001)** | 3.88 **(0.003)** |
| | CR | 1, 75 | 0.14 (0.706) | 0.23 (0.631) | 0.08 (0.772) | 1.43 (0.235) | 0.28 (0.596) | 0.09 (0.768) |
| | CP | 1, 75 | 0.71 (0.401) | 5.54 **(0.021)** | 0.44 (0.508) | 2.85 (0.096) | 1.08 (0.301) | 0.00 (0.963) |
| | CP x CR | 1, 75 | 4.87 **(0.030)** | 3.80 (0.055) | 0.14 (0.708) | 0.05 (0.818) | 0.02 (0.873) | 0.22 (0.642) |
| Mixed-high | B | 5, 75 | 3.02 **(0.015)** | 3.00 **(0.016)** | 1.89 (0.101) | 0.25 (0.941) | 5.25 **(<0.001)** | 4.07 **(0.002)** |
| | CR | 1, 75 | 0.15 (0.700) | 0.28 (0.598) | 0.10 (0.759) | 1.39 (0.242) | 0.28 (0.599) | 0.09 (0.763) |
| | CP | 1, 75 | 8.39 **(0.005)** | 25.18 **(<0.001)** | 8.36 **(0.005)** | 0.38 (0.541) | 0.06 (0.813) | 3.77 (0.056) |
| | CP x CR | 1, 75 | 0.74 (0.393) | 1.72 (0.194) | 1.42 (0.237) | 0.36 (0.549) | 0.04 (0.851) | 0.06 (0.807) |
| Mixed-low | B | 5, 75 | 3.27 **(0.010)** | 2.31 (0.052) | 1.72 (0.140) | 0.27 (0.930) | 5.36 **(<0.001)** | 3.93 **(0.003)** |
| | CR | 1, 75 | 0.16 (0.688) | 0.22 (0.644) | 0.09 (0.769) | 1.51 (0.223) | 0.28 (0.595) | 0.09 (0.767) |
| | CP | 1, 75 | 15.61 **(<0.001)** | 2.21 (0.142) | 2.35 (0.129) | 5.82 **(0.018)** | 1.58 (0.212) | 0.28 (0.598) |
| | CP x CR | 1, 75 | 0.29 (0.589) | 1.10 (0.297) | 0.00 (0.967) | 1.27 (0.263) | 0.12 (0.734) | 0.73 (0.365) |

**Table S2-9:** Summary of linear models (LM) testing Block (B), chemotype presence (CP), plot-level chemotype richness (CR), and the interaction between chemotype presence and plot-level chemotype richness (CP x CR) effects on the logarithm of averaged above-ground dry weight (g) in 2021, and the logarithm of averaged above-ground fresh weight (g) in 2022 of *T. vulgare* plants at plot level. Degrees of freedom, F-statistics, and p-values are reported. Significant values (P <0.05) are reported in bold.

| **log(Average plot-level above-ground dry weight (g))** | | | **Sampling date 28.10.2021** | **log(Average plot-level above-ground fresh weight (g))** | | | **Sampling date 01.06.2021** |
|---|---|---|---|---|---|---|---|
| **Chemotype** | **Factor** | **d.f.** | **F (p-value)** | **Chemotype** | **Factor** | **d.f.** | **F (p-value)** |
| Athu-Bthu | B | 5, 75 | 1.61 (0.167) | Athu-Bthu | B | 5, 75 | 2.94 **(0.018)** |
| | CR | 1, 75 | 0.06 (0.811) | | CR | 1, 75 | 1.07 (0.304) |
| | CP | 1, 75 | 9.89 **(0.002)** | | CP | 1, 75 | 7.24 **(0.009)** |
| | CP x CR | 1, 75 | 2.01 (0.161) | | CP x CR | 1, 75 | 0.19 (0.663) |
| Bthu-high | B | 5, 75 | 1.40 (0.232) | Bthu-high | B | 5, 75 | 2.67 **(0.028)** |
| | CR | 1, 75 | 0.05 (0.824) | | CR | 1, 75 | 0.98 (0.326) |
| | CP | 1, 75 | 0.10 (0.746) | | CP | 1, 75 | 0.00 (0.949) |
| | CP x CR | 1, 75 | 0.63 (0.429) | | CP x CR | 1, 75 | 0.00 (0.950) |
| Bthu-low | B | 5, 75 | 2.51 **(0.037)** | Bthu-low | B | 5, 75 | 3.14 **(0.012)** |
| | CR | 1, 75 | 0.09 (0.766) | | CR | 1, 75 | 1.15 (0.287) |
| | CP | 1, 75 | 59.54 **(<0.001)** | | CP | 1, 75 | 10.70 **(0.002)** |
| | CP x CR | 1, 75 | 0.89 (0.349) | | CP x CR | 1, 75 | 2.61 (0.110) |
| Chrys-acet | B | 5, 75 | 1.39 (0.236) | Chrys-acet | B | 5, 75 | 2.75 **(0.024)** |
| | CR | 1, 75 | 0.05 (0.824) | | CR | 1, 75 | 1.00 (0.319) |
| | CP | 1, 75 | 0.09 (0.761) | | CP | 1, 75 | 1.37 (0.245) |
| | CP x CR | 1, 75 | 0.06 (0.803) | | CP x CR | 1, 75 | 0.91 (0.342) |
| Mixed-high | B | 5, 75 | 1.43 (0.223) | Mixed-high | B | 5, 75 | 2.71 **(0.026)** |
| | CR | 1, 75 | 0.05 (0.822) | | CR | 1, 75 | 0.99 (0.323) |
| | CP | 1, 75 | 1.29 (0.259) | | CP | 1, 75 | 0.06 (0.809) |
| | CP x CR | 1, 75 | 0.76 (0.385) | | CP x CR | 1, 75 | 0.923 (0.340) |
| Mixed-low | B | 5, 75 | 1.41 (0.229) | Mixed-low | B | 5, 75 | 2.74 **(0.025)** |
| | CR | 1, 75 | 0.05 (0.823) | | CR | 1, 75 | 1.00 (0.320) |
| | CP | 1, 75 | 0.83 (0.364) | | CP | 1, 75 | 1.87 (0.175) |
| | CP x CR | 1, 75 | 0.39 (0.532) | | CP x CR | 1, 75 | 0.20 (0.660) |

**Table S2-10:** Summary of linear models (LM) testing Block (B), chemotype presence (CP), plot-level chemotype richness (CR), and the interaction between chemotype presence and plot-level chemotype richness (CP x CR) effects on the logarithm of cumulative number of flower heads in 2021 and flowering index in 2022 of *T. vulgare* plants at plot level. Degrees of freedom, F-statistics, and p-values are reported. Significant values (P <0.05) are reported in bold.

| **log(plot-level cumulative flower heads)** | | | **Sampling date 01.06.2021 - 28.10.2021** | **Plot-level flowering index** | | | **Sampling date 06.07.2022** |
|---|---|---|---|---|---|---|---|
| **Chemotype** | **Factor** | **d.f.** | **F (p-value)** | **Chemotype** | **Factor** | **d.f.** | **F (p-value)** |
| Athu-Bthu | B | 5, 75 | 0.30 (0.910) | Athu-Bthu | B | 5, 75 | 5.31 **(<0.001)** |
| | CR | 1, 75 | 1.49 (0.225) | | CR | 1, 75 | 0.17 (0.682) |
| | CP | 1, 75 | 2.21 (0.141) | | CP | 1, 75 | 0.03 (0.855) |
| | CP x CR | 1, 75 | 3.93 (0.052) | | CP x CR | 1, 75 | 1.14 (0.289) |
| Bthu-high | B | 5, 75 | 0.31 (0.903) | Bthu-high | B | 5, 75 | 5.85 **(<0.001)** |
| | CR | 1, 75 | 1.56 (0.216) | | CR | 1, 75 | 0.19 (0.667) |
| | CP | 1, 75 | 2.36 (0.129) | | CP | 1, 75 | 5.88 **(0.018)** |
| | CP x CR | 1, 75 | 7.08 **(0.010)** | | CP x CR | 1, 75 | 2.98 (0.088) |
| Bthu-low | B | 5, 75 | 0.35 (0.882) | Bthu-low | B | 5, 75 | 6.36 **(<0.001)** |
| | CR | 1, 75 | 1.72 (0.194) | | CR | 1, 75 | 0.20 (0.653) |
| | CP | 1, 75 | 13.08 **(<0.001)** | | CP | 1, 75 | 16.27 **(<0.001)** |
| | CP x CR | 1, 75 | 5.25 **(0.025)** | | CP x CR | 1, 75 | 0.00 (0.981) |
| Chrys-acet | B | 5, 75 | 0.33 (0.894) | Chrys-acet | B | 5, 75 | 5.24 **(<0.001)** |
| | CR | 1, 75 | 1.63 (0.206) | | CR | 1, 75 | 0.17 (0.684) |
| | CP | 1, 75 | 12.12 **(<0.001)** | | CP | 1, 75 | 0.06 (0.805) |
| | CP x CR | 1, 75 | 1.32 (0.255) | | CP x CR | 1, 75 | 0.08 (0.773) |
| Mixed-high | B | 5, 75 | 0.30 (0.913) | Mixed-high | B | 5, 75 | 7.99 **(<0.001)** |
| | CR | 1, 75 | 1.47 (0.228) | | CR | 1, 75 | 0.26 (0.615) |
| | CP | 1, 75 | 0.01 (0.939) | | CP | 1, 75 | 37.90 **(<0.001)** |
| | CP x CR | 1, 75 | 4.98 **(0.029)** | | CP x CR | 1, 75 | 1.69 (0.197) |
| Mixed-low | B | 5, 75 | 0.31 (0.903) | Mixed-low | B | 5, 75 | 6.80 **(<0.001)** |
| | CR | 1, 75 | 1.56 (0.216) | | CR | 1, 75 | 0.22 (0.643) |
| | CP | 1, 75 | 0.00 (0.975) | | CP | 1, 75 | 22.42 **(<0.001)** |
| | CP x CR | 1, 75 | 9.60 **(0.003)** | | CP x CR | 1, 75 | 0.12 (0.727) |

*2.3.1.4. Overyielding calculations*

**Table S2-11:** Overyielding indices (OI) were calculated for each plant trait at plot-level (above-ground dry weight (g) - 2021, above-ground fresh weight (g) - 2022, the cumulative number of flower heads - 2021, and flowering index -2022 of *T. vulgare* plants). The table is partitioned into three sections by block.

| Block | Plot | Level | Plot-level above-ground dry weight (2021) | Plot-level above-ground fresh weight (2022) | Plot-level cumulative number of flower heads (2021) | Plot-level flowering index (2022) |
|---|---|---|---|---|---|---|
| 1 | 1 | 3 | 0.19 | 0.09 | -0.05 | -0.37 |
| 1 | 2 | 2 | -0.12 | -0.22 | 0.06 | -0.02 |
| 1 | 3 | 3 | -0.07 | -0.09 | -0.01 | -0.20 |
| 1 | 4 | 2 | 0.17 | -0.08 | -0.03 | -0.06 |
| 1 | 5 | 3 | -0.14 | -0.10 | -0.13 | -0.55 |
| 1 | 6 | 6 | -0.21 | -0.13 | -0.12 | 0.21 |
| 1 | 7 | 6 | 0.09 | -0.10 | -0.06 | -0.41 |
| 1 | 8 | 2 | -0.02 | -0.16 | -0.04 | -0.45 |
| 1 | 9 | 1 | 0.12 | -0.02 | -0.08 | 0.06 |
| 1 | 10 | 1 | 0.18 | -0.12 | -0.09 | 0.16 |
| 1 | 11 | 2 | -0.04 | -0.37 | -0.06 | -0.53 |
| 1 | 12 | 2 | 0.02 | 0.00 | -0.10 | -0.13 |
| 1 | 13 | 3 | -0.10 | -0.16 | 0.01 | -0.01 |
| 1 | 14 | 3 | 0.07 | 0.17 | -0.07 | 0.10 |
| 2 | 15 | 2 | 0.12 | 0.02 | 0.01 | 0.19 |
| 2 | 16 | 6 | 0.24 | -0.02 | -0.02 | -0.30 |
| 2 | 17 | 2 | -0.22 | -0.09 | -0.01 | 0.10 |
| 2 | 18 | 3 | 0.01 | -0.24 | -0.07 | -0.21 |
| 2 | 19 | 2 | -0.28 | -0.47 | -0.17 | -0.38 |
| 2 | 20 | 2 | 0.25 | 0.01 | -0.05 | 0.11 |
| 2 | 21 | 3 | 0.39 | -0.03 | 0.00 | -0.51 |
| 2 | 22 | 6 | -0.31 | -0.23 | -0.03 | -0.04 |
| 2 | 23 | 3 | 0.04 | -0.06 | -0.06 | -0.20 |
| 2 | 24 | 1 | -0.07 | -0.18 | -0.13 | 0.05 |
| 2 | 25 | 1 | 0.07 | -0.17 | -0.04 | -0.08 |
| 2 | 26 | 3 | 0.08 | 0.05 | -0.11 | -0.56 |
| 2 | 27 | 3 | -0.35 | -0.09 | -0.05 | -0.12 |
| 2 | 28 | 2 | 0.08 | -0.13 | -0.05 | -0.52 |

**Table S2-11** ***(continued)*****:** *This section shows the results for the second part of Table S2-11, partitioned by block.*

| Block | Plot | Level | Plot-level above-ground dry weight (2021) | Plot-level above-ground fresh weight (2022) | Plot-level cumulative number of flower heads (2021) | Plot-level flowering index (2022) |
|---|---|---|---|---|---|---|
| 3 | 29 | 2 | 0.18 | 0.44 | -0.01 | -0.12 |
| 3 | 30 | 3 | 0.29 | 0.28 | -0.08 | -0.26 |
| 3 | 31 | 2 | -0.19 | 0.04 | -0.03 | 0.09 |
| 3 | 32 | 6 | -0.42 | -0.37 | -0.09 | -0.42 |
| 3 | 33 | 2 | -0.17 | -0.27 | -0.16 | -0.46 |
| 3 | 34 | 1 | 0.10 | 0.03 | -0.01 | 0.01 |
| 3 | 35 | 1 | 0.09 | 0.07 | -0.02 | -0.24 |
| 3 | 36 | 3 | -0.18 | 0.12 | 0.00 | -0.40 |
| 3 | 37 | 3 | -0.21 | 0.06 | 0.01 | -0.41 |
| 3 | 38 | 2 | 0.01 | 0.02 | 0.10 | -0.16 |
| 3 | 39 | 3 | -0.07 | 0.26 | 0.14 | -0.05 |
| 3 | 40 | 3 | -0.05 | 0.11 | -0.03 | -0.18 |
| 3 | 41 | 2 | -0.20 | -0.12 | 0.00 | 0.16 |
| 3 | 42 | 6 | 0.40 | 0.44 | -0.09 | -0.28 |
| 4 | 43 | 1 | -0.09 | -0.07 | 0.02 | 0.24 |
| 4 | 44 | 1 | -0.10 | -0.03 | 0.01 | -0.01 |
| 4 | 45 | 2 | -0.27 | -0.23 | 0.06 | -0.57 |
| 4 | 46 | 2 | -0.18 | 0.12 | 0.02 | -0.14 |
| 4 | 47 | 6 | -0.16 | 0.10 | 0.09 | -0.04 |
| 4 | 48 | 3 | -0.04 | 0.11 | 0.00 | -0.30 |
| 4 | 49 | 3 | -0.09 | 0.15 | 0.14 | 0.38 |
| 4 | 50 | 6 | -0.18 | -0.10 | 0.02 | -0.15 |
| 4 | 51 | 3 | -0.02 | 0.24 | 0.02 | -0.52 |
| 4 | 52 | 3 | 0.01 | 0.04 | 0.08 | -0.30 |
| 4 | 53 | 2 | -0.09 | 0.30 | 0.11 | -0.05 |
| 4 | 54 | 2 | -0.03 | 0.08 | 0.07 | -0.23 |
| 4 | 55 | 2 | 0.00 | -0.01 | 0.03 | 0.21 |
| 4 | 56 | 3 | -0.10 | -0.14 | 0.08 | -0.23 |

**Table S2-11 *(continued)***: *This section shows the results for the third part of Table S2-11, partitioned by block.*

| Block | Plot | Level | Plot-level above-ground dry weight (2021) | Plot-level above-ground fresh weight (2022) | Plot-level cumulative number of flower heads (2021) | Plot-level flowering index (2022) |
|---|---|---|---|---|---|---|
| 5 | 57 | 1 | -0.09 | 0.21 | 0.05 | 0.09 |
| 5 | 58 | 3 | -0.08 | 0.06 | 0.03 | -0.49 |
| 5 | 59 | 6 | 0.01 | -0.09 | 0.08 | -0.28 |
| 5 | 60 | 3 | 0.23 | 0.28 | -0.04 | 0.28 |
| 5 | 61 | 3 | -0.03 | 0.13 | 0.01 | -0.04 |
| 5 | 62 | 2 | -0.01 | 0.06 | 0.14 | -0.44 |
| 5 | 63 | 1 | 0.07 | 0.18 | 0.13 | -0.05 |
| 5 | 64 | 3 | -0.23 | -0.26 | 0.08 | -0.43 |
| 5 | 65 | 6 | 0.00 | -0.16 | 0.01 | -0.14 |
| 5 | 66 | 3 | 0.00 | 0.20 | 0.01 | 0.02 |
| 5 | 67 | 2 | -0.15 | 0.02 | 0.15 | 0.14 |
| 5 | 68 | 2 | 0.13 | 0.13 | 0.03 | -0.43 |
| 5 | 69 | 2 | -0.08 | 0.03 | 0.15 | -0.22 |
| 5 | 70 | 2 | -0.02 | 0.09 | 0.02 | -0.26 |
| 6 | 71 | 2 | -0.11 | -0.18 | 0.10 | -0.17 |
| 6 | 72 | 1 | -0.22 | 0.15 | 0.10 | -0.19 |
| 6 | 73 | 2 | 0.34 | 0.60 | 0.00 | -0.67 |
| 6 | 74 | 6 | 0.38 | -0.09 | 0.09 | -0.27 |
| 6 | 75 | 2 | -0.28 | 0.14 | 0.18 | 0.07 |
| 6 | 76 | 3 | -0.35 | -0.11 | 0.09 | -0.26 |
| 6 | 77 | 3 | -0.14 | -0.06 | 0.00 | 0.01 |
| 6 | 78 | 2 | -0.24 | -0.12 | 0.00 | 0.22 |
| 6 | 79 | 3 | -0.21 | -0.15 | 0.07 | -0.34 |
| 6 | 80 | 6 | 0.04 | -0.12 | 0.05 | -0.36 |
| 6 | 81 | 3 | 0.08 | 0.09 | 0.18 | -0.45 |
| 6 | 82 | 1 | -0.12 | 0.02 | 0.08 | -0.06 |
| 6 | 83 | 3 | -0.44 | -0.13 | 0.09 | -0.03 |
| 6 | 84 | 2 | -0.19 | -0.06 | 0.03 | -0.61 |

**Table S2-12:** Summary of linear mixed-effect models (LMM) testing plot-level chemotype richness (CR) effect on overyielding indices (OI). Degrees of freedom, Wald's Chi-square statistics, and p-values are reported. Significant values (P <0.05) are reported in bold.

| | **Factor** | **d.f.** | **χ2 (p-value)** |
|---|---|---|---|
| Above-ground dry weight (g) | CR | 1 | 0.07 (0.792) |
| Above-ground fresh weight (g) | CR | 1 | 1.09 (0.297) |
| Cumulative number of flower heads | CR | 1 | 2.57 (0.109) |
| Flowering index | CR | 1 | 0.68 (0.409) |

*2.3.1.5. Headspace VOC collection*

**Table S2-13:** List of leaf terpenoids identified only by hexane extraction (16), headspace terpenoids identified only in VOC collection (21), and terpenoids identified by both methods (28). Only terpenoids identified with the database were considered for comparison purposes; unknown terpenoids were excluded here.

| Leaf terpenoids (16) | Leaf and headspace terpenoids (21) | Headspace terpenoids (28) |
|---|---|---|
| 4-terpinenyl acetate | (-)-borneol | α-muurolene |
| artemisa alcohol | camphene | (-)-terpinen-4-ol |
| artemisia ketone | camphor | (+)-isothujol |
| artemisia acetate | chrysanthenyl acetate trans | 2-carene |
| cis-chrysanthenol | cis-abinene hydrate | 7-epi-ilphiperfol-5-ene |
| cis-myroxide | eucalyptol | alloaromadendrene |
| cis-verbanol acetate | limonene | bornyl acetate |
| eugenol | p-cimene | camphenol |
| methylisoborneol | pinocarvone | chrysanthenone |
| mustakone | sabinene | cis-3-dexen-1-ol, acetate |
| santolina triene | α-thujone | cis-abinol |
| trans-chrysanthenol | trans-sabinene hydrate | copaene |
| trans-abinyl acetate | trans-β-ocimene | germacrene D |
| yomogi alcohol | α-farnesene | terpinen-4-ol |
| α-chrysanthenyl acetate | α-pinene | terpinolene |
| β-pinene | α-terpinene | trans-myroxide |
| | α-thujene | α-bourbonene |
| | β-caryophyllene | α-cadinol |
| | β-myrcene | α-caryophyllene |
| | β-thujone | α-phellandrene |
| | γ-terpinene | α-terpineol |
| | | β-cubebene |
| | | β-elinene |
| | | β-terpinene |
| | | γ-cadinene |
| | | γ-elemene |
| | | γ-terpineol |
| | | δ-cadinene |

**Table S2-14:** Summary of linear models (LM) testing plot-level chemotype richness (CR) effect on theoretical plot-level leaf terpenoid diversity metrics. Degrees of freedom, F-statistics, p-values, and $R^2$ (when relevant) are reported. Significant values (P <0.05) are reported in bold.

| | Factor | d.f. | F (p-value) | $R^2$ |
|---|---|---|---|---|
| Squared plot-level theoretical total terpenoid concentration (nmol x $g^{-1}$) | CR | 1, 82 | 0.00 (0.967) | - |
| Plot-level theoretical terpenoid richness | CR | 1, 82 | 20.48 **(<0.001)** | 0.20 |
| Plot-level theoretical terpenoid Shannon diversity | CR | 1, 82 | 37.25 **(<0.001)** | 0.31 |
| Plot-level theoretical terpenoid evenness | CR | 1, 82 | 31.78 **(<0.001)** | 0.28 |

**Table S2-15:** Summary of linear mixed-effect models (LMM) testing plot-level chemotype richness (CR) effect on realized plot-level volatile diversity metrics. Degrees of freedom, Wald's Chi-square statistics, and p-values are reported. Significant values (P <0.05) are reported in bold.

| | **Factor** | **d.f.** | **$\chi$2 (p-value)** |
|---|---|---|---|
| Squared root of plot-level realized total volatile terpenoid emission (ng x $h^{-1}$) | CR | 1 | 0.02 (0.897) |
| Squared plot-level realized volatile terpenoid richness | CR | 1 | 0.36 (0.546) |
| Plot-level realized volatile terpenoid Shannon diversity | CR | 1 | 1.60 (0.206) |
| Plot-level theoretical terpenoid evenness | CR | 1 | 0.08 (0.780) |

#### 2.3.2. Figures

*2.3.2.1. Effects of chemotype and plot-level chemotype richness on traits of individual plants*

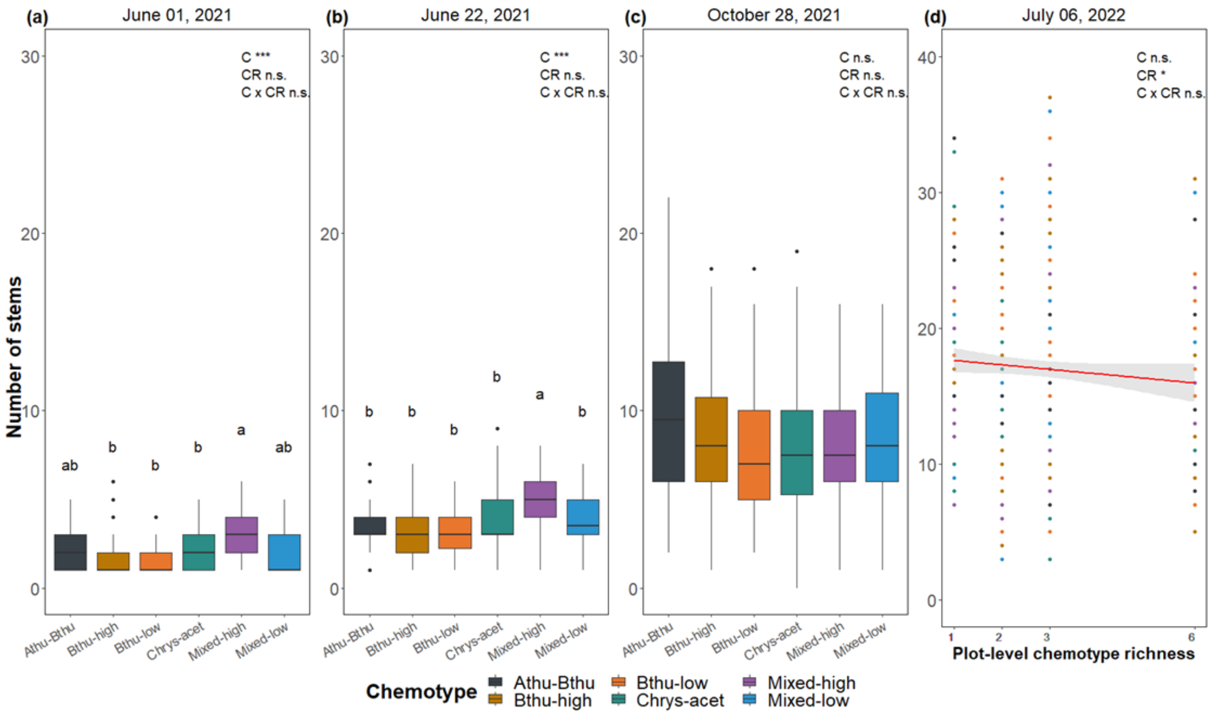


**Figure S2-1:** Effect of chemotype identity (C), plot-level chemotype richness (CR), and the interaction between chemotype identity and plot-level chemotype richness (C x CR) on the number of stems of *T. vulgare* plants: **(a)** June 01, 2021, **(b)** June 22, 2021, **(c)** October 28, 2021, and **(d)** July 06, 2022. The axis breaks are the same for panels of the same year for illustration purposes. Significance is indicated as follows: n.s. = not significant, * P <0.05, ** P <0.01 and *** P <0.001. Degrees of freedom, Wald's Chi-square statistics, and p-values are reported in Table S2-2. Tukey *post hoc* significant differences between chemotypes are indicated with different letters.

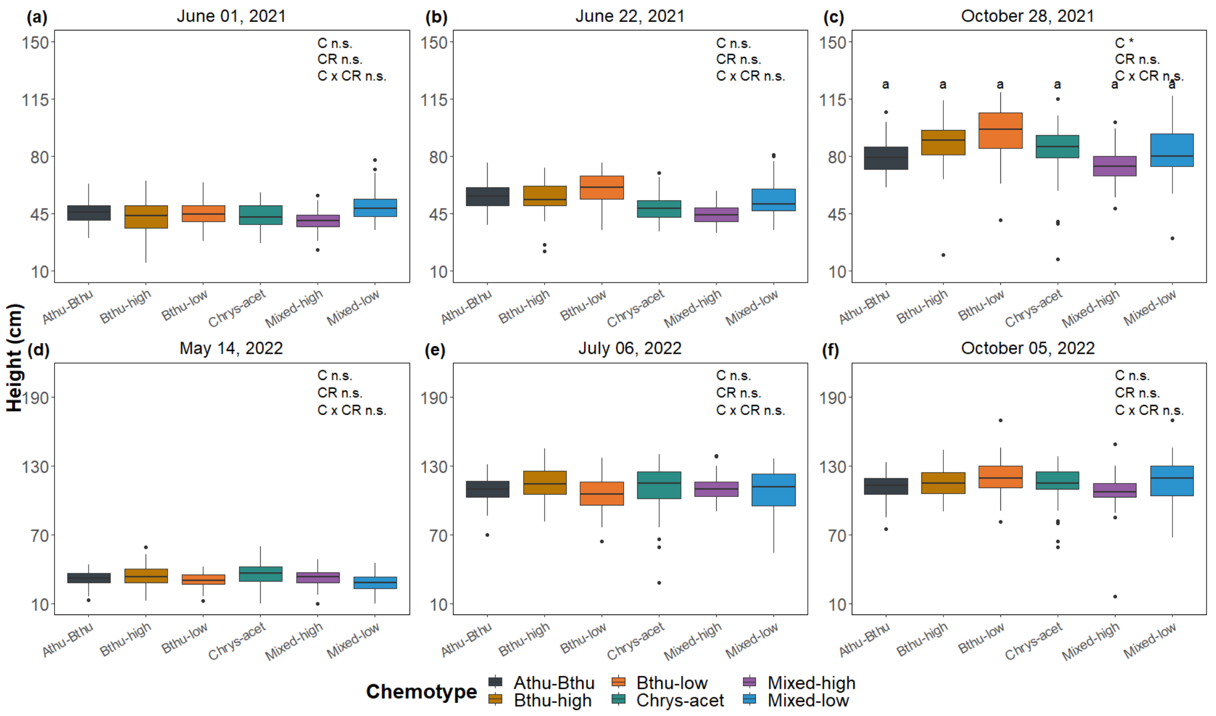


**Figure S2-2:** Effect of chemotype identity (C), plot-level chemotype richness (CR), and the interaction between chemotype identity and plot-level chemotype richness (C x CR) on height (cm) of *T. vulgare* plants: **(a)** June 01, 2021, **(b)** June 22, 2021, **(c)** October 28, 2021, **(d)** May 14, 2022, **(e)** July 06, 2022, and **(f)** October 05, 2022. The axis breaks are the same for panels of the same year for illustration purposes. Significance is indicated as follows: n.s. = not significant, * P <0.05, ** P <0.01 and *** P <0.001. Degrees of freedom, Wald's Chi-square statistics, and p-values are reported in Table S2-3. Tukey *post hoc* significant differences between chemotypes are indicated with different letters.

*2.3.2.2. Effects of daughter and plot-level chemotype richness on traits of individual plants*

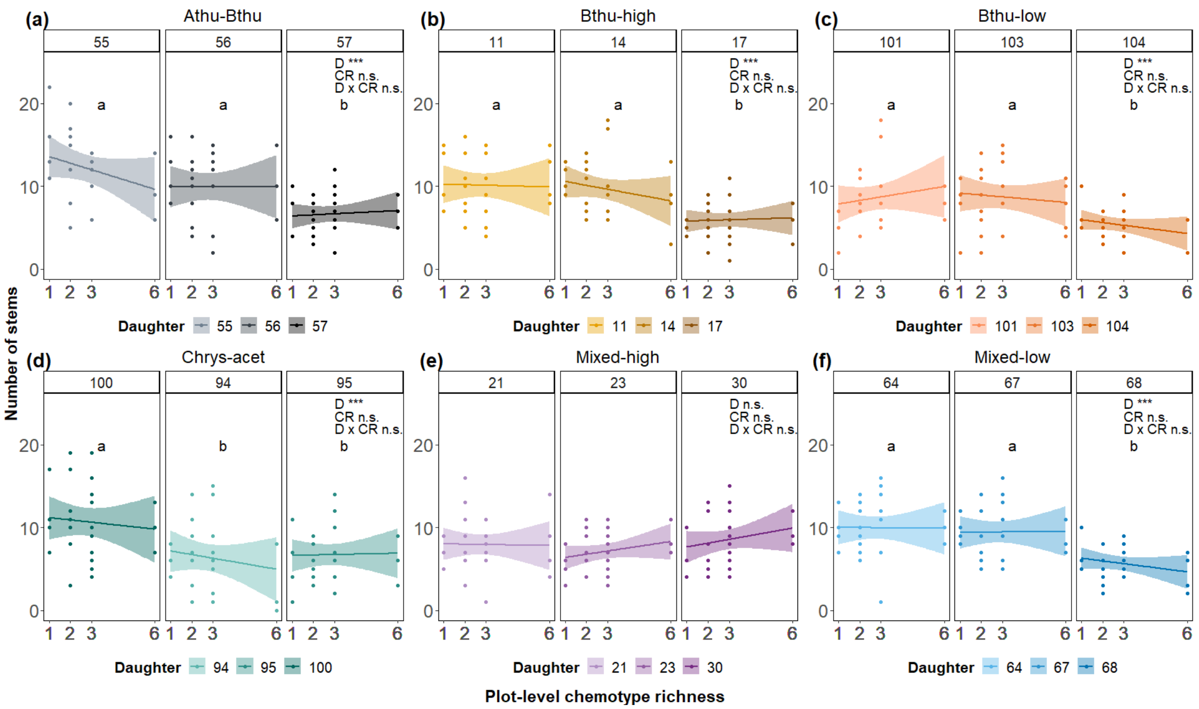


**Figure S2-3:** Effect of daughter identity (D), plot-level chemotype richness (CR), and the interaction between daughter identity and plot-level chemotype richness (D x CR) on the number of stems of *T. vulgare* plants on October 28, 2021. Each panel shows the differences across daughters within chemotype **(a)** Athu-Bthu, **(b)** Bthu-high, **(c)** Bthu-low, **(d)** Chrys-acet, **(e)** Mixed-high, and **(f)** Mixed-low. Significance is indicated as follows: n.s. = not significant, * P <0.05, ** P <0.01 and *** P <0.001. Degrees of freedom, Wald's Chi-square statistics, and p-values are reported in Table S2-6. Tukey *post hoc* significant differences between chemotypes are indicated with different letters.

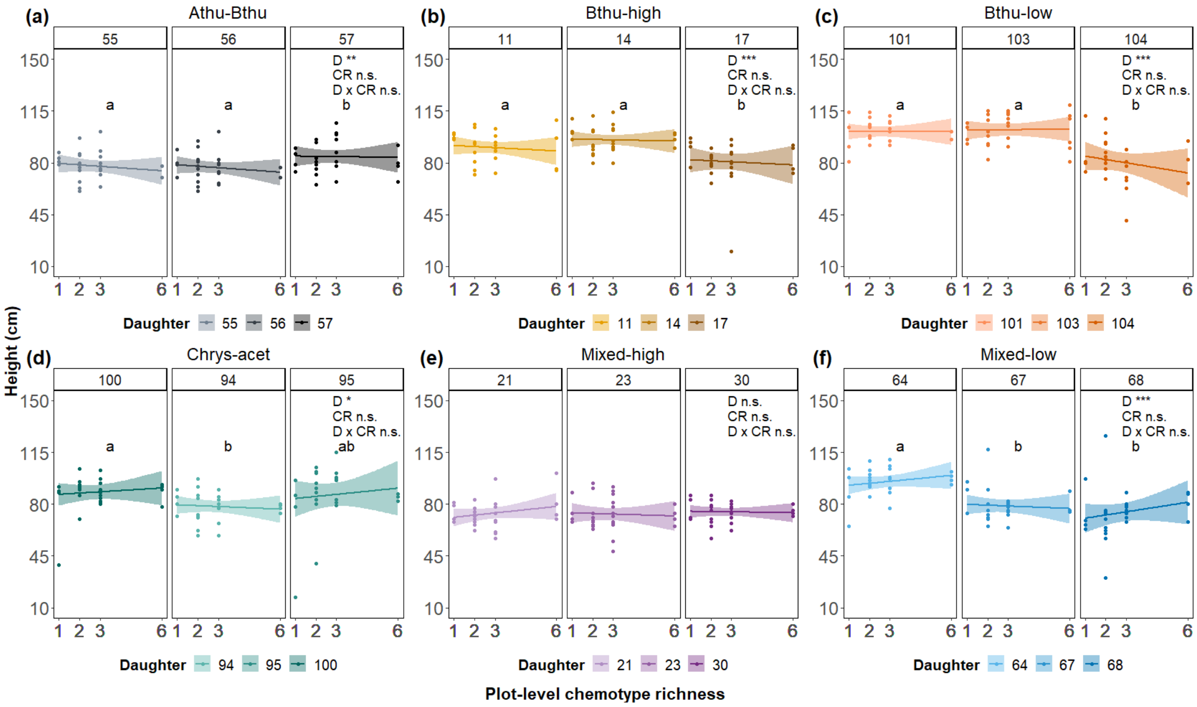


**Figure S2-4:** Effect of daughter identity (D), plot-level chemotype richness (CR), and the interaction between daughter identity and plot-level chemotype richness (D x CR) on height (cm) of *T. vulgare* plants on October 28, 2021. Each panel shows the differences across daughters within chemotype **(a)** Athu-Bthu, **(b)** Bthu-high, **(c)** Bthu-low, **(d)** Chrys-acet, **(e)** Mixed-high, and **(f)** Mixed-low. Significance is indicated as follows: n.s. = not significant, * P <0.05, ** P <0.01 and *** P <0.001. Degrees of freedom, Wald's Chi-square statistics, and p-values are reported in Table S2-6. Tukey *post hoc* significant differences between chemotypes are indicated with different letters.

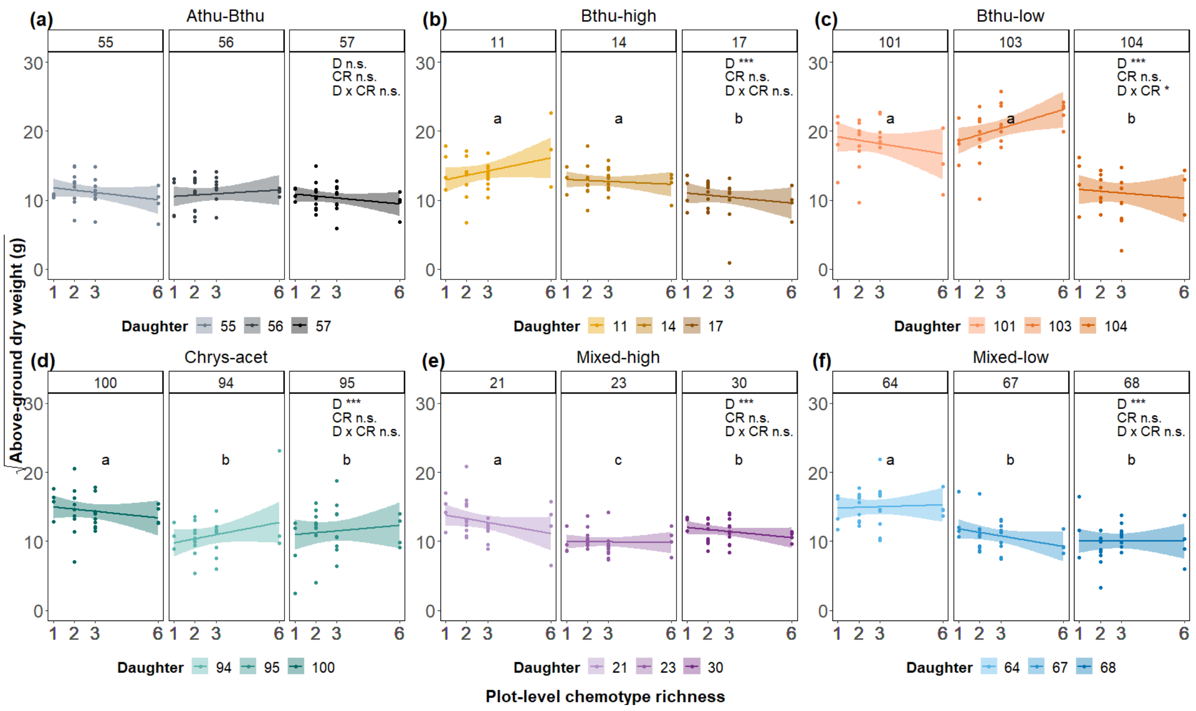


**Figure S2-5:** Effect of daughter identity (D), plot-level chemotype richness (CR), and the interaction between daughter identity and plot-level chemotype richness (D x CR) on the squared root of dry above-ground biomass (g) of *T. vulgare* plants on October 28, 2021. Each panel shows the differences across daughters within chemotype **(a)** Athu-Bthu, **(b)** Bthu-high, **(c)** Bthu-low, **(d)** Chrys-acet, **(e)** Mixed-high, and **(f)** Mixed-low. Significance is indicated as follows: n.s. = not significant, * P <0.05, ** P <0.01 and *** P <0.001. Degrees of freedom, Wald's Chi-square statistics, and p-values are reported in Table S2-6. Tukey *post hoc* significant differences between chemotypes are indicated with different letters.

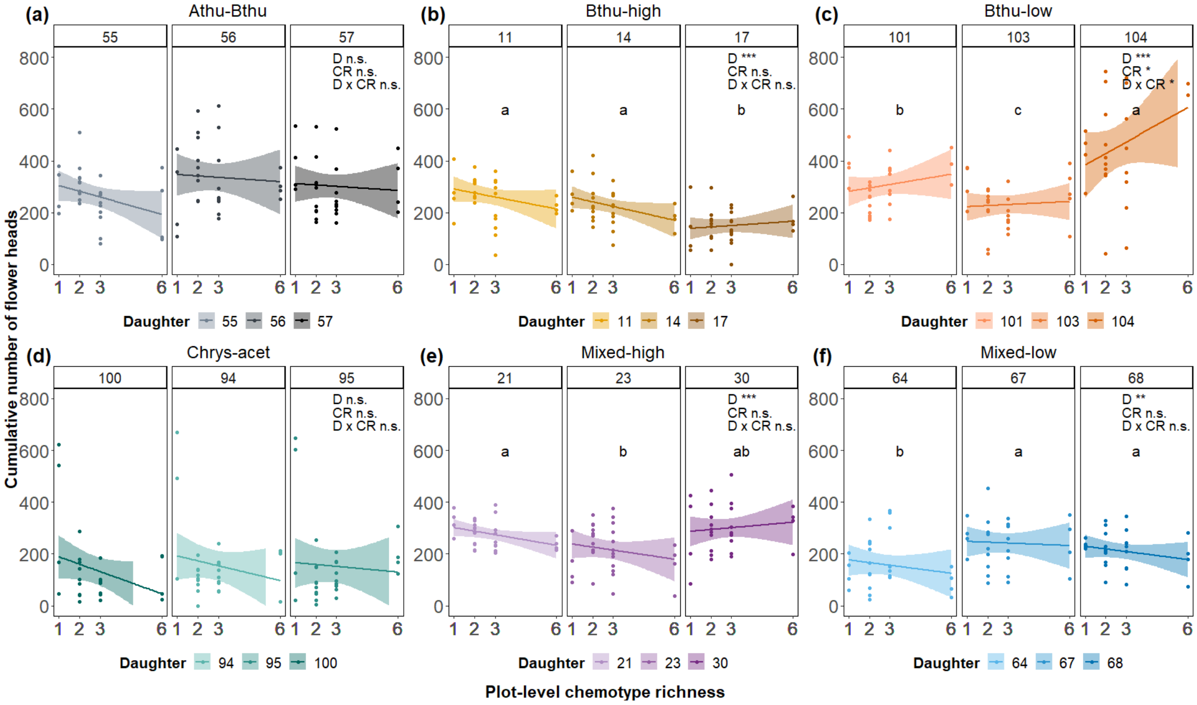


**Figure S2-6:** Effect of daughter identity (D), plot-level chemotype richness (CR), and the interaction between daughter identity and plot-level chemotype richness (D x CR) on the cumulative number of flower heads of *T. vulgare* plants in 2021*.* Each panel shows the differences across daughters within chemotype **(a)** Athu-Bthu, **(b)** Bthu-high, **(c)** Bthu-low, **(d)** Chrys-acet, **(e)** Mixed-high, and **(f)** Mixed-low. Significance is indicated as follows: n.s. = not significant, * P <0.05, ** P <0.01 and *** P <0.001. Degrees of freedom, Wald's Chi-square statistics, and p-values are reported in Table S2-6. Tukey *post hoc* significant differences between chemotypes are indicated with different letters.

*2.3.2.3. Effects of chemotype and plot-level chemotype richness on plot-level means*

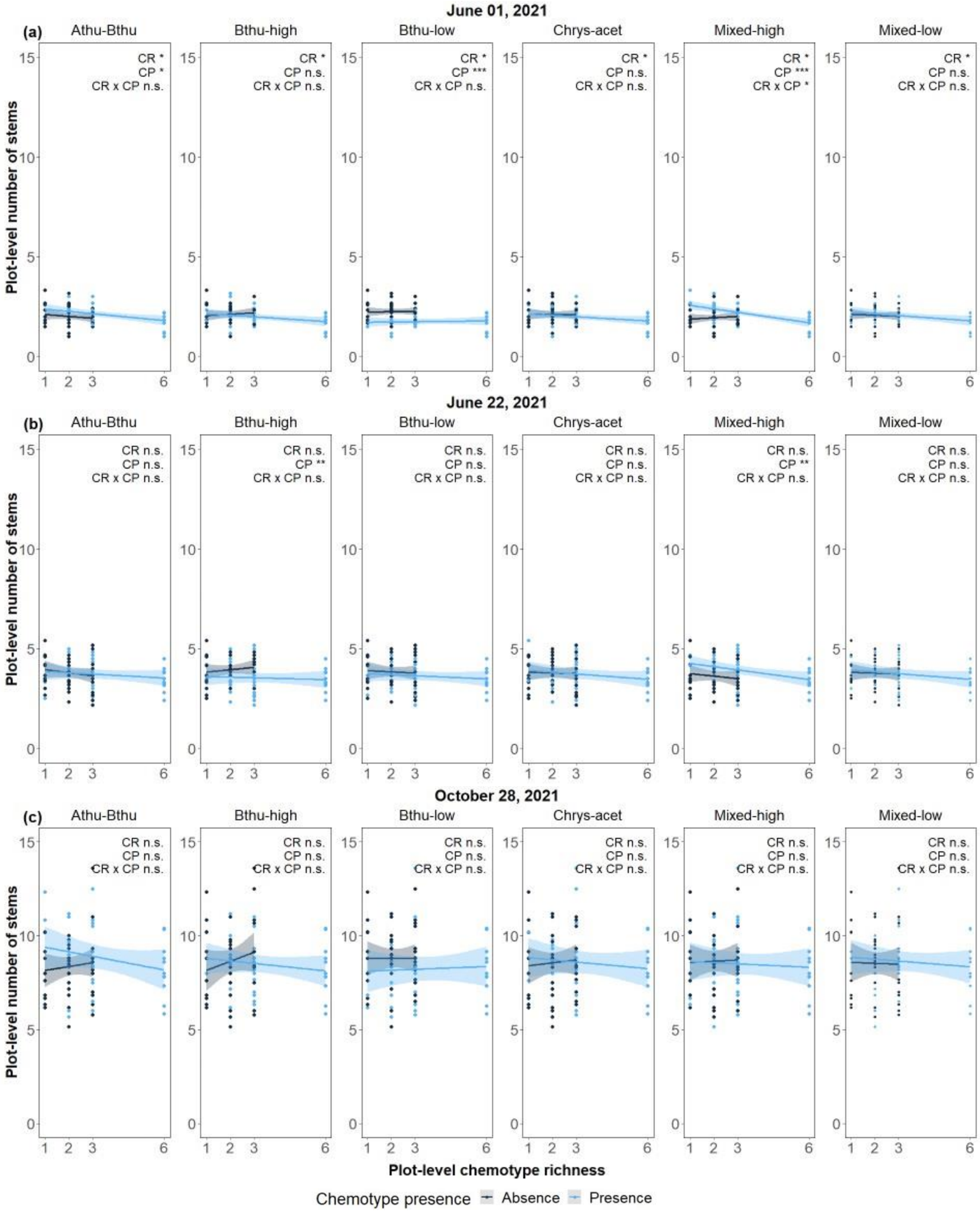


**CR =** Plot-level chemotype richness
**CP =** Chemotype presence
**CR x CP =** Plot-level chemotype richness x Chemotype presence

**Figure S2-7:** Effect of chemotype presence (CP), plot-level chemotype richness (CR), and the interaction between chemotype presence and plot-level chemotype richness (CP x CR) on plot-level number of stems of *T. vulgare* plants in 2021*:* **(a)** June 01, 2021, **(b)** June 22, 2021, and **(c)** October 28, 2021. Significance is indicated as follows: n.s. = not significant, * $P < 0.05$, ** $P < 0.01$ and *** $P < 0.001$. Degrees of freedom, F-statistics, and p-values are reported in Table S2-7.

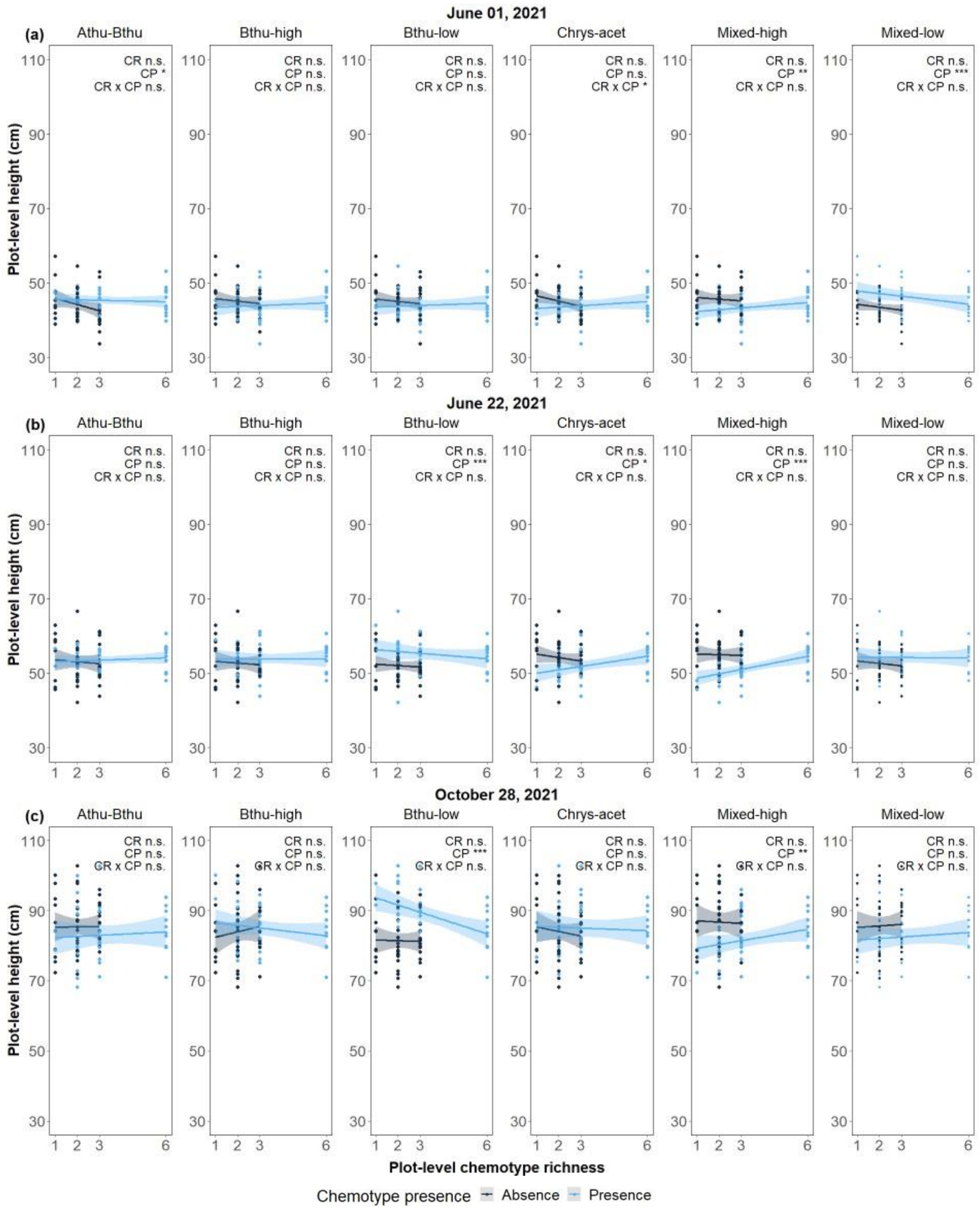


**CR =** Plot-level chemotype richness
**CP =** Chemotype presence
**CR x CP =** Plot-level chemotype richness x Chemotype presence

**Figure S2-8**: Effect of chemotype presence (CP), plot-level chemotype richness (CR), and the interaction between chemotype presence and plot-level chemotype richness (CP x CR) on plot-level height (cm) of *T. vulgare* plants in 2021*:* **(a)** June 01, 2021, **(b)** June 22, 2021, and **(c)** October 28, 2021. Significance is indicated as follows: n.s. = not significant, * P <0.05, ** P <0.01 and *** P <0.001. Degrees of freedom, F-statistics, and p-values are reported in Table S2-8.

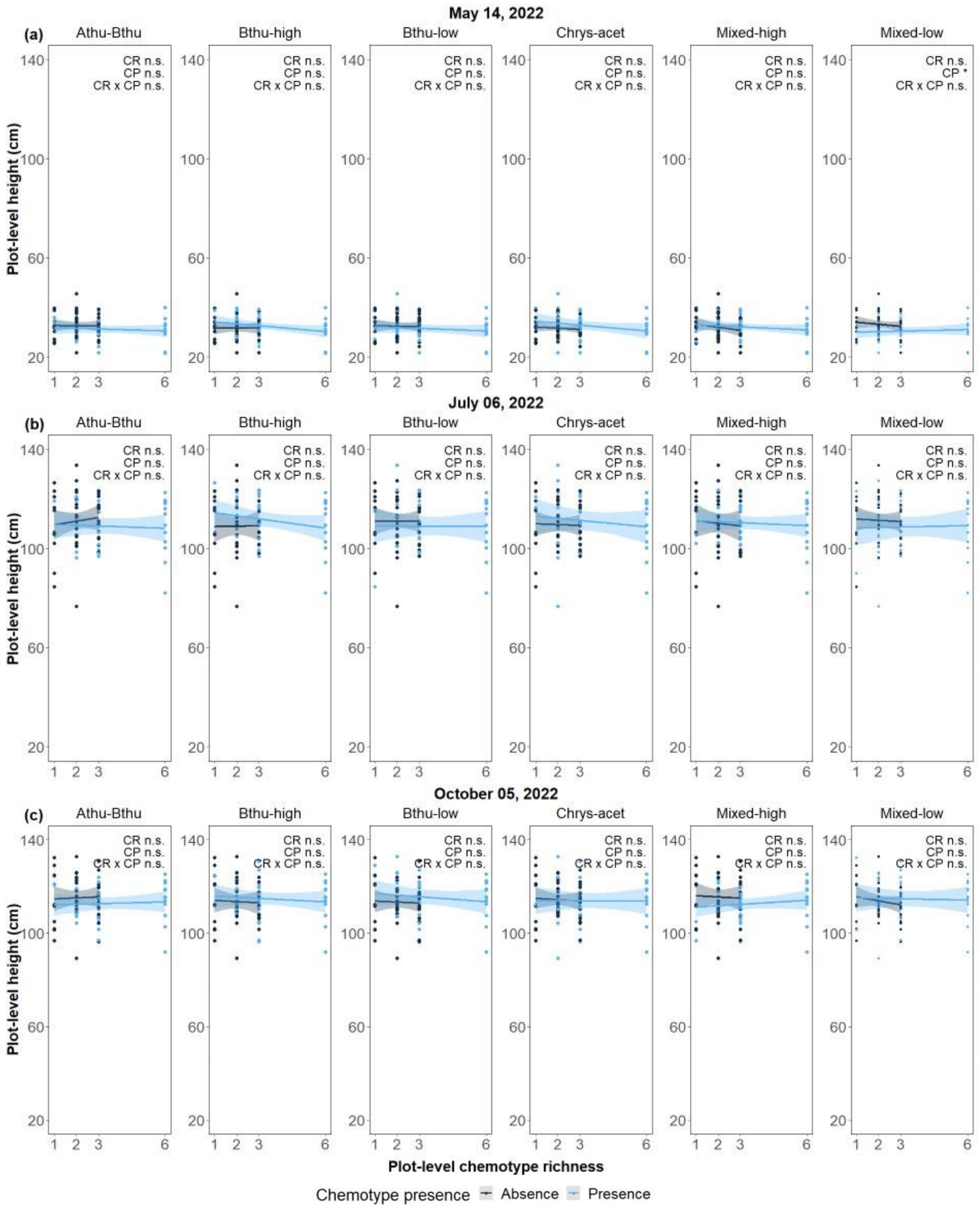


**CR =** Plot-level chemotype richness
**CP =** Chemotype presence
**CR x CP =** Plot-level chemotype richness x Chemotype presence

**Figure S2-9:** Effect of chemotype presence (CP), plot-level chemotype richness (CR), and the interaction between chemotype presence and plot-level chemotype richness (CP x CR) on plot-level height (cm) of *T. vulgare* plants in 2022: **(a)** May 14, 2022, **(b)** July 06, 2022, and **(c)** October 05, 2022. Significance is indicated as follows: n.s. = not significant, * $P < 0.05$, ** $P < 0.01$ and *** $P < 0.001$. Degrees of freedom, F-statistics, and p-values are reported in Table S2-8.

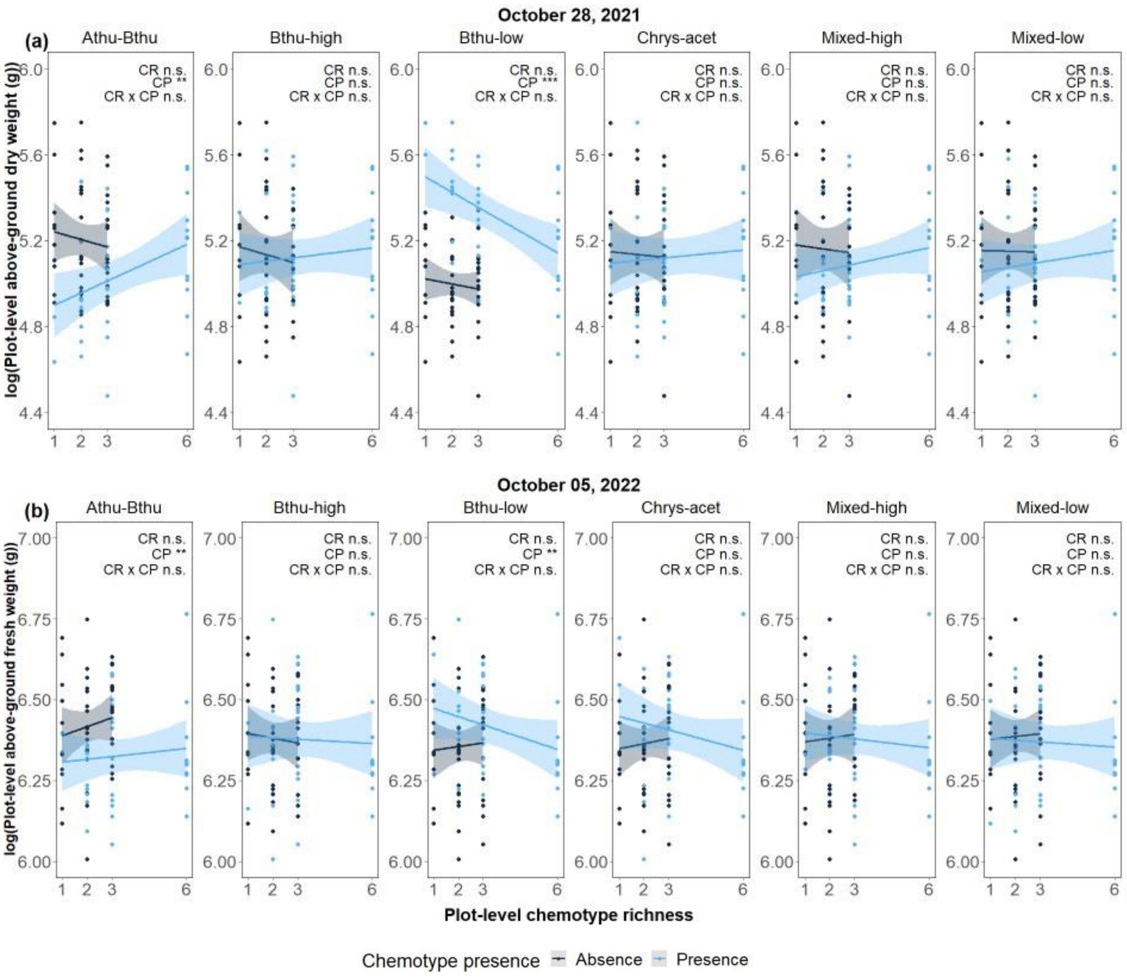


**CR =** Plot-level chemotype richness
**CP =** Chemotype presence
**CR x CP =** Plot-level chemotype richness x Chemotype presence

**Figure S2-10:** Effect of chemotype presence (CP), plot-level chemotype richness (CR), and the interaction between chemotype presence and plot-level chemotype richness (CP x CR) on **(a)** the logarithm of the plot-level above-ground dry weight (g) and **(b)** above-ground fresh weight (g) of *T. vulgare* plants. Significance is indicated as follows: n.s. = not significant, * P <0.05, ** P <0.01 and *** P <0.001. Degrees of freedom, F-statistics, and p-values are reported in Table S2-9.

*2.3.2.4. Overyielding indexes*

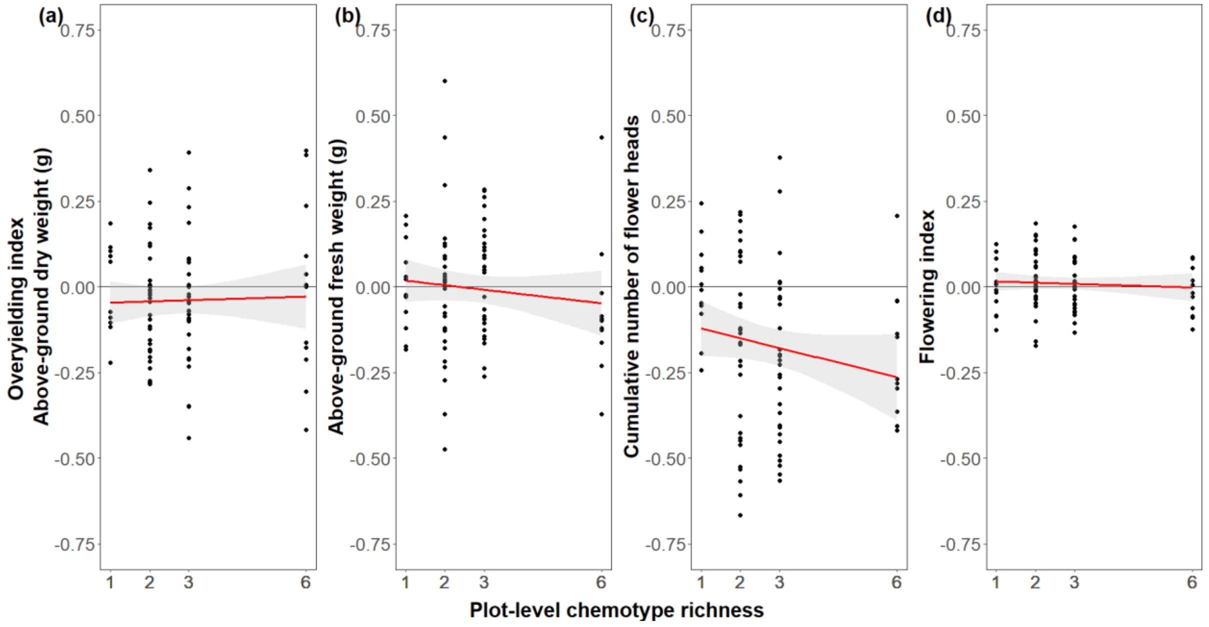


**Figure S2-11:** Effects of plot-level chemotype richness on overyielding indexes calculated for plant traits of *T. vulgare* plants: **(a)** above-ground dry weight (g) in 2021, **(b)** above-ground fresh weight (g) in 2022, **(c)** cumulative number of flower heads, and **(d)** flowering index. Degrees of freedom, Wald's Chi-square statistics, and p-values are reported in Table S2-12.

*2.3.2.5. Headspace VOC collection*

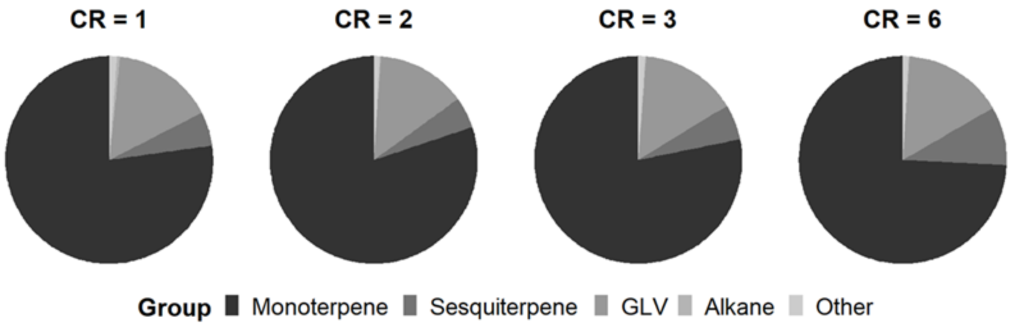


**Figure S2-12:** Headspace VOC major classes (green leaf volatiles (GLV), monoterpenes, sesquiterpenes, alkanes, and others) across plot-level chemotype richness (CR).
