## Supplementary material for "Intraspecific chemical variation of *Tanacetum vulgare* affects plant growth and reproductive traits in field plant communities": Supplmentary Methods

### Supplementary Information 1 (S1): Supplementary Methods

### Table of content

**1.1. Detailed description of methods**

*1.1.1. Chemotypic characterization of T. vulgare lines and biological replicates*

*1.1.2. Propagation of plant material for the field experiment*

*1.1.3. Plant morphological trait measurements*

*1.1.4. Headspace VOC collection*

*1.1.5. Effects of daughter and plot-level chemotype richness on traits of individual plants*

**1.2. Design of the field experiment**

*1.2.1. Leaf terpenoid composition of chemotypes*

*1.2.2. Assignment of plants to plots*

*1.2.3. Stem phenology*

**1.3. Details of the statistical analyses**

*1.3.1. Variables used in the analyses*

*1.3.2. Models of effects on plant traits*

Plant-level traits

1.3.2.1. Effects of chemotype, plot-level chemotype richness, and the interaction between chemotype and plot-level chemotype richness on plant-level traits

1.3.2.2. Effects of daughter, plot-level chemotype richness, and the interaction between daughter identity and plot-level chemotype richness on plant-level traits

Plot-level traits

1.3.2.3. Effects of chemotype presence, plot-level chemotype richness, and the interaction between chemotype presence and plot-level chemotype richness) on plot-level traits

*1.3.3. Models of effects on overyielding indexes*

1.3.3.1. Effects of plot-level chemotype richness on the overyielding indexes

*1.3.4. Models of chemodiversity*

1.3.4.1. Effects of plot-level chemotype richness on the plot-level theoretical leaf chemodiversity metrics

1.3.4.2. Effects of plot-level chemotype richness on the plot-level realized volatile chemodiversity metrics

### 1.1. Detailed description of methods

#### *1.1.1. Chemotypic characterization of T. vulgare lines and biological replicates*

We used six *T. vulgare* chemotypes that were also used in Neuhaus-Harr *et al.* (2023). Based on an unsupervised hierarchical k-means clustering calculated on a dissimilarity matrix of the terpenoid profile obtained from hexane extraction of leaves of 27 *T. vulgare* plants found in Jena, Germany (50.93°N, 11.58°E), we randomly chose two plants from each of 6 of the resulting 7 clusters (hereafter 'mother' plant) and mass-sown their seeds. We selected ten healthy seedlings from each mother plant (120 plants, hereafter 'daughter' plant) and chemotyped those plants. Based on another unsupervised hierarchical clustering calculated on the terpenoid profile from hexane extraction of those 120 daughter plants in 2021, we obtained 6 clusters. We chose three daughter plants from the same mother for each cluster, resulting in 6 chemotypes and 18 daughter plants.

#### *1.1.2. Propagation of plant material for the field experiment*

Stem parts from the 18 selected daughters were trimmed and used to produce 27-40 cuttings per daughter. Individual cuttings were cut 1-2 cm below and 4-5 cm above a leaf node. The leaf size was reduced by 60% to reduce evapotranspiration and improve the survival rate by shortening and trimming the leaflet. Stem cuttings were placed into seedling trays with soaked potting substrate and protected with a clear plastic propagation cover. After three weeks and root and shoot formation, the cover was gradually opened to allow the cuttings to acclimate. As the cuttings grew, they were transplanted into 10 cm pots and later into 17 cm pots before transplanting in the field site in May 2021. Potted plants were automatically watered from the bottom and fertilized as needed with Universol Blue fertilizer ® (18%N - 11%P - 18%K; ICL Deutschland, Nordhorn, Germany). During propagation and growth, plants were not treated with chemical insecticides or fungicides, but we applied biological control when needed. At the end of the growing period, there were multiple cloned plants from each of the 18 daughters. Cloned plants from each of the three daughters of each chemotype were used in the experimental design.

#### *1.1.3. Plant morphological trait measurements*

We measured six morphological plant traits for individual plants, including four growth traits (the number of stems per plant, the height of the highest stem, and above-ground fresh and dry weight) and two reproductive traits (the cumulative number of flower heads and the flowering index).

Three times in 2021 (June 01, June 22, and October 28), we counted the number of stems larger than 5 centimeters and measured plant height as the distance between the ground and the longest stem without stretching out leaves.

As a measure of reproductive success, we counted and cut off all ripe flower heads weekly during the 2021 season (from June 01 to October 28). Flower heads were cut off to avoid re-seeding plots with new plants. We thus refer to the total number of flower heads collected as the cumulative number of flower heads. At the end of the season (October 28, 2021), we obtained the above-ground dry weight of each plant; plants were cut off 5 cm above the ground, placed in single paper bags, and dried for 72 hours at 60°C in a drying room (Thermo-Scientific) of the University of Jena facilities.

In 2022, we counted the number of stems once (July 06) and measured plant height three times (May 14, July 06, and October 05). The phenological state of each stem of each plant was determined once (on July 06) using numerical categories: [1] No inflorescence; [2] Inflorescence visible but closed; [3] Open inflorescence; [4] Beginning seed set (unripe); [5] Ripe seeds; [6] Mature seeds turning brown (Fig. S1-1). To analyze phenology at the plant level, we calculated a flowering index for each plant by dividing the sum of all stem-level phenological stages by the number of stems (between 1 = all stems without inflorescence and 6 = all stems with mature seeds). We averaged plant-level flower indices per plot to obtain a measurement at the plot level.

At the end of the second season, all plants were again cut off 5 cm above ground but immediately weighted in the field to obtain fresh weight (October 05, 2022). From this, we calculated both plant-level and plot-level weights.

*1.1.4. Headspace VOC collection*

Headspace volatile organic compound (VOC) emissions were collected at the plot level (including all six individual plants) between May 12 and 14, 2022, using a closed-push-pull system for 2 hours (Fig. 1f-g). Weeds were removed by hand from the plots ten days before the measurements to reduce the collection of volatiles from plants different from tansy. In addition, one extra new plot was prepared by thoroughly cleaning out the plant material ten days before and used as a volatile background profile (red plot in Fig. 1d).

A squared bag made from Polyethylene terephthalate PET (110 cm x 110 cm x 40 cm) foil was installed for VOC collections, covering the whole plot. Air entered the system after passing through an activated charcoal filter at a flow rate of 1.5 L/min and was pumped out through a trap at 1 L/min. The trap contained 25 mg of Poropak-Q absorbent (Volatile Collection Trap (VCT) LLC, USA) (Fig. 1f). VOCs were collected over a period of three days with three rounds of ten plots per day (two entire blocks + two background plots = 30 plots per day). Plots were assigned beforehand to ensure each plot-level chemotype richness level were included in each round.

After VOC collection, the traps were eluted with 200µl dichloromethane containing nonyl-acetate as an internal standard (Sigma-Aldrich, 10 ng x $\mu l^{-1}$). VOCs were analyzed using a Hewlett-Packard model 6890 gas chromatograph coupled to a Hewlett-Packard model 5973 mass spectrometer (GC-MS). The GC was operated with helium as carrier gas at 1 ml/min, split-less injection (injection temperature: 220 °C, injection volume: 1 µl), a DB-5MS column (30 m × 0.25 mm, 0.25 µm film, J & W Scientific, Folsom, CA, USA), and a temperature program from 40 °C (2 min hold) to 350 °C (2 min hold) with a first gradient of 7 °C $min^{-1}$ to 155 °C, and a second gradient of 60 °C $min^{-1}$ to 300 °C. The GC was coupled to a single quadrupole mass spectrometer (Hewlett-Packard model 5973) operated in electron impact ionization mode. The transfer line temperature was 270 °C, the ionization potential was 70 eV, and a scan range of *m/z* 40–350 was employed.

VOCs were identified by comparing retention times and mass spectra with commercial standards (when available) and by comparison to the mass spectral libraries Wiley 275 (Wiley and National Institute of Standards) and NIST05a (National Institute of Standards and Technology, Gaithersburg, MD). We verified the VOC identity by calculating Kovats retention indices. The qualitative and quantitative analysis of the volatile data is beyond the scope of this study; here, we report only on volatile terpenoid diversity metrics at the plot level.

*1.1.5. Effects of daughter and plot-level chemotype richness on traits of individual plants*

We performed separate analyses per chemotype with Daughter, plot-level chemotype richness, and their interaction as fixed factors, and Plot ID nested in Block ID as a random factor. For count data, we used a GLMM with Poisson distribution and Nelder Mead optimizer and, for the other assessed variables, LMM models.

In a final analysis, we directly compared daughters' performance, using daughter ID (with 18 levels) and plot-level chemotype richness as independent variables, and plot ID and block ID as random factors.

### 1.2. Design of the field experiment

#### *1.2.1. Leaf terpenoid composition of chemotypes*

**Table S1-1:** Leaf terpenoid composition (nmol/g) of chemotypes of *T. vulgare* plants. The table is partitioned into three sections showing different terpenoids in alphabetic order. Each section lists the chemotypes tested (6), the three daughter plants chosen for each chemotype (18), and the concentration of each terpenoid found in each daughter after hexane extraction of leaf material. Chemotypes are named after their dominant compounds, and numbers identify daughters. Plant chemotyping and chemotype selection are described in more detail in Neuhaus-Harr *et al.* (2023).

| Chemotype | Daughter | 1-terpin-4-nyl acetate | α-chrysanthenyl acetate | α-farnesene | α-pinene | α-terpinene | α-thujene | α-thujone | artemisia-β-acetate | artemisia alcohol | artemisia ketone | β-carvyl acetate | β-caryophyllene | β-ocimene | β-pinene | β-thujone | borneol | camphene | camphor |
|---|---|---|---|---|---|---|---|---|---|---|---|---|---|---|---|---|---|---|---|
| Athu-Bthu | 55 | 0.01 | 0.00 | 0.00 | 0.17 | 0.01 | 0.00 | 27.66 | 0.05 | 0.00 | 0.00 | 0.00 | 0.00 | 0.05 | 0.85 | 28.25 | 0.00 | 0.00 | 0.01 |
| Athu-Bthu | 56 | 0.01 | 0.07 | 0.00 | 0.34 | 0.00 | 0.00 | 24.05 | 0.13 | 0.52 | 0.00 | 0.00 | 0.00 | 0.01 | 0.11 | 24.67 | 0.30 | 0.00 | 0.01 |
| Athu-Bthu | 57 | 0.02 | 0.12 | 0.00 | 0.34 | 0.00 | 0.00 | 22.70 | 0.32 | 0.62 | 0.60 | 0.00 | 0.00 | 0.02 | 0.10 | 23.10 | 0.26 | 0.00 | 0.01 |
| Bthu-high | 11 | 0.02 | 0.08 | 0.70 | 0.29 | 0.00 | 0.00 | 0.20 | 0.64 | 0.69 | 0.00 | 0.06 | 0.00 | 0.05 | 3.28 | 39.76 | 0.43 | 0.00 | 0.01 |
| Bthu-high | 14 | 0.01 | 0.16 | 0.00 | 0.37 | 0.00 | 0.00 | 0.19 | 0.00 | 0.00 | 0.05 | 0.00 | 0.00 | 0.00 | 0.24 | 40.37 | 0.54 | 0.00 | 0.01 |
| Bthu-high | 17 | 0.02 | 0.00 | 0.00 | 0.33 | 0.01 | 0.11 | 0.20 | 0.00 | 0.94 | 0.00 | 0.20 | 0.00 | 0.04 | 0.45 | 42.55 | 0.00 | 0.00 | 0.01 |
| Bthu-low | 101 | 0.02 | 0.00 | 0.00 | 0.20 | 0.01 | 0.00 | 0.00 | 0.00 | 0.00 | 0.00 | 0.00 | 0.00 | 0.02 | 0.00 | 29.33 | 0.00 | 0.02 | 0.01 |
| Bthu-low | 103 | 0.02 | 0.10 | 0.00 | 0.22 | 0.01 | 0.00 | 0.00 | 0.00 | 0.00 | 0.40 | 0.00 | 0.00 | 0.03 | 5.59 | 23.29 | 0.41 | 0.01 | 0.01 |
| Bthu-low | 104 | 0.01 | 0.12 | 0.00 | 0.18 | 0.01 | 0.11 | 0.18 | 0.00 | 0.00 | 0.05 | 0.00 | 0.00 | 0.02 | 4.66 | 32.59 | 0.41 | 0.01 | 0.01 |
| Chrys-acet | 94 | 0.01 | 0.05 | 0.00 | 2.09 | 0.00 | 0.00 | 0.00 | 0.03 | 0.00 | 0.25 | 0.00 | 0.00 | 0.02 | 0.07 | 0.00 | 0.18 | 0.42 | 0.02 |
| Chrys-acet | 95 | 0.01 | 0.02 | 0.00 | 2.51 | 0.00 | 0.07 | 0.00 | 0.03 | 0.00 | 0.29 | 0.00 | 0.00 | 0.07 | 0.49 | 0.00 | 0.31 | 0.40 | 0.79 |
| Chrys-acet | 100 | 0.01 | 0.04 | 0.00 | 2.86 | 0.01 | 0.05 | 0.00 | 0.08 | 0.00 | 0.15 | 0.00 | 0.00 | 0.02 | 0.12 | 0.00 | 0.00 | 0.51 | 0.02 |
| Mixed-high | 21 | 0.03 | 0.00 | 0.00 | 2.87 | 0.01 | 0.33 | 0.00 | 0.00 | 0.00 | 0.00 | 0.11 | 0.00 | 0.18 | 0.83 | 0.00 | 7.59 | 2.32 | 0.01 |
| Mixed-high | 23 | 0.01 | 0.00 | 0.00 | 2.30 | 0.01 | 0.16 | 0.00 | 0.20 | 0.78 | 0.00 | 0.00 | 0.00 | 0.08 | 0.73 | 0.00 | 7.63 | 1.80 | 0.01 |
| Mixed-high | 30 | 0.02 | 0.00 | 0.00 | 3.58 | 0.00 | 0.36 | 0.00 | 0.00 | 0.66 | 0.00 | 0.14 | 0.00 | 0.03 | 1.00 | 0.00 | 7.96 | 2.91 | 0.01 |
| Mixed-low | 64 | 0.03 | 0.12 | 0.00 | 6.30 | 0.01 | 0.00 | 0.00 | 0.00 | 0.00 | 0.00 | 0.12 | 0.00 | 0.02 | 2.63 | 0.00 | 0.37 | 0.06 | 0.02 |
| Mixed-low | 67 | 2.80 | 0.00 | 0.00 | 5.35 | 0.01 | 0.07 | 0.00 | 0.09 | 0.00 | 0.00 | 0.00 | 0.75 | 1.55 | 1.04 | 0.00 | 3.63 | 1.50 | 0.05 |
| Mixed-low | 68 | 0.02 | 0.00 | 0.00 | 4.96 | 0.00 | 0.16 | 0.00 | 0.00 | 0.00 | 0.72 | 0.00 | 0.09 | 0.01 | 3.25 | 0.00 | 0.00 | 0.01 | 0.01 |

**Table S1-1 *(continued)***: *This section shows the results for the second part of Table S1-1, partitioned by terpenoid.*

| Chemotype | Daughter | chrysanthenyl acetate | cis-chrysanthenol | cis-miroxide | cis-sabinene hydrate | cis-verbanol acetate | eucalyptol | eugenol | γ-terpinene | limonene | methylisoborneol | mustakone | myrcene | p-cymene | pinocarvone | sabinene | santolina triene | trans-chrysanthenol | trans-sabinene hydrate |
|---|---|---|---|---|---|---|---|---|---|---|---|---|---|---|---|---|---|---|---|
| Athu-Bthu | 55 | 0.00 | 0.00 | 0.02 | 0.04 | 0.00 | 0.00 | 0.00 | 0.05 | 0.00 | 0.00 | 0.00 | 0.10 | 0.51 | 1.09 | 0.85 | 0.01 | 0.00 | 0.10 |
| Athu-Bthu | 56 | 0.00 | 0.00 | 0.01 | 0.00 | 0.00 | 0.01 | 0.02 | 0.06 | 0.00 | 0.00 | 0.00 | 0.00 | 0.32 | 0.01 | 0.51 | 0.00 | 0.09 | 0.00 |
| Athu-Bthu | 57 | 0.00 | 0.02 | 0.01 | 0.07 | 0.00 | 0.02 | 0.01 | 0.03 | 0.00 | 0.05 | 0.00 | 0.00 | 0.24 | 0.46 | 0.42 | 0.01 | 0.00 | 0.00 |
| Bthu-high | 11 | 0.00 | 0.01 | 0.01 | 0.08 | 0.00 | 0.02 | 0.00 | 0.01 | 0.09 | 0.00 | 0.00 | 0.00 | 0.33 | 0.06 | 3.22 | 0.00 | 0.00 | 0.00 |
| Bthu-high | 14 | 0.00 | 0.00 | 0.02 | 0.07 | 0.00 | 0.00 | 0.00 | 0.04 | 0.00 | 0.00 | 0.00 | 0.06 | 0.21 | 0.06 | 1.60 | 0.00 | 0.03 | 0.00 |
| Bthu-high | 17 | 0.00 | 0.01 | 0.05 | 0.07 | 0.00 | 0.02 | 0.04 | 0.01 | 0.00 | 0.00 | 0.00 | 0.00 | 0.43 | 0.00 | 6.52 | 0.00 | 0.00 | 0.00 |
| Bthu-low | 101 | 0.00 | 0.01 | 0.02 | 0.10 | 0.00 | 0.12 | 0.03 | 0.02 | 0.00 | 0.00 | 0.00 | 0.06 | 0.43 | 0.03 | 0.04 | 0.01 | 0.00 | 0.00 |
| Bthu-low | 103 | 0.00 | 0.07 | 0.01 | 0.00 | 0.00 | 0.01 | 0.00 | 0.02 | 0.00 | 0.00 | 0.00 | 0.00 | 0.66 | 0.01 | 5.59 | 0.01 | 0.00 | 0.00 |
| Bthu-low | 104 | 0.00 | 0.00 | 0.01 | 0.06 | 0.00 | 0.24 | 0.03 | 0.01 | 0.00 | 0.02 | 0.00 | 0.10 | 0.45 | 0.76 | 4.68 | 0.01 | 0.05 | 0.12 |
| Chrys-acet | 94 | 48.13 | 2.32 | 0.02 | 0.08 | 0.00 | 0.78 | 0.05 | 0.85 | 0.00 | 0.58 | 0.00 | 0.02 | 0.84 | 0.01 | 0.73 | 0.01 | 0.00 | 0.10 |
| Chrys-acet | 95 | 45.84 | 1.68 | 0.01 | 0.04 | 0.00 | 0.00 | 0.02 | 0.01 | 0.00 | 0.00 | 0.00 | 0.00 | 0.30 | 0.01 | 0.49 | 0.01 | 0.02 | 0.07 |
| Chrys-acet | 100 | 46.13 | 1.95 | 0.01 | 0.08 | 0.00 | 0.00 | 0.02 | 0.70 | 0.00 | 0.00 | 0.00 | 0.02 | 0.75 | 0.00 | 0.80 | 0.00 | 0.00 | 0.04 |
| Mixed-high | 21 | 0.00 | 0.00 | 0.03 | 16.90 | 0.00 | 0.00 | 0.00 | 0.18 | 0.00 | 7.64 | 0.00 | 0.00 | 0.09 | 0.01 | 5.70 | 0.00 | 0.00 | 0.00 |
| Mixed-high | 23 | 0.00 | 0.08 | 0.02 | 17.07 | 0.00 | 0.00 | 0.00 | 0.11 | 0.00 | 7.84 | 3.19 | 0.03 | 0.00 | 0.07 | 3.95 | 0.00 | 0.00 | 1.29 |
| Mixed-high | 30 | 0.00 | 0.01 | 0.01 | 16.57 | 0.00 | 0.04 | 0.01 | 0.45 | 0.00 | 8.03 | 0.00 | 0.03 | 0.00 | 0.00 | 6.41 | 0.00 | 0.00 | 0.10 |
| Mixed-low | 64 | 0.00 | 0.43 | 0.02 | 0.03 | 0.00 | 0.00 | 0.03 | 1.13 | 0.00 | 0.00 | 0.00 | 0.00 | 1.18 | 0.01 | 11.62 | 0.01 | 0.06 | 0.03 |
| Mixed-low | 67 | 0.15 | 0.03 | 0.02 | 0.07 | 2.56 | 0.04 | 0.16 | 2.33 | 0.00 | 1.07 | 0.00 | 0.03 | 0.00 | 0.01 | 6.45 | 0.00 | 0.00 | 0.18 |
| Mixed-low | 68 | 0.00 | 0.01 | 0.02 | 0.42 | 0.00 | 0.03 | 0.00 | 0.51 | 0.00 | 0.00 | 0.00 | 0.11 | 0.04 | 0.01 | 13.04 | 0.01 | 0.00 | 0.26 |

**Table S1-1 *(continued)***: *This section shows the results for the third and final part of Table S1-1, partitioned by terpenoid.*

| Chemotype | Daughter | trans-sabinyl acetate | Unknown terpenoid | Unknown sesquiterpene 1 | Unknown sesquiterpene 2 | Unknown sesquiterpene 3 | Unknown sesquiterpene 4 | Unknown sesquiterpene 5 | yomogi alcohol |
|---|---|---|---|---|---|---|---|---|---|
| Athu-Bthu | 55 | 0.00 | 0.01 | 0.00 | 0.00 | 0.07 | 0.00 | 0.00 | 0.03 |
| Athu-Bthu | 56 | 0.00 | 0.02 | 0.09 | 0.06 | 0.09 | 0.00 | 0.01 | 0.08 |
| Athu-Bthu | 57 | 0.00 | 0.01 | 0.00 | 0.13 | 0.22 | 0.00 | 0.00 | 0.00 |
| Bthu-high | 11 | 0.00 | 0.02 | 0.00 | 0.00 | 0.00 | 2.26 | 0.00 | 0.06 |
| Bthu-high | 14 | 0.00 | 0.01 | 0.00 | 0.16 | 0.15 | 0.00 | 0.00 | 0.09 |
| Bthu-high | 17 | 0.01 | 0.01 | 0.03 | 0.00 | 0.00 | 0.00 | 0.00 | 0.08 |
| Bthu-low | 101 | 0.00 | 0.01 | 0.09 | 0.05 | 0.17 | 0.00 | 0.00 | 0.16 |
| Bthu-low | 103 | 0.00 | 0.01 | 0.00 | 0.00 | 0.00 | 0.00 | 0.00 | 0.13 |
| Bthu-low | 104 | 0.00 | 0.01 | 0.00 | 0.00 | 0.10 | 0.00 | 0.00 | 0.08 |
| Chrys-acet | 94 | 0.00 | 0.81 | 0.02 | 0.00 | 0.00 | 0.00 | 0.13 | 0.05 |
| Chrys-acet | 95 | 0.00 | 0.79 | 0.12 | 0.05 | 0.06 | 0.00 | 0.00 | 0.00 |
| Chrys-acet | 100 | 0.00 | 1.07 | 0.00 | 0.01 | 0.09 | 0.00 | 0.00 | 0.09 |
| Mixed-high | 21 | 0.01 | 6.53 | 0.05 | 0.00 | 0.00 | 0.00 | 0.00 | 0.06 |
| Mixed-high | 23 | 0.00 | 6.92 | 0.00 | 0.06 | 0.25 | 0.00 | 0.44 | 0.06 |
| Mixed-high | 30 | 0.00 | 9.19 | 0.00 | 0.00 | 0.00 | 0.00 | 0.00 | 0.08 |
| Mixed-low | 64 | 0.00 | 0.02 | 0.06 | 0.09 | 0.19 | 0.00 | 0.00 | 0.10 |
| Mixed-low | 67 | 0.00 | 0.00 | 0.00 | 0.00 | 0.00 | 0.00 | 0.86 | 0.09 |
| Mixed-low | 68 | 0.00 | 0.02 | 0.03 | 0.00 | 0.18 | 0.00 | 0.00 | 0.00 |

#### *1.2.2. Assignment of plants to plots*

**Table S1-2:** Assignment of plants to plots. The table is partitioned into 11 sections, showing the plant identity in order. The field experiment contained 84 plots: 12, 30, 30, and 12 replicate plots for plot-level chemotype richness levels 1, 2, 3, and 6, respectively. Plots were distributed equally in six randomized blocks. Each block consisted of 14 plots: two plots of chemotype richness level 1, five plots of chemotype richness level 2, five plots of chemotype richness level 3, and two plots with chemotype richness level 6. Each plot consisted of 6 plants, which position within the plot was randomly assigned (Plot position), and which chemotype and daughter identities are listed here.

| Block | Plot | Plant | Plot position | Plot-level chemotype richness | Chemotype | Daughter |
|---|---|---|---|---|---|---|
| 1 | 1 | 1 | 1 | 3 | Athu-Bthu | 57 |
| 1 | 1 | 2 | 2 | 3 | Chrys-acet | 100 |
| 1 | 1 | 3 | 3 | 3 | Chrys-acet | 94 |
| 1 | 1 | 4 | 4 | 3 | Athu-Bthu | 55 |
| 1 | 1 | 5 | 5 | 3 | Bthu-low | 103 |
| 1 | 1 | 6 | 6 | 3 | Bthu-low | 101 |
| 1 | 2 | 7 | 1 | 2 | Mixed-low | 68 |
| 1 | 2 | 8 | 2 | 2 | Mixed-low | 67 |
| 1 | 2 | 9 | 3 | 2 | Athu-Bthu | 55 |
| 1 | 2 | 10 | 4 | 2 | Athu-Bthu | 57 |
| 1 | 2 | 11 | 5 | 2 | Athu-Bthu | 56 |
| 1 | 2 | 12 | 6 | 2 | Mixed-low | 64 |
| 1 | 3 | 13 | 1 | 3 | Mixed-high | 30 |
| 1 | 3 | 14 | 2 | 3 | Mixed-high | 21 |
| 1 | 3 | 15 | 3 | 3 | Chrys-acet | 94 |
| 1 | 3 | 16 | 4 | 3 | Chrys-acet | 100 |
| 1 | 3 | 17 | 5 | 3 | Bthu-low | 101 |
| 1 | 3 | 18 | 6 | 3 | Bthu-low | 104 |
| 1 | 4 | 19 | 1 | 2 | Athu-Bthu | 56 |
| 1 | 4 | 20 | 2 | 2 | Athu-Bthu | 57 |
| 1 | 4 | 21 | 3 | 2 | Bthu-low | 101 |
| 1 | 4 | 22 | 4 | 2 | Bthu-low | 104 |
| 1 | 4 | 23 | 5 | 2 | Athu-Bthu | 55 |
| 1 | 4 | 24 | 6 | 2 | Bthu-low | 103 |
| 1 | 5 | 25 | 1 | 3 | Chrys-acet | 95 |
| 1 | 5 | 26 | 2 | 3 | Bthu-high | 14 |
| 1 | 5 | 27 | 3 | 3 | Bthu-high | 11 |
| 1 | 5 | 28 | 4 | 3 | Mixed-high | 21 |
| 1 | 5 | 29 | 5 | 3 | Chrys-acet | 94 |
| 1 | 5 | 30 | 6 | 3 | Mixed-high | 30 |
| 1 | 6 | 31 | 1 | 6 | Mixed-low | 68 |
| 1 | 6 | 32 | 2 | 6 | Bthu-low | 104 |
| 1 | 6 | 33 | 3 | 6 | Bthu-high | 11 |
| 1 | 6 | 34 | 4 | 6 | Mixed-high | 30 |
| 1 | 6 | 35 | 5 | 6 | Chrys-acet | 100 |
| 1 | 6 | 36 | 6 | 6 | Athu-Bthu | 55 |

**Table S1-2 *(continued)***: *This section shows the results for the second part of Table S1-2.*

| Block | Plot | Plant | Plot position | Plot-level chemotype richness | Chemotype | Daughter |
|---|---|---|---|---|---|---|
| 1 | 7 | 37 | 1 | 6 | Athu-Bthu | 55 |
| 1 | 7 | 38 | 2 | 6 | Bthu-low | 101 |
| 1 | 7 | 39 | 3 | 6 | Bthu-high | 17 |
| 1 | 7 | 40 | 4 | 6 | Mixed-low | 64 |
| 1 | 7 | 41 | 5 | 6 | Mixed-high | 21 |
| 1 | 7 | 42 | 6 | 6 | Chrys-acet | 94 |
| 1 | 8 | 43 | 1 | 2 | Chrys-acet | 94 |
| 1 | 8 | 44 | 2 | 2 | Bthu-low | 101 |
| 1 | 8 | 45 | 3 | 2 | Chrys-acet | 100 |
| 1 | 8 | 46 | 4 | 2 | Chrys-acet | 95 |
| 1 | 8 | 47 | 5 | 2 | Bthu-low | 103 |
| 1 | 8 | 48 | 6 | 2 | Bthu-low | 104 |
| 1 | 9 | 49 | 1 | 1 | Mixed-high | 21 |
| 1 | 9 | 50 | 2 | 1 | Mixed-high | 30 |
| 1 | 9 | 51 | 3 | 1 | Mixed-high | 21 |
| 1 | 9 | 52 | 4 | 1 | Mixed-high | 30 |
| 1 | 9 | 53 | 5 | 1 | Mixed-high | 23 |
| 1 | 9 | 54 | 6 | 1 | Mixed-high | 23 |
| 1 | 10 | 55 | 1 | 1 | Bthu-high | 11 |
| 1 | 10 | 56 | 2 | 1 | Bthu-high | 17 |
| 1 | 10 | 57 | 3 | 1 | Bthu-high | 14 |
| 1 | 10 | 58 | 4 | 1 | Bthu-high | 14 |
| 1 | 10 | 59 | 5 | 1 | Bthu-high | 11 |
| 1 | 10 | 60 | 6 | 1 | Bthu-high | 17 |
| 1 | 11 | 61 | 1 | 2 | Bthu-high | 17 |
| 1 | 11 | 62 | 2 | 2 | Bthu-high | 11 |
| 1 | 11 | 63 | 3 | 2 | Chrys-acet | 94 |
| 1 | 11 | 64 | 4 | 2 | Chrys-acet | 95 |
| 1 | 11 | 65 | 5 | 2 | Chrys-acet | 100 |
| 1 | 11 | 66 | 6 | 2 | Bthu-high | 14 |
| 1 | 12 | 67 | 1 | 2 | Bthu-low | 103 |
| 1 | 12 | 68 | 2 | 2 | Bthu-low | 101 |
| 1 | 12 | 69 | 3 | 2 | Mixed-high | 23 |
| 1 | 12 | 70 | 4 | 2 | Bthu-low | 104 |
| 1 | 12 | 71 | 5 | 2 | Mixed-high | 21 |
| 1 | 12 | 72 | 6 | 2 | Mixed-high | 30 |
| 1 | 13 | 73 | 1 | 3 | Bthu-low | 104 |
| 1 | 13 | 74 | 2 | 3 | Mixed-low | 68 |
| 1 | 13 | 75 | 3 | 3 | Mixed-low | 64 |
| 1 | 13 | 76 | 4 | 3 | Bthu-low | 101 |
| 1 | 13 | 77 | 5 | 3 | Athu-Bthu | 56 |
| 1 | 13 | 78 | 6 | 3 | Athu-Bthu | 57 |
| 1 | 14 | 79 | 1 | 3 | Athu-Bthu | 55 |
| 1 | 14 | 80 | 2 | 3 | Mixed-low | 64 |
| 1 | 14 | 81 | 3 | 3 | Bthu-high | 11 |
| 1 | 14 | 82 | 4 | 3 | Athu-Bthu | 56 |
| 1 | 14 | 83 | 5 | 3 | Mixed-low | 68 |
| 1 | 14 | 84 | 6 | 3 | Bthu-high | 17 |

**Table S1-2 *(continued)***: *This section shows the results for the third part of Table S1-2.*

| Block | Plot | Plant | Plot position | Plot-level chemotype richness | Chemotype | Daughter |
|---|---|---|---|---|---|---|
| 2 | 15 | 85 | 1 | 2 | Bthu-low | 101 |
| 2 | 15 | 86 | 2 | 2 | Mixed-low | 64 |
| 2 | 15 | 87 | 3 | 2 | Mixed-low | 67 |
| 2 | 15 | 88 | 4 | 2 | Mixed-low | 68 |
| 2 | 15 | 89 | 5 | 2 | Bthu-low | 104 |
| 2 | 15 | 90 | 6 | 2 | Bthu-low | 103 |
| 2 | 16 | 91 | 1 | 6 | Athu-Bthu | 57 |
| 2 | 16 | 92 | 2 | 6 | Chrys-acet | 95 |
| 2 | 16 | 93 | 3 | 6 | Bthu-low | 103 |
| 2 | 16 | 94 | 4 | 6 | Bthu-high | 14 |
| 2 | 16 | 95 | 5 | 6 | Mixed-low | 67 |
| 2 | 16 | 96 | 6 | 6 | Mixed-high | 23 |
| 2 | 17 | 97 | 1 | 2 | Mixed-high | 21 |
| 2 | 17 | 98 | 2 | 2 | Bthu-high | 14 |
| 2 | 17 | 99 | 3 | 2 | Mixed-high | 23 |
| 2 | 17 | 100 | 4 | 2 | Mixed-high | 30 |
| 2 | 17 | 101 | 5 | 2 | Bthu-high | 17 |
| 2 | 17 | 102 | 6 | 2 | Bthu-high | 11 |
| 2 | 18 | 103 | 1 | 3 | Bthu-high | 17 |
| 2 | 18 | 104 | 2 | 3 | Mixed-high | 23 |
| 2 | 18 | 105 | 3 | 3 | Athu-Bthu | 57 |
| 2 | 18 | 106 | 4 | 3 | Bthu-high | 14 |
| 2 | 18 | 107 | 5 | 3 | Mixed-high | 30 |
| 2 | 18 | 108 | 6 | 3 | Athu-Bthu | 55 |
| 2 | 19 | 109 | 1 | 2 | Mixed-low | 68 |
| 2 | 19 | 110 | 2 | 2 | Mixed-low | 64 |
| 2 | 19 | 111 | 3 | 2 | Chrys-acet | 95 |
| 2 | 19 | 112 | 4 | 2 | Mixed-low | 67 |
| 2 | 19 | 113 | 5 | 2 | Chrys-acet | 100 |
| 2 | 19 | 114 | 6 | 2 | Chrys-acet | 94 |
| 2 | 20 | 115 | 1 | 2 | Bthu-high | 17 |
| 2 | 20 | 116 | 2 | 2 | Athu-Bthu | 56 |
| 2 | 20 | 117 | 3 | 2 | Bthu-high | 14 |
| 2 | 20 | 118 | 4 | 2 | Athu-Bthu | 57 |
| 2 | 20 | 119 | 5 | 2 | Bthu-high | 11 |
| 2 | 20 | 120 | 6 | 2 | Athu-Bthu | 55 |
| 2 | 21 | 121 | 1 | 3 | Mixed-high | 23 |
| 2 | 21 | 122 | 2 | 3 | Mixed-high | 30 |
| 2 | 21 | 123 | 3 | 3 | Athu-Bthu | 55 |
| 2 | 21 | 124 | 4 | 3 | Chrys-acet | 95 |
| 2 | 21 | 125 | 5 | 3 | Athu-Bthu | 57 |
| 2 | 21 | 126 | 6 | 3 | Chrys-acet | 100 |
| 2 | 22 | 127 | 1 | 6 | Athu-Bthu | 56 |
| 2 | 22 | 128 | 2 | 6 | Mixed-high | 23 |
| 2 | 22 | 129 | 3 | 6 | Bthu-high | 14 |
| 2 | 22 | 130 | 4 | 6 | Bthu-low | 104 |
| 2 | 22 | 131 | 5 | 6 | Mixed-low | 64 |
| 2 | 22 | 132 | 6 | 6 | Chrys-acet | 95 |

**Table S1-2 *(continued)*:** *This section shows the results for the fourth part of Table S1-2.*

| Block | Plot | Plant | Plot position | Plot-level chemotype richness | Chemotype | Daughter |
|---|---|---|---|---|---|---|
| 2 | 23 | 133 | 1 | 3 | Bthu-high | 14 |
| 2 | 23 | 134 | 2 | 3 | Mixed-low | 67 |
| 2 | 23 | 135 | 3 | 3 | Bthu-high | 11 |
| 2 | 23 | 136 | 4 | 3 | Mixed-low | 68 |
| 2 | 23 | 137 | 5 | 3 | Bthu-low | 104 |
| 2 | 23 | 138 | 6 | 3 | Bthu-low | 103 |
| 2 | 24 | 139 | 1 | 1 | Bthu-low | 104 |
| 2 | 24 | 140 | 2 | 1 | Bthu-low | 104 |
| 2 | 24 | 141 | 3 | 1 | Bthu-low | 101 |
| 2 | 24 | 142 | 4 | 1 | Bthu-low | 101 |
| 2 | 24 | 143 | 5 | 1 | Bthu-low | 103 |
| 2 | 24 | 144 | 6 | 1 | Bthu-low | 103 |
| 2 | 25 | 145 | 1 | 1 | Mixed-low | 67 |
| 2 | 25 | 146 | 2 | 1 | Mixed-low | 64 |
| 2 | 25 | 147 | 3 | 1 | Mixed-low | 68 |
| 2 | 25 | 148 | 4 | 1 | Mixed-low | 67 |
| 2 | 25 | 149 | 5 | 1 | Mixed-low | 68 |
| 2 | 25 | 150 | 6 | 1 | Mixed-low | 64 |
| 2 | 26 | 151 | 1 | 3 | Bthu-high | 14 |
| 2 | 26 | 152 | 2 | 3 | Bthu-high | 17 |
| 2 | 26 | 153 | 3 | 3 | Mixed-low | 67 |
| 2 | 26 | 154 | 4 | 3 | Chrys-acet | 95 |
| 2 | 26 | 155 | 5 | 3 | Mixed-low | 64 |
| 2 | 26 | 156 | 6 | 3 | Chrys-acet | 100 |
| 2 | 27 | 157 | 1 | 3 | Bthu-high | 14 |
| 2 | 27 | 158 | 2 | 3 | Bthu-high | 17 |
| 2 | 27 | 159 | 3 | 3 | Bthu-low | 104 |
| 2 | 27 | 160 | 4 | 3 | Mixed-high | 23 |
| 2 | 27 | 161 | 5 | 3 | Mixed-high | 30 |
| 2 | 27 | 162 | 6 | 3 | Bthu-low | 103 |
| 2 | 28 | 163 | 1 | 2 | Mixed-high | 23 |
| 2 | 28 | 164 | 2 | 2 | Chrys-acet | 95 |
| 2 | 28 | 165 | 3 | 2 | Chrys-acet | 94 |
| 2 | 28 | 166 | 4 | 2 | Chrys-acet | 100 |
| 2 | 28 | 167 | 5 | 2 | Mixed-high | 21 |
| 2 | 28 | 168 | 6 | 2 | Mixed-high | 30 |
| 3 | 29 | 169 | 1 | 2 | Bthu-low | 104 |
| 3 | 29 | 170 | 2 | 2 | Bthu-high | 14 |
| 3 | 29 | 171 | 3 | 2 | Bthu-low | 103 |
| 3 | 29 | 172 | 4 | 2 | Bthu-high | 17 |
| 3 | 29 | 173 | 5 | 2 | Bthu-low | 101 |
| 3 | 29 | 174 | 6 | 2 | Bthu-high | 11 |
| 3 | 30 | 175 | 1 | 3 | Mixed-high | 23 |
| 3 | 30 | 176 | 2 | 3 | Mixed-high | 21 |
| 3 | 30 | 177 | 3 | 3 | Bthu-high | 11 |
| 3 | 30 | 178 | 4 | 3 | Bthu-low | 101 |
| 3 | 30 | 179 | 5 | 3 | Bthu-high | 17 |
| 3 | 30 | 180 | 6 | 3 | Bthu-low | 103 |

**Table S1-2 *(continued)***: *This section shows the results for the fifth part of Table S1-2.*

| Block | Plot | Plant | Plot position | Plot-level chemotype richness | Chemotype | Daughter |
|---|---|---|---|---|---|---|
| 3 | 31 | 181 | 1 | 2 | Mixed-low | 68 |
| 3 | 31 | 182 | 2 | 2 | Mixed-high | 21 |
| 3 | 31 | 183 | 3 | 2 | Mixed-high | 30 |
| 3 | 31 | 184 | 4 | 2 | Mixed-low | 67 |
| 3 | 31 | 185 | 5 | 2 | Mixed-high | 23 |
| 3 | 31 | 186 | 6 | 2 | Mixed-low | 64 |
| 3 | 32 | 187 | 1 | 6 | Bthu-low | 103 |
| 3 | 32 | 188 | 2 | 6 | Mixed-low | 68 |
| 3 | 32 | 189 | 3 | 6 | Chrys-acet | 94 |
| 3 | 32 | 190 | 4 | 6 | Mixed-high | 23 |
| 3 | 32 | 191 | 5 | 6 | Athu-Bthu | 56 |
| 3 | 32 | 192 | 6 | 6 | Bthu-high | 11 |
| 3 | 33 | 193 | 1 | 2 | Athu-Bthu | 56 |
| 3 | 33 | 194 | 2 | 2 | Athu-Bthu | 55 |
| 3 | 33 | 195 | 3 | 2 | Chrys-acet | 100 |
| 3 | 33 | 196 | 4 | 2 | Chrys-acet | 95 |
| 3 | 33 | 197 | 5 | 2 | Chrys-acet | 94 |
| 3 | 33 | 198 | 6 | 2 | Athu-Bthu | 57 |
| 3 | 34 | 199 | 1 | 1 | Athu-Bthu | 56 |
| 3 | 34 | 200 | 2 | 1 | Athu-Bthu | 55 |
| 3 | 34 | 201 | 3 | 1 | Athu-Bthu | 57 |
| 3 | 34 | 202 | 4 | 1 | Athu-Bthu | 56 |
| 3 | 34 | 203 | 5 | 1 | Athu-Bthu | 57 |
| 3 | 34 | 204 | 6 | 1 | Athu-Bthu | 55 |
| 3 | 35 | 205 | 1 | 1 | Chrys-acet | 94 |
| 3 | 35 | 206 | 2 | 1 | Chrys-acet | 100 |
| 3 | 35 | 207 | 3 | 1 | Chrys-acet | 100 |
| 3 | 35 | 208 | 4 | 1 | Chrys-acet | 95 |
| 3 | 35 | 209 | 5 | 1 | Chrys-acet | 95 |
| 3 | 35 | 210 | 6 | 1 | Chrys-acet | 94 |
| 3 | 36 | 211 | 1 | 3 | Chrys-acet | 100 |
| 3 | 36 | 212 | 2 | 3 | Mixed-high | 21 |
| 3 | 36 | 213 | 3 | 3 | Bthu-high | 11 |
| 3 | 36 | 214 | 4 | 3 | Bthu-high | 14 |
| 3 | 36 | 215 | 5 | 3 | Chrys-acet | 94 |
| 3 | 36 | 216 | 6 | 3 | Mixed-high | 23 |
| 3 | 37 | 217 | 1 | 3 | Athu-Bthu | 56 |
| 3 | 37 | 218 | 2 | 3 | Mixed-low | 67 |
| 3 | 37 | 219 | 3 | 3 | Chrys-acet | 94 |
| 3 | 37 | 220 | 4 | 3 | Athu-Bthu | 57 |
| 3 | 37 | 221 | 5 | 3 | Chrys-acet | 100 |
| 3 | 37 | 222 | 6 | 3 | Mixed-low | 68 |
| 3 | 38 | 223 | 1 | 2 | Mixed-high | 30 |
| 3 | 38 | 224 | 2 | 2 | Mixed-high | 21 |
| 3 | 38 | 225 | 3 | 2 | Athu-Bthu | 56 |
| 3 | 38 | 226 | 4 | 2 | Mixed-high | 23 |
| 3 | 38 | 227 | 5 | 2 | Athu-Bthu | 57 |
| 3 | 38 | 228 | 6 | 2 | Athu-Bthu | 55 |

**Table S1-2 *(continued)***: *This section shows the results for the sixth part of Table S1-2.*

| Block | Plot | Plant | Plot position | Plot-level chemotype richness | Chemotype | Daughter |
|---|---|---|---|---|---|---|
| 3 | 39 | 229 | 1 | 3 | Mixed-low | 67 |
| 3 | 39 | 230 | 2 | 3 | Mixed-low | 64 |
| 3 | 39 | 231 | 3 | 3 | Mixed-high | 21 |
| 3 | 39 | 232 | 4 | 3 | Bthu-high | 17 |
| 3 | 39 | 233 | 5 | 3 | Bthu-high | 11 |
| 3 | 39 | 234 | 6 | 3 | Mixed-high | 30 |
| 3 | 40 | 235 | 1 | 3 | Mixed-low | 68 |
| 3 | 40 | 236 | 2 | 3 | Mixed-high | 21 |
| 3 | 40 | 237 | 3 | 3 | Athu-Bthu | 57 |
| 3 | 40 | 238 | 4 | 3 | Athu-Bthu | 56 |
| 3 | 40 | 239 | 5 | 3 | Mixed-high | 23 |
| 3 | 40 | 240 | 6 | 3 | Mixed-low | 64 |
| 3 | 41 | 241 | 1 | 2 | Bthu-high | 11 |
| 3 | 41 | 242 | 2 | 2 | Bthu-high | 14 |
| 3 | 41 | 243 | 3 | 2 | Mixed-low | 67 |
| 3 | 41 | 244 | 4 | 2 | Mixed-low | 64 |
| 3 | 41 | 245 | 5 | 2 | Mixed-low | 68 |
| 3 | 41 | 246 | 6 | 2 | Bthu-high | 17 |
| 3 | 42 | 247 | 1 | 6 | Chrys-acet | 100 |
| 3 | 42 | 248 | 2 | 6 | Bthu-low | 103 |
| 3 | 42 | 249 | 3 | 6 | Mixed-low | 64 |
| 3 | 42 | 250 | 4 | 6 | Mixed-high | 21 |
| 3 | 42 | 251 | 5 | 6 | Bthu-high | 11 |
| 3 | 42 | 252 | 6 | 6 | Athu-Bthu | 55 |
| 4 | 43 | 253 | 1 | 1 | Chrys-acet | 100 |
| 4 | 43 | 254 | 2 | 1 | Chrys-acet | 95 |
| 4 | 43 | 255 | 3 | 1 | Chrys-acet | 95 |
| 4 | 43 | 256 | 4 | 1 | Chrys-acet | 94 |
| 4 | 43 | 257 | 5 | 1 | Chrys-acet | 94 |
| 4 | 43 | 258 | 6 | 1 | Chrys-acet | 100 |
| 4 | 44 | 259 | 1 | 1 | Athu-Bthu | 57 |
| 4 | 44 | 260 | 2 | 1 | Athu-Bthu | 55 |
| 4 | 44 | 261 | 3 | 1 | Athu-Bthu | 56 |
| 4 | 44 | 262 | 4 | 1 | Athu-Bthu | 55 |
| 4 | 44 | 263 | 5 | 1 | Athu-Bthu | 56 |
| 4 | 44 | 264 | 6 | 1 | Athu-Bthu | 57 |
| 4 | 45 | 265 | 1 | 2 | Athu-Bthu | 55 |
| 4 | 45 | 266 | 2 | 2 | Athu-Bthu | 57 |
| 4 | 45 | 267 | 3 | 2 | Athu-Bthu | 56 |
| 4 | 45 | 268 | 4 | 2 | Chrys-acet | 94 |
| 4 | 45 | 269 | 5 | 2 | Chrys-acet | 100 |
| 4 | 45 | 270 | 6 | 2 | Chrys-acet | 95 |
| 4 | 46 | 271 | 1 | 2 | Bthu-high | 17 |
| 4 | 46 | 272 | 2 | 2 | Bthu-high | 14 |
| 4 | 46 | 273 | 3 | 2 | Mixed-low | 64 |
| 4 | 46 | 274 | 4 | 2 | Mixed-low | 68 |
| 4 | 46 | 275 | 5 | 2 | Bthu-high | 11 |
| 4 | 46 | 276 | 6 | 2 | Mixed-low | 67 |

**Table S1-2 *(continued)***: *This section shows the results for the seventh part of Table S1-2.*

| Block | Plot | Plant | Plot position | Plot-level chemotype richness | Chemotype | Daughter |
|---|---|---|---|---|---|---|
| 4 | 47 | 277 | 1 | 6 | Athu-Bthu | 57 |
| 4 | 47 | 278 | 2 | 6 | Mixed-low | 67 |
| 4 | 47 | 279 | 3 | 6 | Mixed-high | 21 |
| 4 | 47 | 280 | 4 | 6 | Bthu-low | 104 |
| 4 | 47 | 281 | 5 | 6 | Chrys-acet | 100 |
| 4 | 47 | 282 | 6 | 6 | Bthu-high | 14 |
| 4 | 48 | 283 | 1 | 3 | Bthu-low | 101 |
| 4 | 48 | 284 | 2 | 3 | Bthu-high | 11 |
| 4 | 48 | 285 | 3 | 3 | Mixed-high | 21 |
| 4 | 48 | 286 | 4 | 3 | Bthu-low | 103 |
| 4 | 48 | 287 | 5 | 3 | Bthu-high | 17 |
| 4 | 48 | 288 | 6 | 3 | Mixed-high | 23 |
| 4 | 49 | 289 | 1 | 3 | Mixed-low | 67 |
| 4 | 49 | 290 | 2 | 3 | Mixed-high | 21 |
| 4 | 49 | 291 | 3 | 3 | Mixed-high | 30 |
| 4 | 49 | 292 | 4 | 3 | Bthu-high | 11 |
| 4 | 49 | 293 | 5 | 3 | Mixed-low | 68 |
| 4 | 49 | 294 | 6 | 3 | Bthu-high | 17 |
| 4 | 50 | 295 | 1 | 6 | Mixed-high | 23 |
| 4 | 50 | 296 | 2 | 6 | Bthu-high | 14 |
| 4 | 50 | 297 | 3 | 6 | Bthu-low | 101 |
| 4 | 50 | 298 | 4 | 6 | Mixed-low | 67 |
| 4 | 50 | 299 | 5 | 6 | Athu-Bthu | 56 |
| 4 | 50 | 300 | 6 | 6 | Chrys-acet | 94 |
| 4 | 51 | 301 | 1 | 3 | Bthu-high | 14 |
| 4 | 51 | 302 | 2 | 3 | Bthu-high | 11 |
| 4 | 51 | 303 | 3 | 3 | Chrys-acet | 95 |
| 4 | 51 | 304 | 4 | 3 | Chrys-acet | 100 |
| 4 | 51 | 305 | 5 | 3 | Mixed-high | 21 |
| 4 | 51 | 306 | 6 | 3 | Mixed-high | 23 |
| 4 | 52 | 307 | 1 | 3 | Athu-Bthu | 56 |
| 4 | 52 | 308 | 2 | 3 | Chrys-acet | 100 |
| 4 | 52 | 309 | 3 | 3 | Athu-Bthu | 55 |
| 4 | 52 | 310 | 4 | 3 | Mixed-low | 64 |
| 4 | 52 | 311 | 5 | 3 | Mixed-low | 67 |
| 4 | 52 | 312 | 6 | 3 | Chrys-acet | 95 |
| 4 | 53 | 313 | 1 | 2 | Mixed-low | 68 |
| 4 | 53 | 314 | 2 | 2 | Mixed-high | 23 |
| 4 | 53 | 315 | 3 | 2 | Mixed-low | 64 |
| 4 | 53 | 316 | 4 | 2 | Mixed-high | 21 |
| 4 | 53 | 317 | 5 | 2 | Mixed-low | 67 |
| 4 | 53 | 318 | 6 | 2 | Mixed-high | 30 |
| 4 | 54 | 319 | 1 | 2 | Bthu-high | 11 |
| 4 | 54 | 320 | 2 | 2 | Bthu-high | 14 |
| 4 | 54 | 321 | 3 | 2 | Bthu-low | 104 |
| 4 | 54 | 322 | 4 | 2 | Bthu-high | 17 |
| 4 | 54 | 323 | 5 | 2 | Bthu-low | 101 |
| 4 | 54 | 324 | 6 | 2 | Bthu-low | 103 |

**Table S1-2 *(continued)***: *This section shows the results for the eighth part of Table S1-2.*

| Block | Plot | Plant | Plot position | Plot-level chemotype richness | Chemotype | Daughter |
|---|---|---|---|---|---|---|
| 4 | 55 | 325 | 1 | 2 | Athu-Bthu | 56 |
| 4 | 55 | 326 | 2 | 2 | Athu-Bthu | 57 |
| 4 | 55 | 327 | 3 | 2 | Mixed-high | 30 |
| 4 | 55 | 328 | 4 | 2 | Athu-Bthu | 55 |
| 4 | 55 | 329 | 5 | 2 | Mixed-high | 21 |
| 4 | 55 | 330 | 6 | 2 | Mixed-high | 23 |
| 4 | 56 | 331 | 1 | 3 | Mixed-low | 68 |
| 4 | 56 | 332 | 2 | 3 | Athu-Bthu | 56 |
| 4 | 56 | 333 | 3 | 3 | Mixed-low | 67 |
| 4 | 56 | 334 | 4 | 3 | Athu-Bthu | 55 |
| 4 | 56 | 335 | 5 | 3 | Mixed-high | 23 |
| 4 | 56 | 336 | 6 | 3 | Mixed-high | 21 |
| 5 | 57 | 337 | 1 | 1 | Mixed-low | 67 |
| 5 | 57 | 338 | 2 | 1 | Mixed-low | 64 |
| 5 | 57 | 339 | 3 | 1 | Mixed-low | 68 |
| 5 | 57 | 340 | 4 | 1 | Mixed-low | 64 |
| 5 | 57 | 341 | 5 | 1 | Mixed-low | 68 |
| 5 | 57 | 342 | 6 | 1 | Mixed-low | 67 |
| 5 | 58 | 343 | 1 | 3 | Bthu-high | 17 |
| 5 | 58 | 344 | 2 | 3 | Mixed-low | 64 |
| 5 | 58 | 345 | 3 | 3 | Bthu-high | 14 |
| 5 | 58 | 346 | 4 | 3 | Chrys-acet | 94 |
| 5 | 58 | 347 | 5 | 3 | Mixed-low | 68 |
| 5 | 58 | 348 | 6 | 3 | Chrys-acet | 95 |
| 5 | 59 | 349 | 1 | 6 | Athu-Bthu | 57 |
| 5 | 59 | 350 | 2 | 6 | Mixed-low | 67 |
| 5 | 59 | 351 | 3 | 6 | Bthu-high | 17 |
| 5 | 59 | 352 | 4 | 6 | Bthu-low | 103 |
| 5 | 59 | 353 | 5 | 6 | Chrys-acet | 95 |
| 5 | 59 | 354 | 6 | 6 | Mixed-high | 21 |
| 5 | 60 | 355 | 1 | 3 | Bthu-high | 14 |
| 5 | 60 | 356 | 2 | 3 | Bthu-low | 103 |
| 5 | 60 | 357 | 3 | 3 | Bthu-low | 104 |
| 5 | 60 | 358 | 4 | 3 | Mixed-high | 23 |
| 5 | 60 | 359 | 5 | 3 | Bthu-high | 17 |
| 5 | 60 | 360 | 6 | 3 | Mixed-high | 30 |
| 5 | 61 | 361 | 1 | 3 | Mixed-high | 23 |
| 5 | 61 | 362 | 2 | 3 | Mixed-high | 30 |
| 5 | 61 | 363 | 3 | 3 | Athu-Bthu | 57 |
| 5 | 61 | 364 | 4 | 3 | Athu-Bthu | 56 |
| 5 | 61 | 365 | 5 | 3 | Bthu-high | 17 |
| 5 | 61 | 366 | 6 | 3 | Bthu-high | 14 |
| 5 | 62 | 367 | 1 | 2 | Mixed-high | 21 |
| 5 | 62 | 368 | 2 | 2 | Chrys-acet | 95 |
| 5 | 62 | 369 | 3 | 2 | Chrys-acet | 100 |
| 5 | 62 | 370 | 4 | 2 | Mixed-high | 30 |
| 5 | 62 | 371 | 5 | 2 | Chrys-acet | 94 |
| 5 | 62 | 372 | 6 | 2 | Mixed-high | 23 |

**Table S1-2 *(continued)***: *This section shows the results for the ninth part of Table S1-2.*

| Block | Plot | Plant | Plot position | Plot-level chemotype richness | Chemotype | Daughter |
|---|---|---|---|---|---|---|
| 5 | 63 | 373 | 1 | 1 | Bthu-low | 101 |
| 5 | 63 | 374 | 2 | 1 | Bthu-low | 104 |
| 5 | 63 | 375 | 3 | 1 | Bthu-low | 103 |
| 5 | 63 | 376 | 4 | 1 | Bthu-low | 104 |
| 5 | 63 | 377 | 5 | 1 | Bthu-low | 101 |
| 5 | 63 | 378 | 6 | 1 | Bthu-low | 103 |
| 5 | 64 | 379 | 1 | 3 | Mixed-high | 30 |
| 5 | 64 | 380 | 2 | 3 | Athu-Bthu | 57 |
| 5 | 64 | 381 | 3 | 3 | Mixed-high | 23 |
| 5 | 64 | 382 | 4 | 3 | Athu-Bthu | 56 |
| 5 | 64 | 383 | 5 | 3 | Chrys-acet | 94 |
| 5 | 64 | 384 | 6 | 3 | Chrys-acet | 95 |
| 5 | 65 | 385 | 1 | 6 | Bthu-high | 17 |
| 5 | 65 | 386 | 2 | 6 | Mixed-high | 30 |
| 5 | 65 | 387 | 3 | 6 | Mixed-low | 68 |
| 5 | 65 | 388 | 4 | 6 | Athu-Bthu | 56 |
| 5 | 65 | 389 | 5 | 6 | Chrys-acet | 95 |
| 5 | 65 | 390 | 6 | 6 | Bthu-low | 103 |
| 5 | 66 | 391 | 1 | 3 | Bthu-high | 11 |
| 5 | 66 | 392 | 2 | 3 | Bthu-low | 103 |
| 5 | 66 | 393 | 3 | 3 | Mixed-low | 64 |
| 5 | 66 | 394 | 4 | 3 | Bthu-low | 104 |
| 5 | 66 | 395 | 5 | 3 | Bthu-high | 14 |
| 5 | 66 | 396 | 6 | 3 | Mixed-low | 68 |
| 5 | 67 | 397 | 1 | 2 | Mixed-high | 30 |
| 5 | 67 | 398 | 2 | 2 | Mixed-high | 23 |
| 5 | 67 | 399 | 3 | 2 | Bthu-high | 11 |
| 5 | 67 | 400 | 4 | 2 | Bthu-high | 14 |
| 5 | 67 | 401 | 5 | 2 | Bthu-high | 17 |
| 5 | 67 | 402 | 6 | 2 | Mixed-high | 21 |
| 5 | 68 | 403 | 1 | 2 | Mixed-low | 67 |
| 5 | 68 | 404 | 2 | 2 | Mixed-low | 68 |
| 5 | 68 | 405 | 3 | 2 | Chrys-acet | 94 |
| 5 | 68 | 406 | 4 | 2 | Chrys-acet | 100 |
| 5 | 68 | 407 | 5 | 2 | Chrys-acet | 95 |
| 5 | 68 | 408 | 6 | 2 | Mixed-low | 64 |
| 5 | 69 | 409 | 1 | 2 | Athu-Bthu | 56 |
| 5 | 69 | 410 | 2 | 2 | Bthu-high | 11 |
| 5 | 69 | 411 | 3 | 2 | Bthu-high | 17 |
| 5 | 69 | 412 | 4 | 2 | Bthu-high | 14 |
| 5 | 69 | 413 | 5 | 2 | Athu-Bthu | 55 |
| 5 | 69 | 414 | 6 | 2 | Athu-Bthu | 57 |
| 5 | 70 | 415 | 1 | 2 | Mixed-low | 68 |
| 5 | 70 | 416 | 2 | 2 | Mixed-low | 64 |
| 5 | 70 | 417 | 3 | 2 | Mixed-low | 67 |
| 5 | 70 | 418 | 4 | 2 | Bthu-low | 104 |
| 5 | 70 | 419 | 5 | 2 | Bthu-low | 103 |
| 5 | 70 | 420 | 6 | 2 | Bthu-low | 101 |

**Table S1-2 *(continued)*:** *This section shows the results for the tenth part of Table S1-2.*

| Block | Plot | Plant | Plot position | Plot-level chemotype richness | Chemotype | Daughter |
|---|---|---|---|---|---|---|
| 6 | 71 | 421 | 1 | 2 | Bthu-low | 101 |
| 6 | 71 | 422 | 2 | 2 | Athu-Bthu | 57 |
| 6 | 71 | 423 | 3 | 2 | Athu-Bthu | 56 |
| 6 | 71 | 424 | 4 | 2 | Bthu-low | 103 |
| 6 | 71 | 425 | 5 | 2 | Athu-Bthu | 55 |
| 6 | 71 | 426 | 6 | 2 | Bthu-low | 104 |
| 6 | 72 | 427 | 1 | 1 | Bthu-high | 14 |
| 6 | 72 | 428 | 2 | 1 | Bthu-high | 17 |
| 6 | 72 | 429 | 3 | 1 | Bthu-high | 11 |
| 6 | 72 | 430 | 4 | 1 | Bthu-high | 11 |
| 6 | 72 | 431 | 5 | 1 | Bthu-high | 17 |
| 6 | 72 | 432 | 6 | 1 | Bthu-high | 14 |
| 6 | 73 | 433 | 1 | 2 | Chrys-acet | 94 |
| 6 | 73 | 434 | 2 | 2 | Bthu-low | 103 |
| 6 | 73 | 435 | 3 | 2 | Chrys-acet | 95 |
| 6 | 73 | 436 | 4 | 2 | Bthu-low | 104 |
| 6 | 73 | 437 | 5 | 2 | Chrys-acet | 100 |
| 6 | 73 | 438 | 6 | 2 | Bthu-low | 101 |
| 6 | 74 | 439 | 1 | 6 | Athu-Bthu | 55 |
| 6 | 74 | 440 | 2 | 6 | Mixed-low | 68 |
| 6 | 74 | 441 | 3 | 6 | Bthu-low | 101 |
| 6 | 74 | 442 | 4 | 6 | Bthu-high | 11 |
| 6 | 74 | 443 | 5 | 6 | Mixed-high | 30 |
| 6 | 74 | 444 | 6 | 6 | Chrys-acet | 94 |
| 6 | 75 | 445 | 1 | 2 | Mixed-high | 30 |
| 6 | 75 | 446 | 2 | 2 | Bthu-low | 104 |
| 6 | 75 | 447 | 3 | 2 | Mixed-high | 21 |
| 6 | 75 | 448 | 4 | 2 | Bthu-low | 103 |
| 6 | 75 | 449 | 5 | 2 | Mixed-high | 23 |
| 6 | 75 | 450 | 6 | 2 | Bthu-low | 101 |
| 6 | 76 | 451 | 1 | 3 | Bthu-low | 101 |
| 6 | 76 | 452 | 2 | 3 | Chrys-acet | 100 |
| 6 | 76 | 453 | 3 | 3 | Mixed-high | 30 |
| 6 | 76 | 454 | 4 | 3 | Bthu-low | 104 |
| 6 | 76 | 455 | 5 | 3 | Mixed-high | 21 |
| 6 | 76 | 456 | 6 | 3 | Chrys-acet | 95 |
| 6 | 77 | 457 | 1 | 3 | Athu-Bthu | 55 |
| 6 | 77 | 458 | 2 | 3 | Bthu-low | 104 |
| 6 | 77 | 459 | 3 | 3 | Mixed-low | 64 |
| 6 | 77 | 460 | 4 | 3 | Athu-Bthu | 57 |
| 6 | 77 | 461 | 5 | 3 | Bthu-low | 101 |
| 6 | 77 | 462 | 6 | 3 | Mixed-low | 67 |
| 6 | 78 | 463 | 1 | 2 | Athu-Bthu | 56 |
| 6 | 78 | 464 | 2 | 2 | Athu-Bthu | 55 |
| 6 | 78 | 465 | 3 | 2 | Mixed-low | 67 |
| 6 | 78 | 466 | 4 | 2 | Mixed-low | 64 |
| 6 | 78 | 467 | 5 | 2 | Athu-Bthu | 57 |
| 6 | 78 | 468 | 6 | 2 | Mixed-low | 68 |

**Table S1-2 *(continued)***: *This section shows the results for the eleventh part of Table S1-2.*

| Block | Plot | Plant | Plot position | Plot-level chemotype richness | Chemotype | Daughter |
|---|---|---|---|---|---|---|
| 6 | 79 | 469 | 1 | 3 | Bthu-high | 14 |
| 6 | 79 | 470 | 2 | 3 | Mixed-high | 30 |
| 6 | 79 | 471 | 3 | 3 | Mixed-high | 21 |
| 6 | 79 | 472 | 4 | 3 | Chrys-acet | 100 |
| 6 | 79 | 473 | 5 | 3 | Bthu-high | 11 |
| 6 | 79 | 474 | 6 | 3 | Chrys-acet | 94 |
| 6 | 80 | 475 | 1 | 6 | Bthu-low | 103 |
| 6 | 80 | 476 | 2 | 6 | Mixed-low | 64 |
| 6 | 80 | 477 | 3 | 6 | Chrys-acet | 100 |
| 6 | 80 | 478 | 4 | 6 | Mixed-high | 30 |
| 6 | 80 | 479 | 5 | 6 | Athu-Bthu | 57 |
| 6 | 80 | 480 | 6 | 6 | Bthu-high | 17 |
| 6 | 81 | 481 | 1 | 3 | Athu-Bthu | 55 |
| 6 | 81 | 482 | 2 | 3 | Chrys-acet | 95 |
| 6 | 81 | 483 | 3 | 3 | Bthu-low | 101 |
| 6 | 81 | 484 | 4 | 3 | Bthu-low | 103 |
| 6 | 81 | 485 | 5 | 3 | Chrys-acet | 94 |
| 6 | 81 | 486 | 6 | 3 | Athu-Bthu | 56 |
| 6 | 82 | 487 | 1 | 1 | Mixed-high | 23 |
| 6 | 82 | 488 | 2 | 1 | Mixed-high | 21 |
| 6 | 82 | 489 | 3 | 1 | Mixed-high | 30 |
| 6 | 82 | 490 | 4 | 1 | Mixed-high | 23 |
| 6 | 82 | 491 | 5 | 1 | Mixed-high | 30 |
| 6 | 82 | 492 | 6 | 1 | Mixed-high | 21 |
| 6 | 83 | 493 | 1 | 3 | Mixed-low | 67 |
| 6 | 83 | 494 | 2 | 3 | Athu-Bthu | 57 |
| 6 | 83 | 495 | 3 | 3 | Mixed-low | 68 |
| 6 | 83 | 496 | 4 | 3 | Bthu-high | 17 |
| 6 | 83 | 497 | 5 | 3 | Athu-Bthu | 55 |
| 6 | 83 | 498 | 6 | 3 | Bthu-high | 11 |
| 6 | 84 | 499 | 1 | 2 | Chrys-acet | 94 |
| 6 | 84 | 500 | 2 | 2 | Chrys-acet | 100 |
| 6 | 84 | 501 | 3 | 2 | Bthu-high | 14 |
| 6 | 84 | 502 | 4 | 2 | Chrys-acet | 95 |
| 6 | 84 | 503 | 5 | 2 | Bthu-high | 11 |
| 6 | 84 | 504 | 6 | 2 | Bthu-high | 17 |

*1.2.3. Stem phenology*

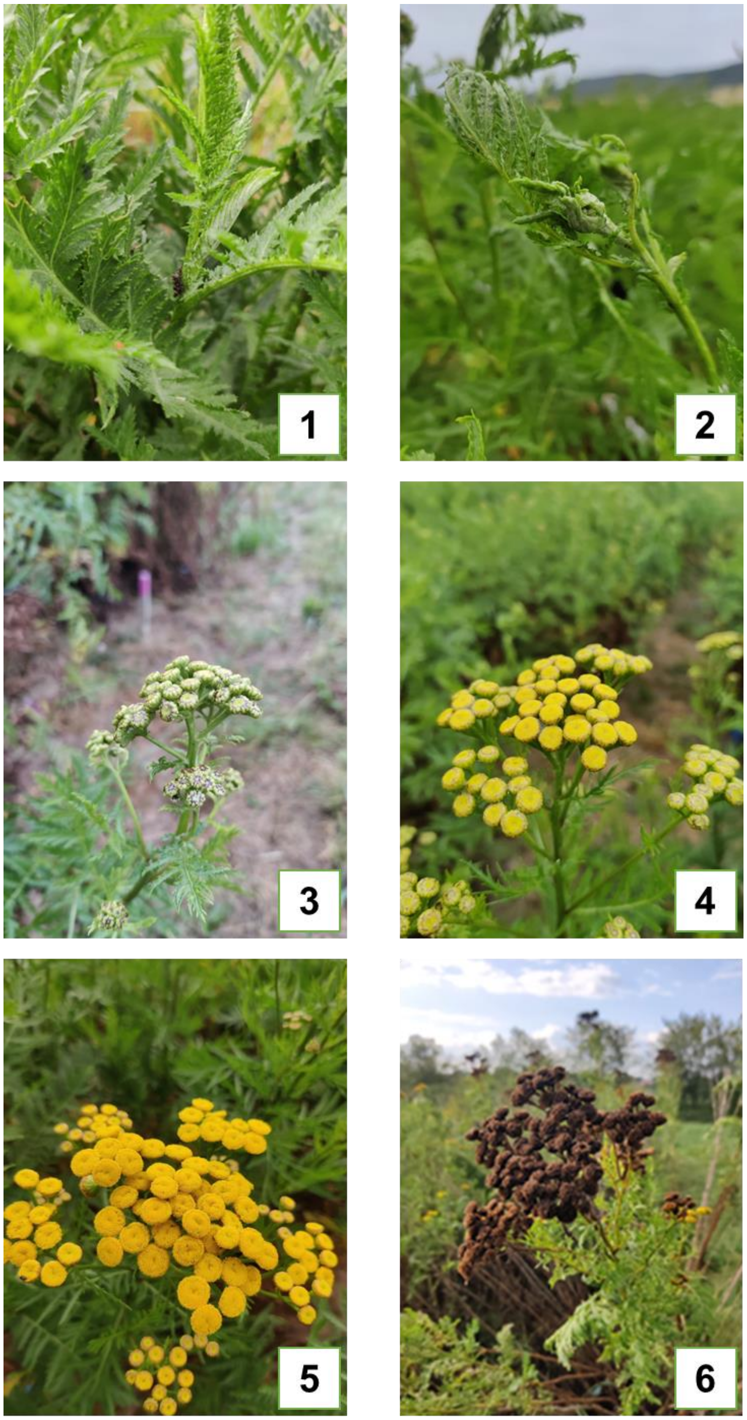


**Figure S1-1:** Stem phenology (*left-to-right then top-to-bottom*): **[1]** No inflorescence; **[2]** Inflorescence visible but closed; **[3]** Open inflorescence; **[4]** Beginning seed set (unripe); **[5]** Ripe seeds; **[6]** Mature seeds turning brown.

### 1.3. Details of the statistical analyses

#### *1.3.1. Variables used in the analyses*

| Variable name | Description | Variable type | Factor levels |
|---|---|---|---|
| B | Experimental block | Factor | 1 to 6 |
| P | Experimental plot | Factor | 1 to 84 |
| CR | Plot-level chemotype richness | Integer | 1, 2, 3, 6 |
| C | Chemotype identity | Factor | Athu-Bthu, Bthu-high, Bthu-low, Chrys-acet, Mixed-high, Mixed-low |
| D | Daughter identity | Factor | Daughter ID of a particular chemotype |
| D: Athu-Bthu | Daughter identity | Factor | 55, 56, 57 |
| D: Bthu-high | Daughter identity | Factor | 11, 14, 17 |
| D: Bthu-low | Daughter identity | Factor | 101, 103, 104 |
| D: Chrys-acet | Daughter identity | Factor | 94, 95, 100 |
| D: Mixed-high | Daughter identity | Factor | 21, 23, 30 |
| D: Mixed-low | Daughter identity | Factor | 64, 67, 68 |
| CP | Chemotype presence | Factor, binary | Presence/absence (1/0) of a particular chemotype |
| CP: Athu-Bthu | Particular chemotype | Factor, binary | Athu-Bthu presence/absence (1/0) at plot level |
| CP: Bthu-high | Particular chemotype | Factor, binary | Bthu-high presence/absence (1/0) at plot level |
| CP: Bthu-low | Particular chemotype | Factor, binary | Bthu-low presence/absence (1/0) at plot level |
| CP: Chrys-acet | Particular chemotype | Factor, binary | Chrys-acet presence/absence (1/0) at plot level |
| CP: Mixed-high | Particular chemotype | Factor, binary | Mixed-high presence/absence (1/0) at plot level |
| CP: Mixed-low | Particular chemotype | Factor, binary | Mixed-low presence/absence (1/0) at plot level |
| Day | Day of headspace VOC collection | Factor | 1 to 3 |

*1.3.2. Models of effects on plant traits*

For each analysis, we give the model run in R.

**Plant-level traits**

*1.3.2.1. Effects of chemotype, plot-level chemotype richness, and the interaction between chemotype and plot-level chemotype richness on plant-level traits*

We first analyzed the effect of chemotype identity and plot-level chemotype richness on the performance of individual plants within the plots.

*Growth traits*

*Number of stems:* We analyzed four measurements of the number of stems per plant separately by time point. Here, we analyzed the number of stems of the individual plants within the stands differing in chemotype richness. We evaluated the effects of chemotype identity (C), plot-level chemotype richness (CR), and their interaction (C x CR). Random effects included were Plot nested in Block (B/P) and Daughter (D). Post hoc tests were only calculated if differences within a factor in the analysis of variance were significant.

| **Number of stems** | **Model** | **Analysis of variance** | **Post hoc test** |
|---|---|---|---|
| June 01, 2023 | glmer(Stems_01.06.2021 ~ C * CR + (1\|B/P) + (1\|D), family = poisson, control = glmerControl(optimizer ="Nelder_Mead") | Anova() | emmeans(model, list(pairwise ~ C), adjust = "tukey") |
| June 22, 2021 | glmer(Stems_22.06.2021 ~ C * CR + (1\|B/P) + (1\|D), family = poisson, control = glmerControl(optimizer ="Nelder_Mead")) | Anova() | emmeans(model, list(pairwise ~ C), adjust = "tukey") |
| October 28, 2021 | glmer(Stems_28.10.2021 ~ C * CR + (1\|B/P) + (1\|D), family = poisson) | Anova() | - |
| July 06, 2022 | glmer(Stems_06.07.2022 ~ C * CR + (1\|B/P) + (1\|D), family = poisson) | Anova() | - |

*Plant height:* We analyzed six measurements of the height per plant separately by time point. Here, we analyzed the number of stems of plants' height (cm) within the stands differing in chemotype richness. We evaluated the effects of chemotype identity (C), plot-level chemotype richness (CR), and their interaction (C x CR). Random effects included were Plot nested in Block

(B/P) and Daughter (D). Post hoc tests were only calculated if differences within a factor in the analysis of variance were significant.

| Height (cm) | Model | Analysis of variance | Post hoc test |
|---|---|---|---|
| June 01, 2023 | lmer(Height_01.06.2021 ~ C x CR + (1\|B/P) + (1\|D)) | Anova() | - |
| June 22, 2021 | lmer(Height_22.06.2021 ~ C x CR + (1\|B/P) + (1\|D)) | Anova() | - |
| October 28, 2021 | lmer(Height_28.10.2021 ~ C x CR + (1\|B/P) + (1\|D)) | Anova() | emmeans(model, list(pairwise ~ Chemotype), adjust = "tukey") |
| May 14, 2022 | lmer(Height_14.05.2022 ~ C x CR + (1\|B/P) + (1\|D)) | Anova() | - |
| July 06, 2022 | lmer(Height_06.07.2022 ~ C x CR + (1\|B/P) + (1\|D)) | Anova() | - |
| October 05, 2022 | lmer(Height_05.10.2022 ~ C x CR + (1\|B/P) + (1\|D)) | Anova() | - |

*Above-ground dry weight:* There was one measurement of the above-ground dry weight per plant on October 28, 2021. Here, we analyzed plants' weight (g) within the stands differing in chemotype richness. We evaluated the effects of chemotype identity (C), plot-level chemotype richness (CR), and their interaction (C x CR). Random effects included were Plot nested in Block (B/P), and Daughter (D). Post hoc tests were only calculated if differences within a factor in the analysis of variance were significant.

| Above-ground dry weight (g) | Model | Analysis of variance | Post hoc test |
|---|---|---|---|
| October 28, 2021 | lmer(sqrt(Dry_biomass_28.10.2021) ~ C * CR + (1\|B/P) + (1\|D) | Anova() | - |

*Above-ground fresh weight:* There was one measurement of the above-ground fresh weight per plant on October 05, 2022. Here, we analyzed plants' weight (g) within the stands differing in chemotype richness. We evaluated the effects of chemotype identity (C), plot-level chemotype richness (CR), and their interaction (C x CR). Random effects included were Plot nested in Block (B/P), and Daughter (D). Post hoc tests were only calculated if differences within a factor in the analysis of variance were significant.

| Above-ground fresh weight (g) | Model | Analysis of variance | Post hoc test |
|---|---|---|---|

| | | | |
|---|---|---|---|
| October 05, 2022 | lmer(sqrt(Fresh_biomass_05.10.2022) ~ C * CR + (1\|B/P) + (1\|D) | Anova() | - |

*Reproductive traits*

*Cumulative number of flower heads:* From June 01 to October 28, 2021, mature flower heads were counted and collected from individual plants. Here, we analyzed the total number of flower heads produced by each plant within the stands differing in chemotype richness during that period. We evaluated the effects of chemotype identity (C), plot-level chemotype richness (CR), and their interaction (C x CR). Random effects included were Plot nested in Block (B/P) and Daughter (D). Post hoc tests were only calculated if differences within a factor in the analysis of variance were significant.

| **Cumulative number of flower heads** | **Model** | **Analysis of variance** | **Post hoc test** |
|---|---|---|---|
| From June 01, 2021, to October 28, 2021 | glmer(Flowerheads ~ C * CR + (1\|B/P) + (1\|D), family = poisson) | Anova() | emmeans(model1, list(pairwise ~ C), adjust = "tukey") |

*Flowering index:* There was one measurement of the flowering index per plant on July 06, 2022. Here, we analyzed the flowering index of plants within the stands differing in chemotype richness. We evaluated the effects of chemotype identity (C), plot-level chemotype richness (CR), and their interaction (C x CR). Random effects included were Plot nested in Block (B/P) and Daughter (D). Post hoc tests were only calculated if differences within a factor in the analysis of variance were significant.

| **Cumulative number of flower heads** | **Model** | **Analysis of variance** | **Post hoc test** |
|---|---|---|---|
| July 06, 2022 | lmer(Floweringindex_06.07.2022 ~ C * CR + (1\|B/P) + (1\|D)) | Anova() | emmeans(model1, list(pairwise ~ C), adjust = "tukey") |

*1.3.2.2. Effects of daughter, plot-level chemotype richness, and the interaction between daughter and plot-level chemotype richness on plant-level traits*

We second analyzed the effect of daughter identity (D), plot-level chemotype richness (CR), and their interaction (D x CR) on four plant traits of individual plants within the plots: Number of stems, height (cm), and above-ground dry weight (g) on October 28, 2021, and cumulative number of flower heads produced in 2021. We carried out a separate analysis per chemotype to test for differences between daughters within a chemotype, and another analysis to directly compare daughters by using all 18 levels of daughter identity (D). We included Plot nested in Block (B/P) as a random factor. Post hoc tests were only calculated if differences within a factor in the analysis of variance were significant.

| Plant trait | Model | Analysis of variance | Post hoc test |
|---|---|---|---|
| **Number of stems: October 28, 2021** | | | |
| Athu-Bthu (D: 55, 56, 57) | glmer(Stems_28.10.2021 ~ D * CR + (1\|B/P), family = poisson) | Anova() | emmeans(model1, list(pairwise ~ D), adjust = "tukey") |
| Bthu-high (D: 11, 14, 17) | glmer(Stems_28.10.2021 ~ D * CR + (1\|B/P), family = poisson) | Anova() | emmeans(model1, list(pairwise ~ D), adjust = "tukey") |
| Bthu-low (D: 101, 103, 104) | glmer(Stems_28.10.2021 ~ D * CR + (1\|B/P), family = poisson) | Anova() | emmeans(model1, list(pairwise ~ D), adjust = "tukey") |
| Chrys-acet (D: 94, 95, 100) | glmer(Stems_28.10.2021 ~ D * CR + (1\|B/P), family = poisson) | Anova() | emmeans(model1, list(pairwise ~ D), adjust = "tukey") |
| Mixed-high (D: 21, 23, 30) | glmer(Stems_28.10.2021 ~ D * CR + (1\|B/P), family = poisson) | Anova() | - |
| Mixed-low (D: 64, 67, 68) | glmer(Stems_28.10.2021 ~ D * CR + (1\|B/P), family = poisson) | Anova() | emmeans(model1, list(pairwise ~ D), adjust = "tukey") |
| Daughter ID (D: 11, 14, 17, 21, 23, 30, 55, 56, 57, 64, 67, 68, 94, 95, 100, 101, 103, 104) | glmer(Stems_28.10.2021 ~ D * CR + (1\|B/P), control = glmerControl(optimizer = "bobyqa", optCtrl = list(maxfun = 2e5)), family = poisson) | Anova() | emmeans(model1, list(pairwise ~ D), adjust = "tukey") |
| **Height (cm): October 28, 2021** | | | |
| Athu-Bthu (D: 55, 56, 57) | lmer(Height_28.10.2021 ~ D * CR + (1\|B/P)) | Anova() | emmeans(model1, list(pairwise ~ D), adjust = "tukey") |
| Bthu-high (D: 11, 14, 17) | lmer(Height_28.10.2021 ~ D * CR + (1\|B/P)) | Anova() | emmeans(model1, list(pairwise ~ D), adjust = "tukey") |
| Bthu-low (D: 101, 103, 104) | lmer(Height_28.10.2021 ~ D * CR + (1\|B/P)) | Anova() | emmeans(model1, list(pairwise ~ D), adjust = "tukey") |
| Chrys-acet (D: 94, 95, 100) | lmer(Height_28.10.2021 ~ D * CR + (1\|B/P)) | Anova() | emmeans(model1, list(pairwise ~ D), adjust = "tukey") |
| Mixed-high (D: 21, 23, 30) | lmer(Height_28.10.2021 ~ D * CR + (1\|B/P)) | Anova() | - |
| Mixed-low (D: 64, 67, 68) | lmer(Height_28.10.2021 ~ D * CR + (1\|B/P)) | Anova() | emmeans(model1, list(pairwise ~ D), adjust = "tukey") |
| Daughter ID | lmer(Height_28.10.2021 ~ D * CR + (1\|B/P)) | Anova() | emmeans(model1, list(pairwise ~ D), adjust = "tukey") |

| | | | |
|---|---|---|---|
| (D: 11, 14, 17, 21, 23, 30, 55, 56, 57, 64, 67, 68, 94, 95, 100, 101, 103, 104) | | | |
| **Above-ground dry biomass (g): October 28, 2021** | | | |
| Athu-Bthu (D: 55, 56, 57) | lmer(sqrt(Dry_biomass_28.10.2021 ~ D * CR + (1\|B/P)) | Anova() | - |
| Bthu-high (D: 11, 14, 17) | lmer(sqrt(Dry_biomass_28.10.2021 ~ D * CR + (1\|B/P)) | Anova() | emmeans(model1, list(pairwise ~ D), adjust = "tukey") |
| Bthu-low (D: 101, 103, 104) | lmer(sqrt(Dry_biomass_28.10.2021 ~ D * CR + (1\|B/P)) | Anova() | emmeans(model1, list(pairwise ~ D), adjust = "tukey") |
| Chrys-acet (D: 94, 95, 100) | lmer(sqrt(Dry_biomass_28.10.2021 ~ D * CR + (1\|B/P)) | Anova() | emmeans(model1, list(pairwise ~ D), adjust = "tukey") |
| Mixed-high (D: 21, 23, 30) | lmer(sqrt(Dry_biomass_28.10.2021 ~ D * CR + (1\|B/P)) | Anova() | emmeans(model1, list(pairwise ~ D), adjust = "tukey") |
| Mixed-low (D: 64, 67, 68) | lmer(sqrt(Dry_biomass_28.10.2021 ~ D * CR + (1\|B/P)) | Anova() | emmeans(model1, list(pairwise ~ D), adjust = "tukey") |
| Daughter ID (D: 11, 14, 17, 21, 23, 30, 55, 56, 57, 64, 67, 68, 94, 95, 100, 101, 103, 104) | lmer(sqrt(Dry_biomass_28.10.2021 ~ D * CR + (1\|B/P)) | Anova() | emmeans(model1, list(pairwise ~ D), adjust = "tukey") |
| **Cumulative number of flower heads: From June 01, 2021, to October 28, 2021** | | | |
| Athu-Bthu (D: 55, 56, 57) | lmer(Flowerheads ~ D * CR + (1\|B/P)) | Anova() | - |
| Bthu-high (D: 11, 14, 17) | lmer(Flowerheads ~ D * CR + (1\|B/P)) | Anova() | emmeans(model1, list(pairwise ~ D), adjust = "tukey") |
| Bthu-low (D: 101, 103, 104) | lmer(Flowerheads ~ D * CR + (1\|B/P)) | Anova() | emmeans(model1, list(pairwise ~ D), adjust = "tukey") |
| Chrys-acet (D: 94, 95, 100) | lmer(Flowerheads ~ D * CR + (1\|B/P)) | Anova() | - |
| Mixed-high (D: 21, 23, 30) | lmer(Flowerheads ~ D * CR + (1\|B/P)) | Anova() | emmeans(model1, list(pairwise ~ D), adjust = "tukey") |
| Mixed-low (D: 64, 67, 68) | lmer(Flowerheads ~ D * CR + (1\|B/P)) | Anova() | emmeans(model1, list(pairwise ~ D), adjust = "tukey") |
| Daughter ID (D: 11, 14, 17, 21, 23, 30, 55, 56, 57, 64, 67, 68, 94, 95, 100, 101, 103, 104) | lmer(Flowerheads ~ D * CR + (1\|B/P)) | Anova() | emmeans(model1, list(pairwise ~ D), adjust = "tukey") |

**Plot-level traits**

*1.3.2.3. Effects of chemotype presence, plot-level chemotype richness, and the interaction between chemotype presence and plot-level chemotype richness on plot-level traits*

We first averaged each plant trait at the plot level to test the effects of chemotype and chemotype richness on plot-level plant traits (Community-Weighted Means -CWM-). To test the effect of the presence/absence of a chemotype on plot-level trait values, we carried out a separate analysis per chemotype. This was done by running a linear model with Block (B), chemotype presence (CP- indicating whether a specific chemotype is present in the plot), plot-level chemotype richness (CR), and their interaction (CP x CR) as fixed effects, separately for each chemotype.

*Growth traits*

*Number of stems:* We analyzed four measurements of the number of stems per plot separately by time point. Here, we analyzed the average number of stems of plots of different chemotype richness. We evaluated the effects of block (B), chemotype presence (CP), plot-level chemotype richness (CR), and their interaction (CP x CR). Post hoc tests were only calculated if differences within a factor in the analysis of variance were significant.

| **Plot-level number of stems** | **Model** | **Analysis of variance** |
|---|---|---|
| **June 01, 2021** | | |
| Athu-Bthu (CP: 1/0) | lm(Stems_01.06.2021 ~ B + CR * CP) | anova() |
| Bthu-high (CP: 1/0) | lm(Stems_01.06.2021 ~ B + CR * CP) | anova() |
| Bthu-low (CP: 1/0) | lm(Stems_01.06.2021 ~ B + CR * CP) | anova() |
| Chrys-acet (CP: 1/0) | lm(Stems_01.06.2021 ~ B + CR * CP) | anova() |
| Mixed-high (CP: 1/0) | lm(Stems_01.06.2021 ~ B + CR * CP) | anova() |
| Mixed-low (CP: 1/0) | lm(Stems_01.06.2021 ~ B + CR * CP) | anova() |
| **June 22, 2021** | | |
| Athu-Bthu (CP: 1/0) | lm(Stems_22.06.2021 ~ B + CR * CP) | anova() |

| | | |
|---|---|---|
| Bthu-high (CP: 1/0) | lm(Stems_22.06.2021 ~ B + CR * CP) | anova() |
| Bthu-low (CP: 1/0) | lm(Stems_22.06.2021 ~ B + CR * CP) | anova() |
| Chrys-acet (CP: 1/0) | lm(Stems_22.06.2021 ~ B + CR * CP) | anova() |
| Mixed-high (CP: 1/0) | lm(Stems_22.06.2021 ~ B + CR * CP) | anova() |
| Mixed-low (CP: 1/0) | lm(Stems_22.06.2021 ~ B + CR * CP) | anova() |
| **October 28, 2022** | | |
| Athu-Bthu (CP: 1/0) | lm(Stems_28.10.2021 ~ B + CR * CP) | anova() |
| Bthu-high (CP: 1/0) | lm(Stems_28.10.2021 ~ B + CR * CP) | anova() |
| Bthu-low (CP: 1/0) | lm(Stems_28.10.2021 ~ B + CR * CP) | anova() |
| Chrys-acet (CP: 1/0) | lm(Stems_28.10.2021 ~ B + CR * CP) | anova() |
| Mixed-high (CP: 1/0) | lm(Stems_28.10.2021 ~ B + CR * CP) | anova() |
| Mixed-low (CP: 1/0) | lm(Stems_28.10.2021 ~ B + CR * CP) | anova() |
| **July 06, 2022** | | |
| Athu-Bthu (CP: 1/0) | lm(Stems_06.07.2022 ~ B + CR * CP) | anova() |
| Bthu-high (CP: 1/0) | lm(Stems_06.07.2022 ~ B + CR * CP) | anova() |
| Bthu-low (CP: 1/0) | lm(Stems_06.07.2022 ~ B + CR * CP) | anova() |
| Chrys-acet (CP: 1/0) | lm(Stems_06.07.2022 ~ B + CR * CP) | anova() |
| Mixed-high (CP: 1/0) | lm(Stems_06.07.2022 ~ B + CR * CP) | anova() |
| Mixed-low (CP: 1/0) | lm(Stems_06.07.2022 ~ B + CR * CP) | anova() |

*Plant height:* We analyzed six measurements of the height (cm) per plot separately by time point. Here, we analyzed the average height of plots of different chemotype richness. We evaluated the effects of block (B), chemotype presence (CP), plot-level chemotype richness (CR), and their interaction (CP x CR). Post hoc tests were only calculated if differences within a factor in the analysis of variance were significant.

| Plot-level height (cm) | Model | Analysis of variance |
|---|---|---|
| **June 01, 2021** | | |
| Athu-Bthu (CP: 1/0) | lm(Height_01.06.2021 ~ B + CR * CP) | anova() |
| Bthu-high (CP: 1/0) | lm(Height_01.06.2021 ~ B + CR * CP) | anova() |
| Bthu-low (CP: 1/0) | lm(Height_01.06.2021 ~ B + CR * CP) | anova() |
| Chrys-acet (CP: 1/0) | lm(Height_01.06.2021 ~ B + CR * CP) | anova() |
| Mixed-high (CP: 1/0) | lm(Height_01.06.2021 ~ B + CR * CP) | anova() |
| Mixed-low (CP: 1/0) | lm(Height_01.06.2021 ~ B + CR * CP) | anova() |
| **June 22, 2021** | | |
| Athu-Bthu (CP: 1/0) | lm(Height_22.06.2021 ~ B + CR * CP) | anova() |
| Bthu-high (CP: 1/0) | lm(Height_22.06.2021 ~ B + CR * CP) | anova() |
| Bthu-low (CP: 1/0) | lm(Height_22.06.2021 ~ B + CR * CP) | anova() |
| Chrys-acet (CP: 1/0) | lm(Height_22.06.2021 ~ B + CR * CP) | anova() |
| Mixed-high (CP: 1/0) | lm(Height_22.06.2021 ~ B + CR * CP) | anova() |
| Mixed-low (CP: 1/0) | lm(Height_22.06.2021 ~ B + CR * CP) | anova() |
| **October 28, 2022** | | |
| Athu-Bthu (CP: 1/0) | lm(Height_28.10.2021 ~ B + CR * CP) | anova() |
| Bthu-high (CP: 1/0) | lm(Height_28.10.2021 ~ B + CR * CP) | anova() |
| Bthu-low (CP: 1/0) | lm(Height_28.10.2021 ~ B + CR * CP) | anova() |
| Chrys-acet (CP: 1/0) | lm(Height_28.10.2021 ~ B + CR * CP) | anova() |
| Mixed-high | lm(Height_28.10.2021 ~ B + CR * CP) | anova() |

| | | |
|---|---|---|
| (CP: 1/0) | | |
| Mixed-low (CP: 1/0) | lm(Height_28.10.2021 ~ B + CR * CP) | anova() |
| **May 14, 2022** | | |
| Athu-Bthu (CP: 1/0) | lm(Height_14.05.2022 ~ B + CR * CP) | anova() |
| Bthu-high (CP: 1/0) | lm(Height_14.05.2022 ~ B + CR * CP) | anova() |
| Bthu-low (CP: 1/0) | lm(Height_14.05.2022 ~ B + CR * CP) | anova() |
| Chrys-acet (CP: 1/0) | lm(Height_14.05.2022 ~ B + CR * CP) | anova() |
| Mixed-high (CP: 1/0) | lm(Height_14.05.2022 ~ B + CR * CP) | anova() |
| Mixed-low (CP: 1/0) | lm(Height_14.05.2022 ~ B + CR * CP) | anova() |
| **July 06, 2022** | | |
| Athu-Bthu (CP: 1/0) | lm(Height_06.07.2022 ~ B + CR * CP) | anova() |
| Bthu-high (CP: 1/0) | lm(Height_06.07.2022 ~ B + CR * CP) | anova() |
| Bthu-low (CP: 1/0) | lm(Height_06.07.2022 ~ B + CR * CP) | anova() |
| Chrys-acet (CP: 1/0) | lm(Height_06.07.2022 ~ B + CR * CP) | anova() |
| Mixed-high (CP: 1/0) | lm(Height_06.07.2022 ~ B + CR * CP) | anova() |
| Mixed-low (CP: 1/0) | lm(Height_06.07.2022 ~ B + CR * CP) | anova() |
| **October 05, 2022** | | |
| Athu-Bthu (CP: 1/0) | lm(Height_05.10.2022 ~ B + CR * CP) | anova() |
| Bthu-high (CP: 1/0) | lm(Height_05.10.2022 ~ B + CR * CP) | anova() |
| Bthu-low (CP: 1/0) | lm(Height_05.10.2022 ~ B + CR * CP) | anova() |
| Chrys-acet (CP: 1/0) | lm(Height_05.10.2022 ~ B + CR * CP) | anova() |
| Mixed-high (CP: 1/0) | lm(Height_05.10.2022 ~ B + CR * CP) | anova() |

| | | |
|---|---|---|
| Mixed-low (CP: 1/0) | lm(Height_05.10.2022 ~ B + CR * CP) | anova() |

*Above-ground dry weight:* We analyzed one measurement of the above-ground dry weight (g) at plot-level. Here, we analyzed the average weight of plots of different chemotype richness. We evaluated the effects of block (B), chemotype presence (CP), plot-level chemotype richness (CR), and their interaction (CP x CR). Post hoc tests were only calculated if differences within a factor in the analysis of variance were significant.

| **Plot-level above-ground dry weight (g)** | **Model** | **Analysis of variance** |
|---|---|---|
| **October 28, 2021** | | |
| Athu-Bthu (CP: 1/0) | lm(log(Dry_biomass_28.10.2021 ~ B + CR * CP) | anova() |
| Bthu-high (CP: 1/0) | lm(log(Dry_biomass_28.10.2021 ~ B + CR * CP) | anova() |
| Bthu-low (CP: 1/0) | lm(log(Dry_biomass_28.10.2021 ~ B + CR * CP) | anova() |
| Chrys-acet (CP: 1/0) | lm(log(Dry_biomass_28.10.2021 ~ B + CR * CP) | anova() |
| Mixed-high (CP: 1/0) | lm(log(Dry_biomass_28.10.2021 ~ B + CR * CP) | anova() |
| Mixed-low (CP: 1/0) | lm(log(Dry_biomass_28.10.2021 ~ B + CR * CP) | anova() |

*Above-ground fresh weight:* We analyzed one measurement of the above-ground fresh weight (g) at plot-level. Here, we analyzed the average weight of plants within the stands differing in chemotype richness. We evaluated the effects of block (B), chemotype presence (CP), plot-level chemotype richness (CR), and their interaction (CP x CR). Post hoc tests were only calculated if differences within a factor in the analysis of variance were significant.

| **Plot-level above-ground fresh weight (g)** | **Model** | **Analysis of variance** |
|---|---|---|
| **October 05, 2022** | | |
| Athu-Bthu (CP: 1/0) | lm(log(Fresh_biomass_05.10.2022 ~ B + CR * CP) | anova() |
| Bthu-high (CP: 1/0) | lm(log(Fresh_biomass_05.10.2022 ~ B + CR * CP) | anova() |
| Bthu-low (CP: 1/0) | lm(log(Fresh_biomass_05.10.2022 ~ B + CR * CP) | anova() |

| | | |
|---|---|---|
| Chrys-acet (CP: 1/0) | lm(log(Fresh_biomass_05.10.2022 ~ B + CR * CP) | anova() |
| Mixed-high (CP: 1/0) | lm(log(Fresh_biomass_05.10.2022 ~ B + CR * CP) | anova() |
| Mixed-low (CP: 1/0) | lm(log(Fresh_biomass_05.10.2022 ~ B + CR * CP) | anova() |

*Reproductive traits*

*Cumulative number of flower heads:* From June 01 to October 28, 2021, mature flower heads were counted and collected from individual plants. Here, we analyzed the average cumulative number of flower heads of plots of different chemotype richness. We evaluated the effects of block (B), chemotype presence (CP), plot-level chemotype richness (CR), and their interaction (CP x CR). Post hoc tests were only calculated if differences within a factor in the analysis of variance were significant.

| **Plot-level cumulative number of flower heads** | **Model** | **Analysis of variance** |
|---|---|---|
| **June 01 - October 28, 2021** | | |
| Athu-Bthu (CP: 1/0) | lm(log(Flowerheads ~ B + CR * CP) | anova() |
| Bthu-high (CP: 1/0) | lm(log(Flowerheads ~ B + CR * CP) | anova() |
| Bthu-low (CP: 1/0) | lm(log(Flowerheads ~ B + CR * CP) | anova() |
| Chrys-acet (CP: 1/0) | lm(log(Flowerheads ~ B + CR * CP) | anova() |
| Mixed-high (CP: 1/0) | lm(log(Flowerheads ~ B + CR * CP) | anova() |
| Mixed-low (CP: 1/0) | lm(log(Flowerheads ~ B + CR * CP) | anova() |

*Flowering index:* There was one measurement of the flowering index per plant on July 06, 2022. Here, the average flowering index of plots of different chemotype richness. We evaluated the effects of block (B), chemotype presence (CP), plot-level chemotype richness (CR), and their interaction (CP x CR). Post hoc tests were only calculated if differences within a factor in the analysis of variance were significant.

| **Plot-level flowering index** | **Model** | **Analysis of variance** |
|---|---|---|
| **July 06, 2022** | | |
| Athu-Bthu (CP: 1/0) | lm(Floweringindex_06.07.2022 ~ B + CR * CP) | anova() |
| Bthu-high (CP: 1/0) | lm(Floweringindex_06.07.2022 ~ B + CR * CP) | anova() |
| Bthu-low (CP: 1/0) | lm(Floweringindex_06.07.2022 ~ B + CR * CP) | anova() |
| Chrys-acet (CP: 1/0) | lm(Floweringindex_06.07.2022 ~ B + CR * CP) | anova() |
| Mixed-high (CP: 1/0) | lm(Floweringindex_06.07.2022 ~ B + CR * CP) | anova() |
| Mixed-low (CP: 1/0) | lm(Floweringindex_06.07.2022 ~ B + CR * CP) | anova() |

*1.3.3. Models of effects on overyielding indexes*

For each analysis, we give the model run in R.

*1.3.3.1. Effects of plot-level chemotype richness on the overyielding indexes*

We analyzed the effect of plot-level chemotype richness (CR) on the overyielding indexes of four plant traits calculated at plot-level. Random effect included was Block (B).

| **Overyielding index** | **Model** | **Analysis of variance** |
|---|---|---|
| Above-ground dry biomass (g): October 28, 2021 | lmer(Dry_biomass_28.10.2021 ~ CR + (1\|B)) | Anova() |
| Above-ground fresh biomass (g): October 05, 2022 | lmer(Fresh_biomass_05.10.2022 ~ CR + (1\|B)) | Anova() |
| Cumulative number of flower heads: From June 01, 2021, to October 28, 2021 | lmer(Flowerheads ~ CR + (1\|B)) | Anova() |
| Flowering index: July 06, 2022 | lmer(Floweringindex_06.07.2022 ~ CR + (1\|B)) | Anova() |

*1.3.4. Models of chemodiversity*

For each analysis, we give the model run in R.

*1.3.4.1. Effects of plot-level chemotype richness on the plot-level theoretical leaf chemodiversity metrics*

Theoretical plot-level leaf terpenoid diversity metrics were calculated by summing up the absolute leaf terpenoid concentrations (nmol x $g^{-1}$) of each chemotype/daughter present in each plot, based on the chemical analysis of leaves from greenhouse plants in 2020 (Neuhaus-Harr *et al*., 2023). We analyzed the effect of plot-level chemotype richness (CR) on each of the diversity metrics (terpenoid concentration, richness, Shannon diversity and evenness) using a linear model.

| **Diversity metric** | **Model** | **Analysis of variance** |
|---|---|---|
| Squared plot-level theoretical total terpenoid concentration (nmol x $g^{-1}$) | lm((Concentration)^2 ~ CR) | Anova() |
| Plot-level theoretical terpenoid richness | lm(Richness ~ CR) | Anova() |
| Plot-level theoretical terpenoid Shannon diversity | lm(Shannon ~ CR) | Anova() |
| Plot-level theoretical terpenoid evenness | lm(Evenness ~ CR) | Anova() |

*1.3.4.2. Effects of plot-level chemotype richness on the plot-level realized volatile chemodiversity metrics*

Realized plot-level volatile chemodiversity was based on terpenoids collected in the headspace (ng x $h^{-1}$) in May 2022. We analyzed the effect of plot-level chemotype richness (CR) on each of the diversity metrics (terpenoid concentration, richness, Shannon diversity, and evenness) using a linear mixed model that included Block (B) and Day of headspace VOC collection (D) as random factors.

| **Diversity metric** | **Model** | **Analysis of variance** |
|---|---|---|
| Squared root of plot-level realized total volatile terpenoid emission (ng x $h^{-1}$) | lmer(sqrt(Concentration) ~ CR + (1\|B) + (1\|Day)) | Anova() |
| Squared plot-level realized volatile terpenoid richness | lmer((Richness)^2 ~ CR + (1\|B) + (1\|Day)) | Anova() |
| Plot-level realized volatile terpenoid Shannon diversity | lmer(Shannon ~ CR + (1\|B) + (1\|Day)) | Anova() |
| Plot-level theoretical terpenoid evenness | lmer(Evenness ~ CR + (1\|B) + (1\|Day)) | Anova() |
